# Electrical Excitability of the Endoplasmic Reticulum Membrane Drives Electrical Bursting and the Pulsatile Secretion of Insulin in a Pancreatic Beta Cell Model

**DOI:** 10.1101/249805

**Authors:** Javier Gómez-Barriocanal

## Abstract

Pancreatic *β*-cells secrete insulin, the hormone that controls glucose homeostasis in vertebrates. When activated by glucose, *β*-cells display a biphasic electrical response. An initial phase, in which the cell fires action potentials continuously, is followed by a phase with a characteristic firing pattern, known as electrical bursting, that consists on brief pulses of action potentials separated by intervals of rest. Electrical bursting is believed to mediate the pulsatile secretion of insulin. The electrical response of *β*-cells has been extensively studied at experimental and theoretical level. However, there is still no consensus on the cellular mechanisms that underlie each of the phases of the response. In this paper, I propose the hypothesis that the pattern of the plasma membrane (PM) response of stimulated *β*-cells is generated by the electrical activity of the endoplasmic reticulum (ER) membrane. In this hypothesis, the interaction of the two excitable membranes, PM and ER membrane, each operating at a different time scale, generates both, the initial continuous phase and the periodic bursting phase. A mathematical model based on the hypothesis is presented. The behavior of the model *β*-cell replicates the main features of the physiological response of pancreatic *β*-cells to nutrients and to neuro-endocrine regulatory factors. The model cell displays a biphasic response to the simulated elevation of glucose. It generates electrical bursting with frequencies comparable to those observed in live cells. The simulation of the action of regulatory factors mimics the actual effect of the factors on the frequency of bursting. Finally, the model shows that a cell with a defective ER response behaves like a dysfunctional *β*-cell from individuals with type 2 diabetes mellitus, a result that suggests that the electrical malfunction of the ER membrane may represent one of the primary causes of type 2 diabetes. Dynamic analysis of the ER behavior has revealed that, depending on the transport rates of Ca^2+^ in and out of the ER, the system has three possible dynamic states. They consist on the hyperpolarization of the ER membrane, periodic oscillations of the electric potential across the membrane, and the depolarization of the membrane. Each of these states determines a different functional program in the cell. The hyperpolarized state maintains the cell at rest, in a non-secreting state. Periodic oscillations of the ER membrane cause electrical bursting in the PM and the consequent pulsatile secretion of insulin. Finally, the depolarized state causes continuous firing and an acute secretory activity, the hyperactive conditions of the initial phase of the *β*-cell response to glucose. The dynamic states of the ER are also associated with different long-term effects. So, conditions that induce the hyperactive depolarized state in *β*-cells also potentiate apoptosis. The induction of the oscillatory state by glucose and neuro-endocrine factors seems to activate also cell proliferation. In extreme conditions though, such as the chronic treatment of T2DM with incretin analogs, the activation of the oscillatory state may lead to the appearance of cancer. The mathematical model presented here is an illustration of how, even in a extremely simplified system, the nonlinearity or excitability of the ER membrane can produce a repertoire of dynamic states that are able to generate a complex response comparable to the response observed experimentally in pancreatic *β*-cells. In actual cells, with a much higher number of parameters susceptible to be modified by environmental and genetic factors, the ER membrane is likely to have a significantly bigger set of dynamic states each capable to direct the cell in a particular functional or developmental direction. The potential role of the electrical activity of the ER membrane in cellular processes such as fertilization, cell proliferation and differentiation, and cell death, as well as in the development of diverse pathological conditions is analyzed in the discussion.

## 1 Introduction

### 1.1 Regulation of Insulin Secretion

Insulin, the hormone secreted by the *β*-cells in the islets of Langerhans of the pancreas, is involved in the control of glucose homeostasis in higher vertebrates (1). Insulin secretion is regulated by glucose and other nutrients, as well as by hormonal and neural factors (2, 3). A defective regulation results in serious pathological conditions. Unrestrained secretion of insulin causes hyperinsulinism (4, 5), whereas low levels of insulin, due to *β*-cell death or malfunction, are responsible for the development of type 1 and type 2 diabetes (6–8).

The stimulation of pancreatic *β*-cells by glucose elicits a biphasic electrical response (9, 10). A brief initial phase, lasting a few minutes, is characterized by the continuous firing of action potentials. In this phase there is a very active secretion of insulin that causes a stark elevation of the hormone concentration in the bloodstream (11–13). The initial response is followed by a prolonged bursting phase. During this second phase, cells fire rhythmic bursts of action potentials, each lasting 10 to 20 seconds, separated by intervals of electrical inactivity of similar duration (14, 15). The level of insulin secretion is considerably lower than in the initial phase and takes place in pulses that correlate with the electrical bursts (16–20).

In parallel with the electrical oscillations of the plasma membrane (PM), stimulated cells also display oscillations of the cytosolic concentration of calcium ([Ca^2+^]_*CYT*_). This calcium response is also biphasic. A sustained initial transient is followed by periodic oscillations that have the same frequency that the bursting oscillations in the PM (21–23).

Glucose stimulates insulin secretion by means of a synergistic interaction between several signaling pathways (15, 24–26). The best characterized of these pathways involves the metabolism of glucose and K_*ATP*_, an ATP-regulated potassium channel expressed in the plasma membrane of *β*-cells (27). The elevation of blood glucose accelerates the rate of glycolysis in the mitochondria and the production of ATP in *β*-cells. The rise of the ATP/ADP ratio in the cytosol causes the closing of K_*ATP*_ channels and the consequent depolarization of the PM (28–30). When the PM potential (V_*PM*_) reaches the threshold for the activation of voltage-gated calcium channels (Ca_*V*_), the cell starts the firing of action potentials. The influx of Ca^2+^ through the opened Ca_*V*_ channels elevates the cytosolic concentration of calcium and triggers the secretion of insulin (15). This signaling process is known as the glucose triggering pathway (31).

There is evidence that glucose also stimulates the release of insulin by acting directly on the secretory process itself enhancing the response of insulin-containing granules to cytosolic calcium (32–34). The effect of glucose on the late stages of secretion is known as the glucose amplifying pathway (31).

The triggering and amplifying pathways have represented for many years the standard explanation of how glucose induces the release of insulin. However, recent pharmacological and genetic evidence has revealed that this is only part of a more complex picture. The chemical inactivation (32–35), as well as the genetic depletion of K_AT P_ channels (36–38), contrary to what was expected, does not abolish the response of *β*-cell to glucose. So, for instance, mutant mice lacking any of the two K_*ATP*_ subunits, Sur1 or Kir6.2, are nevertheless normoglycemic (39, 40). Interestingly, the genetic inactivation of K_*ATP*_ channels does not even suppress electrical bursting in mutant *β*-cells (41). Additional signaling systems, capable to trigger the glucose-induced secretion of insulin independently of K_*ATP*_, must therefore exist in pancreatic *β*-cells. Two candidates to be involved in this alternative signaling are the inositol and the cAMP pathways.

#### 1.1.1 The Inositol Pathway

The inositol pathway is a signaling system that is usually activated by agonists that bind to G-protein coupled receptors. The binding causes the activation of some form of phospholipase C (PLC), an enzyme that catalyzes the hydrolisis of phosphatidylinositol (4,5)bisphosphate to produce inositol 1,4,5 trisphosphate, IP3, and diacylglycerol (DAG) (42). DAG activates protein kinase C (PKC), whereas inositol trisphosphate binds to IP3 receptors (IP3Rs) in the membrane of the endoplasmic reticulum (ER) and other internal stores (43, 44). IP3Rs are calcium channels gated by Ca^2+^ and IP3 (45, 46). The binding of IP3 in the presence of calcium causes the opening of the channels and the release of calcium from the lumen of the ER into the cytosol. The release has a dual effect, it elevates [Ca^2+^]_*CYT*_ (43) and activates inward currents in the PM known as store operated currents (SOC) or calcium-release activated currents (CRAC) (47) that depolarize the membrane.

In *β*-cells, the inositol pathway is activated by acetylcholine (ACh) and cholinergic agonists such as carbachol that bind to the M3 muscarinic receptor (48, 49). The binding of ACh to the M3 receptor activates PLC-*β* through the action of a heterotrimeric G protein of the G_*q*_ family. The activated PLC raises the cytosolic levels of DAG and IP3. The primary function of DAG in *β*-cells is to activate protein kinase C, although additional roles involving its phosphorilated derivative phosphatidic acid have also been suggested (50). The effects of PKC on insulin secretion are still controversial (51, 52), although they seem to target the late stages of secretion, affecting the amplifying pathway (53).

The IP3-dependent release of calcium from the ER causes [Ca^2+^]_
*CYT*_ oscillations (54) and the activation of an inward Na^+^ current (55, 56). Additionally, the binding of agonists to the M3 receptor activates an IP3-independent inward current probably through a sodium channel, NALCN, associated with the receptor (57, 58).

As a result of all these effects, the activation of the M3 receptor by cholinergic agonists potentiates the glucose-dependent secretion of insulin. The M3 pathway though can not induce by itself the release of insulin in the absence of glucose (59, 60).

The inositol pathway is also directly activated by glucose in *β*-cells, as first reported by Best (61) and Montague (62). These authors observed that the elevation of IP3 required the metabolism of glucose (62, 63) but not the influx of extracellular calcium (62, 64), showing that it was independent of the closing of the K_*ATP*_ channels. These observations have been since thoroughly confirmed and it is well established that the elevation of IP3 is a key component of the K_*ATP*_-independent mechanism by which glucose stimulates the secretion of insulin (26, 65, 66).

#### 1.1.2 The cAMP Pathway

Historically, cAMP was one of the first messengers proposed to mediate the stimulatory action of glucose in *β*-cells (67, 68). Although sometimes questioned, specially after the discovery of the role of the K_*ATP*_ channels, there is now solid evidence that cAMP plays a central role in the induction of insulin secretion by glucose (69–71). Glucose raises the level of cAMP by activating a membrane-bound form of adenylyl cyclase (70) and perhaps also a soluble form of the enzyme (72). The level of cAMP can also be elevated in *β*-cells by incretin hormones such as glucagon-like peptide-1 (GLP-1) and glucose-dependent insulinotropic polypeptide (GIP) (73–75). This pathway has proven to have an important therapeutic value in the treatment of type 2 diabetes (76–80).

cAMP activates protein kinase A (PKA) and a factor known as Epac (exchange protein activated by cAMP). PKA phosphorilates several downstream targets, among them SUR1, the regulatory subunit of the K_*ATP*_ channel. The phosphorilation disrupts the binding of ADP and thus facilitates the closing of the channel and the secretion of insulin (81, 82). Epac is a cAMP-regulated guanine nucleotide exchange protein. This small G-protein activates PLC, causing the elevation of IP3 and opening of IP3Rs (83, 84). It also activates the ryanodine receptor (RyR) type of calcium channels in the ER (85). The combined action of Epac on IP3 and ryanodine receptors produces a significant release of calcium from the internal stores. The importance of this particular effect of cAMP has become evident in genetic experiments in which the inactivation of the genes encoding Epac2 or its downstream effector PLC-*ε* caused a strong reduction of the secretory response of the mutant *β*-cells to both glucose and GLP-1 (86, 87).

Recently, it has been reported that GLP-1 can activate PLC and elevate cytosolic IP3 independently of cAMP by interacting with a G protein of the G_αq_ family (80, 88). These results corroborate the important role that the release of calcium from internal stores plays in promoting the release of insulin.

#### 1.1.3 The NAADP Pathway

A third signaling system associated with the glucose regulation of insulin secretion is the NAADP pathway. Nicotinic acid adenine dinucleotide phosphate (NAADP) is a second messenger involved in the control of multiple cellular processes. NAADP mobilizes calcium from lysosomes and other acidic organelles by activating two calcium channels of the two-pore channel (TPC) family, TPC1 and TPC2 (89, 90). Glucose, as well as GLP-1, stimulates the formation of NAADP in *β*-cells (91, 92). NAADP elevates [Ca^2+^]_*CYT*_ and induces periodic depolarizing currents in the plasma membrane (93, 94). The deletion of TPC1 (94), although apparently not that of TPC2 (95), reduces the response of the mutant cells to glucose. NAADP is therefore part of the K_*ATP*_-independent mechanism involved in the stimulation of insulin secretion by glucose.

#### 1.1.4 Store Operated Currents

A common result in all three K_*ATP*_-independent pathways described above is the activation of inward currents in the PM by the release of calcium from internal stores. This type of store operated currents have been identified in excitable and non-excitable cells and play important roles in a wide variety of physiological processes (96, 97).

A calcium current, activated by the depletion of calcium from the ER, was postulated by Putney in 1986 (98). The actual current was first observed by Hoth et al. in 1992 (99). The calcium channel responsible for the current, Orai1, was identified and characterized in 2006 (100). At about the same time it was identified STIM1 (101, 102), an ER membrane protein that is believed to sense the lumenal Ca^2+^ concentration. When [Ca^2+^]_*ER*_ is low, STIM1 translocates to the proximity of the PM, interacts with Orai1 and activates the channel causing the entrance of extracellular Ca^2+^ into the cytosol and the depolarization of the PM (103). The Orai1-dependent current was initially believed to be a mechanism to replenish the ER of Ca^2+^, but it is well established now that the current is involved in more complex functions (47).

Whereas the STIM1/Orai1-dependent SOC is selective for calcium, a second type of store operated current is mediated by a family of non-selective cation channels (104). The prototype of the family is TRP (transient receptor potential), a channel discovered in the photoreceptors of Drosophila melanogaster (105). TRP is involved in the transduction of the light signal into neural impulses. Light activates PLC, elevates IP3 and causes the release of calcium from the ER. As a result, TRP becomes open and conducts a current that depolarizes the plasma membrane and triggers the firing of action potentials by the photoreceptor (106).

Both types of channels, Orai1 and TRP are expressed in *β*-cells and participate in the regulation of insulin secretion (107, 108). Orai1 forms a ternary complex with STIM1 and TRPC1, a member of the TRP family. The complex is required for the normal induction of insulin secretion by glucose and cholinergic agonists (109). Another member of the TRP family that has been shown to be involved in the regulation of insulin secretion is TRPM5. TRPM5 was first identified in taste cells and found to cause the depolarization of the PM in response to the IP3-induced Ca^2+^ release from the ER (110–112). The channel is expressed in *β*-cells (113, 114) and its inactivation reduces the response of the cells to glucose and GLP-1 (114–117). Mice carrying a Trpm5 −/− mutation display elevated levels of blood glucose and a type 2 diabetes-like phenotype (114, 115). Interestingly, the inactivation also abolishes the normal bursting response of *β*-cells to glucose (114). In humans, genetic variants within the TRPM5 locus have been found associated with an increased risk of developing type 2 diabetes (118). These results show that the store operated currents play a key role in the induction of insulin secretion in *β*-cells.

In summary, glucose stimulates the secretion of insulin by acting on two separate targets, the plasma membrane and the internal stores. On the PM, glucose closes K_*ATP*_ channels and causes the depolarization of the membrane. At the level of the internal stores, glucose activates the release of Ca^2+^ into the cytosol through the IP3R, RyR, TPC1 and perhaps other calcium channels. The release elevates [Ca^2+^]_*CYT*_ and activates store operated currents that contribute to the depolarization of the PM. The synergistic interaction of the two levels of signaling causes the secretion of insulin. The interaction of the two pathways may also represent the mechanism by which *β*-cells generate electrical bursting.

### 1.2 Mathematical Models of Electrical Bursting

As described in the previous section, the activation of *β*-cells produces electrical bursting, brief episodes of electrical activity in which the PM becomes depolarized and the cells fire action potentials, separated by short intervals of electrical rest caused by the repolarization of the PM (15, 119). The electrical oscillations of the PM are associated with parallel oscillations of the calcium concentration in the cytoplasm (21, 22, 120). Bursting has been observed in isolated islets (9, 10) as well as in recordings made in whole animals (121, 122) and seems critical for the regulatory function of insulin (123, 124). The loss of bursting has been associated with the development of hyperglycemia and T2DM (20, 125–127).

Electrical bursting has been observed in many other excitable cells such as the R15 neuron of Aplysia (128), the pituitary gonadotropes (129, 130), neurons in the mammalian neocortex (131) or the dopamine midbrain neurons involved in the development of Parkinson’s disease (132, 133). The exact role of bursting in these excitable cells is still not known.

Since its discovery in the late sixties (9, 10), electrical bursting in *β*-cells has been the object of intensive investigation at both experimental and theoretical levels (134–137). In spite of this effort, the cellular mechanisms underlying the electrical oscillations of the PM that cause bursting still remain poorly understood. In the words of Berggren and Barker (138), the repolarization of the beta-cell after a bursting event is the least well understood process in the *β*-cell stimulus-secretion coupling.

#### 1.2.1 Cytosolic Calcium Hypotheses

Several mechanisms have been postulated to try to explain electrical bursting in terms of the oscillations of the cytosolic concentration of calcium. One of the earliest was proposed by Atwater and colleagues (139, 140). This group suggested that bursting could be driven by a feedback mechanism dependent on [Ca^2+^]_*CYT*_ oscillations. According to these authors, Ca^2+^ would regulate the opening state of a calcium-activated potassium channel (K_*Ca*_) and in this way generate the oscillations of PM potential that initiate and end each bursting cycle. In the absence of glucose, [Ca^2+^]_*CYT*_ would be elevated, keeping the K_*Ca*_ channels open and the PM hyperpolarized. The addition of glucose would activate Ca^2+^ pumps that extrude calcium from the cytosol and thus reduce [Ca^2+^]_*CYT*_. At sufficiently low [Ca^2+^]_*CYT*_, the K_*Ca*_ channels would become closed causing the depolarization of the PM. When the PM potential reaches the threshold for the activation of voltage-gated calcium channels (Ca*_V_*), the cell would start firing action potentials. The entrance of calcium through the opened Ca_V_ channels would cause a gradual elevation of [Ca^2+^]_*CYT*_ that eventually would reach the level for the reactivation of the K_*Ca*_ channels. The opening of the K_*Ca*_ channels would cause the repolarization of the PM and restore the electrically inactive initial conditions. Then the whole cycle would be repeated again (140).

Interestingly, in their experimental studies the authors measured the electrical resistance of the PM during bursting and found a decrease of the resistance during the active phase, a result more consistent with the active phase being caused by the opening of some channel or channels rather than the closing of a calcium-regulated K_*Ca*_ channel as they proposed (139).

The Atwater’s hypothesis was implemented in a mathematical model by Chay and Keizer in 1983 (141). The model assumed a Hodgkin and Huxley formalism (142) to describe the ionic currents across the PM. The rate of change of the PM potential is thus given by the equation:

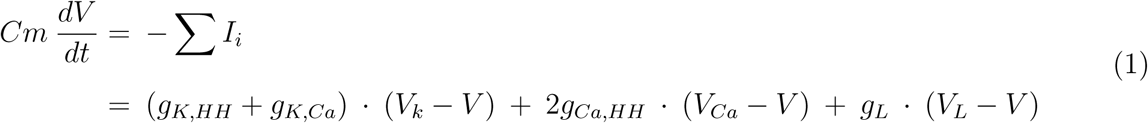

where *Cm* and *V* are respectively the capacitance and the electric potential of the plasma membrane, and *g*_*i*_ represents the conductance of each of the channels in the model. These include a voltage-dependent Ca channel, *Ca*_*HH*_, that played the role of the voltage-dependent sodium channel in the Hodgkin and Huxley model. The conductance of this channel, *g*_*Ca,HH*_, is a function of the time-dependent variables m and h, the activating and inactivating parameters of the channel, as in the Hodgkin and Huxley scheme:

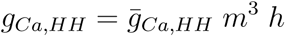

In addition, the model included two potassium channels regulated by voltage, *K*_*HH*_, and calcium, *K*_*Ca*_, and a leak channel. Finally, *V*_*i*_ represents the reversal potential for each of the ions involved. An additional equation for the balance of calcium completed the model.

Simulations using this model showed that the mechanism proposed by Atwater’s group could indeed generate electrical bursting comparable to that observed in *β*-cells (141).

The Chay and Keizer model represented an important breakthrough in the theoretical analysis of electrical bursting and has exerted a considerable influence in the subsequent development of the field. In particular, the use of the Hodgkin and Huxley formalism as shown in Eq. 1 has been adopted, with only small variations, in practically all mathematical models developed afterwards.

The actual measurement of [Ca^2+^]_*CYT*_ in pancreatic cells (21, 22) revealed the limitations of the Atwater’s hypothesis. The dynamics of calcium in *β*-cells is significantly different from that assumed in the model. The [Ca^2+^]_*CYT*_ is low in cells at rest, rises sharply at the beginning of a bursting cycle and remains elevated during the whole active phase. There is no gradual increase of the concentration leading to a critical [Ca^2+^]_*CYT*_ at the end of the active phase that terminates the firing of action potentials as proposed by Atwater.

This square wave type of dynamics poses a problem to any bursting model based on a PM channel regulated by calcium. The pacemaker channel must respond slowly to changes of [Ca^2+^]_*CYT*_, with a delay comparable to the length of the active phase. In a mathematical model this can be easily achieved just by using a Hodgkin and Huxley type of gating variable such as *m* or *h* of Eq. 1. The rate of change of these variables has the general expression:

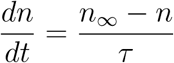

where *n*_∞_ represents the value of the variable when gating is instantaneous, usually a Michaelis-Menten function of [Ca^2+^]_*CYT*_, and *τ* is an arbitrary relaxation time parameter. By selecting the appropriate value of tau, it is possible to adjust the delay response of any channel to the level required by the experimental observations.

This strategy has been used in many models. A slowly inactivating voltage-dependent calcium channel, Ca_*V*_, was postulated in *β*-cell models developed by Teresa Chay (143) and Keizer and Smolen (144). Other groups have maintained the K_*Ca*_ channel proposed by Atwater’s group as the bursting pacemaker but assuming a slow gating mechanism (136, 145, 146).

The physiological relevance of these hypotheses depends on the experimental identification of the postulated slow gating channel. In 1999, Gopel et al. (145) reported some indirect evidence of a slow current that could be mediated by a slow K_*Ca*_ channel involved in bursting. However, inhibitors that block this slow K^+^ current do not prevent electrical bursting in glucose stimulated cells (147). Using other specific inhibitors, it has also been possible to discarded the involvement of the main known K_*Ca*_ channels (148), the large conductance, BK (149, 150), the intermediate conductance, IK or SK4 (149, 151), and the small conductance SK1-3 (152, 153), in the generation of electrical bursting in *β*-cells.

Currently, no slow gating K_*Ca*_ channel able to couple [Ca^2+^]_*CYT*_ with the electrical oscillations of the PM in *β*-cells has been identified.

As an alternative to the regulation of PM potential through the gating of a slow K_*Ca*_ channel, several groups have proposed that bursting can be generated by oscillations of ATP caused by fluctuations of the glycolitic process in the mitochondria of *β*-cells. The oscillations of the ATP/ADP ratio would then affect the conductance of the K_*ATP*_ channels and generate the oscillations of the PM potential. The first model using this approach was developed by Keizer and Magnus in 1989 (154). For these authors, the metabolic oscillations were caused by the oscillations of [Ca^2+^]_*CYT*_ and had therefore the same frequency (155, 156). In other models the glycolitic oscillations are autonomous and thus are not synchronized with the oscillations of cytosolic calcium (157–159). A more complex model, that involves both slow K_*Ca*_ channels and the effect of metabolic oscillations on K_*ATP*_ has been developed by Bertram, Sherman and Satin (160, 161). This model, referred to as the double oscillator model, can reproduce most of the observed electrical responses of *β*-cells, including electrical bursting.

There is experimental evidence that ATP and K_*ATP*_ conductance oscillate in glucose stimulated *β*-cells (162–164). However, these oscillations are considerably slower than bursting and they have been observed in non-stimulated cells where there is no bursting and in over-stimulated cells where spike firing is continuous (164). More importantly, the genetic inactivation of K_*ATP*_ channels in SUR1 −/− and Kir6.2 −/− *β*-cells does not abolish bursting (38, 41). Yildirim et al. (165) have suggested that in theses mutant cells, Kir2.1, a voltage-dependent inward rectifier potassium channel, could compensate for the lack of K_*ATP*_ channels.

#### 1.2.2 ER Calcium Hypotheses

The discovery of currents activated by the depletion of calcium from internal stores, in particular the finding that the release of calcium from the ER caused the activation of an inward current in the PM of *β*-cells (166, 167), opened the possibility to explain electrical bursting using a different strategy. In the new type of models, the active phase of bursting will be initiated by the opening of store operated channels and the activation of an inward depolarizing current. The active phase would then terminate when the channels become closed.

In 1996, Teresa Chay, aware of the limitations of her previous models based on the oscillations of cytosolic calcium (143, 168) started to explore the possibility that the slow variable driving electrical bursting was the lumenal concentration of calcium ([Ca^2+^]_*ER*_) rather than the concentration of calcium in the cytosol ([Ca^2+^]_*CYT*_). She developed a mathematical model implementing this hypothesis (169, 170). In the model, she postulated that the alternative Ca^2+^ depletion and refilling of the ER would cause the activation and termination of a store operated current responsible for bursting. In resting cells, the ER would have an elevated concentration of calcium. The store operated current would be inactive and the PM hyperpolarized. Glucose would cause the Ca^2+^ depletion of the ER, activate a store operated current and depolarize the PM. This would trigger the firing of action potentials initiating an active bursting phase. The influx of calcium through the voltage-activated Ca^2+^ channels would cause the refill of the ER, leading to the inactivation of the store operated current, the repolarization of the PM and the termination of the active phase of bursting. During the subsequent silent phase, glucose would again cause the depletion of the ER until a low [Ca^2+^]_*ER*_ initiates a new bursting cycle. Simulations using the equations of the model were able to reproduce the observed electrical bursting (169, 170).

The main objection raised to this model was that the elevation of glucose does not deplete the ER of calcium but rather stimulates the transport of the ion into the organelle, probably by activating the ATP-dependent SERCA pump (23, 171). A mechanism based on the concentration of calcium in the lumen of the ER was therefore unlikely to be the pacemaker of electrical bursting in *β*-cells.

The store operated currents have been incorporated in other models of electrical activity of *β*-cells. Bertram and colleagues were actually the first to include SO currents in a mathematical model to explain the depolarizing effect of ACh (172), although in this model bursting is generated not by the store-regulated current but by a slow inactivating K_*Ca*_ channel in the plasma membrane. Roe et al. also proposed the involvement of SO currents in bursting (173) and Fridlyand et al. incorporated the concept in a mathematical model (174).

Although the concentration of calcium in the lumen of the ER seems unlikely to represent the slow variable responsible for bursting, there is compelling evidence that the store operated currents play nevertheless a fundamental role in the generation of the slow oscillations of PM potential seen during bursting. The molecular characterization of some of the store operated channels in *β*-cells has allowed to prove their direct involvement in the stimulus-secretion process. In particular, the observation that the genetic inactivation of TRPM5 (114) has a deleterious effect on both bursting and insulin secretion shows that the store operated channels, and therefore the control of PM currents by the ER, represent a fundamental part of the mechanism that mediates electrical bursting and the secretory response of *β*-cells to glucose.

The multiplicity of hypothesis to explain electrical bursting in glucose stimulated *β*-cells underscores the lack of consensus on the actual cellular mechanism that drives the process (25). In *β*-cells, intracellular calcium oscillates with the same frequency observed in bursting even in cells where the electrical activity has been blocked. It is thus natural to consider that intracellular oscillations of calcium determine the rhythm of bursting. In fact, since Chay and Keizer (141) virtually all theoretical models of electrical bursting are based, at least in part, on actions caused by the cytosolic oscillation of Ca^2+^. However, there has been so far no convincing way to link the oscillations of the Ca^2+^ concentration in the cytosol or in the lumen of the ER with the electrical oscillations of the PM.

An important element though has been overlooked in all the previous studies and that is the effect the intracellular dynamics of calcium has on the electric potential of the ER membrane (V*_ER_*). The pumping of Ca^2+^ into the ER and the associated counterion flows generate ionic gradients, mainly of Ca^2+^ and K^+^, across the ER membrane (171, 175, 176). At steady state, these concentration gradients are equilibrated by the electric potential across the ER membrane (177), much as the PM potential equilibrates the ionic gradients across the plasma membrane. The opening of IP3R and other calcium channels and the consequent release of calcium into the cytosol causes the depolarization of the ER membrane. The oscillations of cytosolic and lumenal [Ca^2+^] are therefore associated with parallel oscillations of the ER membrane potential (178). Changes of the ER membrane potential may affect the activity of other components of the cellular machinery. V_*ER*_, for instance, may represent the actual factor that alters the electrical conductance of the store operated channels in the PM that are regulated by the release of calcium from the ER.

### 1.3 Electrical Excitability of the Endoplasmic Reticulum Membrane

The endoplasmic reticulum is a multifunctional organelle formed by a continuous membrane complex that extends from the plasma membrane to the nucleus in all eukaryotic cells (179–182).

Based only on ultrastructural analysis, Ruska et al. postulated in 1958 that the intracellular membranes …*must possess transmembrane potentials and that they serve to the conduction of excitation as does the plasma membrane* (183).

The involvement of the ER in intracellular signaling has since been thoroughly confirmed. A wide variety of physiological processes, including fertilization, cell proliferation and differentiation, the immune response, even learning and memory in the nervous system, are initiated and controlled by the spatiotemporal dynamics of cytosolic calcium that includes calcium oscillations and propagating waves (184, 185). This complex dynamics depends entirely on the ER.

The cytosolic oscillations and waves of calcium are normally initiated by an elevation of IP3 concentration that causes the release of calcium from the ER through IP3R calcium channels. The singular biophysical properties of the IP3R are the main cause of the nonlinear dynamics of calcium inside the cell. The channels display a biphasic response to calcium. At steady state, the open probability of an IP3R is a bell-shaped function of [Ca^2+^]_*CYT*_. This means that the channel is initially activated by the increase of [Ca^2+^]_*CYT*_ but after a critical level further increases of the concentration of free calcium induce the closing of the channel (46, 186–188). In addition to the bell-shaped response, the activation and inactivation of the channel operate in two different time scales, with an almost instantaneous activation and a much slower inactivation (189–192).

Several mathematical models have been developed to analyze how the biophysical properties of the IP3Rs generate the complex dynamics of calcium observed in stimulated cells (193). One of the earliest and most influential is the kinetic model of DeYoung and Keizer (194). This model assumed an IP3R formed by three identical subunits each containing two binding sites for calcium and one for IP3. The subunits therefore can transition between eight different substates. The model included the kinetic equations required to describe all these transitions. Although mathematically complex, it was able to reproduce the most significant behavior of IP3R.

A simplification of the model was developed by Atri et al. (195) and by Li and Rinzel (196). These authors integrated individual kinetic variables used by DeYoung and Keizer into a single gating variable modeled after the inactivation variable h of the Hodgkin and Huxley model (142). Tang et al. (197) have shown that these simplified models are mathematically equivalent to the original DeYoung and Keizer formulation.

In the Li and Rinzel model, the flow of calcium through the IP3R channel, *J*_*IP3R*_, is given by the equation:

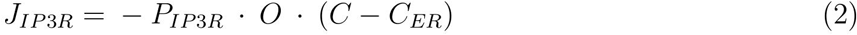

*P*_*IP3R*_ being the total permeability of the IP3R channels in the cell, *C* the cytosolic concentration of calcium and *C*_*ER*_ the concentration of calcium in the lumen of the ER.

*O* corresponds to the fraction of IP3R channels that are open at a given time. This fraction or open probability is determined by the concentrations of IP3 and calcium in the cytosol according to the expression:

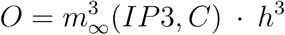

where *IP3* stands for the concentration of IP3 in the cytosol.

*m*_∞_ represents the equilibrium value of the activation gate. It is a function of the IP3 and calcium concentrations in the cytosol

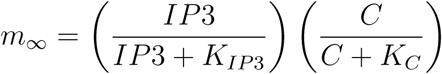

where *K*_*IP3*_ and *K*_*C*_ correspond to the Michaelis-Menten constants for the binding of IP3 and Ca^2+^ to the corresponding activating binding sites in the IP3R channel. Both bindings are assumed to be instantaneous.

*h* is the slow inactivation variable. Its rate of change depends on the time constant *τ* according to the expression:

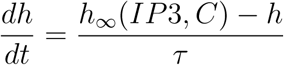

*h*_∞_ is the equilibrium value for the inactivating gate. Its value is

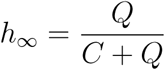

with Q being the effective Michaelis-Menten constant characterizing the channel’s Ca^2+^ inactivation gate.

The Hodgkin-Huxley formalism used to develop this simplified model made evident the parallelism that exists between the IP3Rs in the ER membrane and the voltage-dependent Na^+^ channels (Na_*V*_) in the plasma membrane of excitable cells. In the firing of an action potential, a single variable V, the plasma membrane potential, first triggers the opening of the Na_V_ channel and then, at higher values, the same variable causes the slow closing of the channel. The activation of voltage-dependent K_*V*_ channels then restores the resting conditions. In the ER the single variable [Ca^2+^]_*CYT*_, like V in the PM, first triggers the opening of the IP3R channel and causes a similar excitatory transient. Then when [Ca^2+^]_*CYT*_ reaches a critical point it causes the slow closing of the channel allowing the SERCA pumps to restore the resting conditions. This parallelism inspired Li and colleagues to propose the concept of calcium excitability of the ER as the mechanism underlying the generation of cytosolic calcium oscillations and waves (198, 199).

In their model though, Li and Rinzel (196) assumed that the flow of calcium in and out of the ER has no significant effect on the electric potential across the ER membrane and thus the concentration gradient of calcium, *C* − *C*_*ER*_, is the only factor that drives the flow of the ion through the IP3R channel. However, if the effect on the ER membrane potential is taken into account, then the flow of calcium is driven by its electrochemical gradient across the membrane and the current of calcium, *I*_*CaIP3R*_, through the IP3R would be:

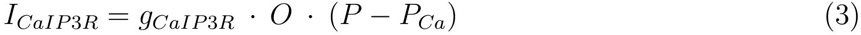

*g*_*CaIP3R*_ represents the maximal conductance of the IP3R channels in the cell. *O* is the proportion of the channels open, as in Eq. 2. *P* represents the electric potential across the ER membrane and *P*_*Ca*_ is the reversal potential of calcium as determined by the Nernst equation:

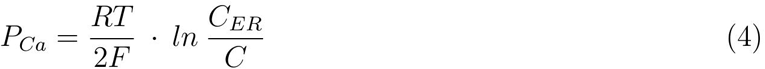

With this formulation, the Li and Rinzel concept of calcium excitability of the ER becomes generalized to the concept of electrical excitability of the ER membrane.

## 2 Mathematical Model

I have developed a minimal mathematical model of a pancreatic *β*-cell in which the excitable ER membrane is assumed to be able to regulate the electrical conductance of the PM. The goal is to test the hypothesis that the oscillations of the ER membrane potential (V_*ER*_) represent the main force driving electrical bursting in glucose stimulated *β*-cells.

The structure of the model is similar to the ER-based model of *β*-cell developed by Teresa Chay (169, 170). A significant difference though is the way the store operated current that generates the slow PM potential oscillations is regulated. Whereas in the Chay model the current is activated by the depletion of calcium in the ER, here it is the depolarization of the ER membrane, caused by the release of calcium, what activates the pacemaker current. The V_*ER*_-regulated channel assumed to be opened by the depolarization of the ER membrane is modeled after the store operated channel TRPM5, known to be involved in the generation of electrical bursting (114).

The model consists of four sets of equations. The first two describe the electric potential across the plasma and endoplasmic reticulum membranes. A third set of equations summarizes the balance of calcium in the different subcellular compartments. Finally, the last equation describes the dynamics of inositol 1,4,5 trisphosphate in the cytosol.

### 2.1 Voltage across the plasma membrane

The equations describing the rate of change of the electric potential across the PM and ER membrane are based on the equation for the PM voltage (Eq. 1) of the Chay and Keizer model (141).

The voltage across the plasma membrane is determined by the current across five different ion channels. Two voltage-dependent channels are responsible for the generation of action potentials, Ca_*V*_ that carries the *I*_*CaV*_ current, and K_*V*_ that conducts the delayed rectified *I*_*K*_ current. A second calcium channel, C*a*_*P*_, carries the *I*_*CaP*_ current. This corresponds to the V_ER_-activated current hypothesized by the model. This is a small current that could also have been modeled as Na^+^ - specific or even non-selective without altering significantly the behavior of the model. The K_*ATP*_ channel, that carries the *I*_*K*(*ATP*)_ current, incorporates the direct regulation of the PM potential by glucose into the model. Finally a nonregulated calcium channel *Ca*_*l*_ carries a leak current *I*_
*Cal*_ that balances the activity of the Ca^2+^-ATPase pump when the cell is at rest and prevents the depletion of the cell.

The rate of change of the voltage *V* across the plasma membrane is given by the expression:

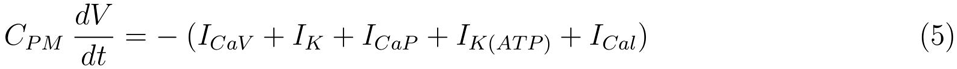

Where *C*_*PM*_ represents the capacitance of the plasma membrane, and the currents are defined by the following equations:

The voltage-dependent calcium current is

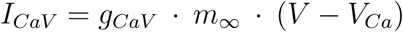

where *m*_∞_ is the steady state value of the gating variable *m* for the Ca_*V*_ channel. The steady state gating variables such as *m*_∞_ are sigmoidal functions of voltage. They are usually represented by a Boltzmann expression:

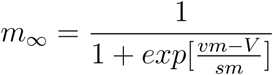

which contains two parameters, a half-maximal potential *v*_*m*_ and the gating slope *s*_*m*_ at *v*_*m*_. The gating of the Ca_*V*_ channel is assumed to be instantaneous and therefore the gating variable *m* is independent of time and always equal to the steady state value *m*_∞_.

The delayed rectifier K^+^ current is

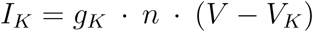

The K_*V*_ channel responds to changes of V with a delay determined by the rate of change of the gating variable *n*:

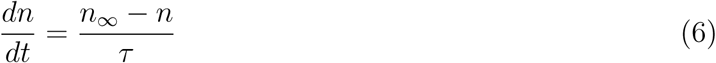

where *n*_∞_ is given by the expression:

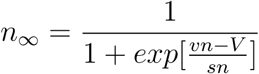

and *τ* represents the relaxation time constant.

The V*_ER_*-dependent calcium current is

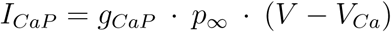

where *p*_∞_ is given by:

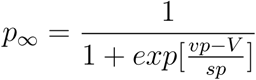

The activation of this channel is also assumed to be instantaneous with the gating variable *p* always equal to *p*_*∞*_.

The ATP-dependent potassium current is

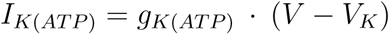

Finally, the Ca^2+^ leak current is

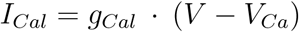

The values of the reverse potential for Ca^2+^, *V*_*Ca*_, and K^+^, *V*_*K*_, and other parameters used in the model are given in Table 1.

**Table 1:**
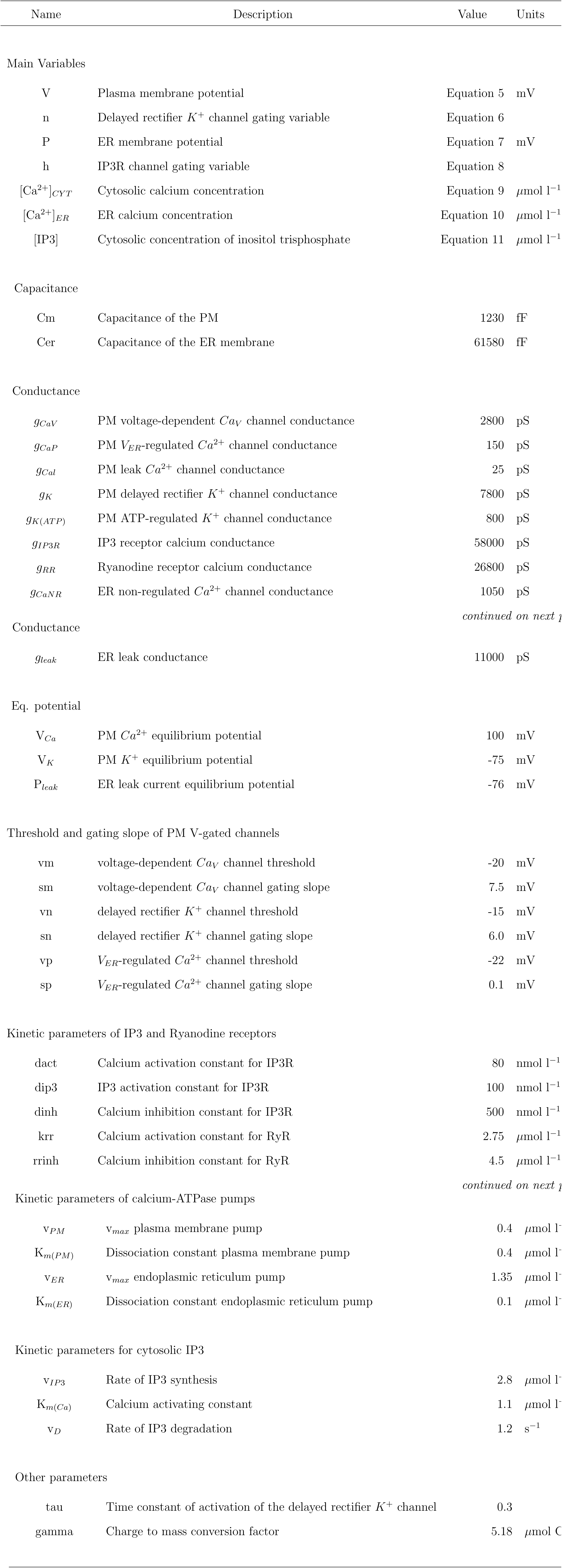
Variables and Parameters

| Name | Description | Value | Units |
| --- | --- | --- | --- |
| Main Variables |  |  |  |
| V | Plasma membrane potential | Equation 5 | mV |
| n | Delayed rectifier $K^+$ channel gating variable | Equation 6 | |
| P | ER membrane potential | Equation 7 | mV |
| h | IP3R channel gating variable | Equation 8 |  |
| $[Ca^{2+}]_{CYT}$ | Cytosolic calcium concentration | Equation 9 | $\mu\text{mol l}^{-1}$ |
| $[Ca^{2+}]_{ER}$ | ER calcium concentration | Equation 10 | $\mu\text{mol l}^{-1}$ |
| [IP3] | Cytosolic concentration of inositol trisphosphate | Equation 11 | $\mu\text{mol l}^{-1}$ |
| Capacitance |  |  |  |
| Cm | Capacitance of the PM | 1230 | fF |
| Cer | Capacitance of the ER membrane | 61580 | fF |
| Conductance |  |  |  |
| $g_{CaV}$ | PM voltage-dependent $Ca_V$ channel conductance | 2800 | pS |
| $g_{CaP}$ | PM $V_{ER}$-regulated $Ca^{2+}$ channel conductance | 150 | pS |
| $g_{Cal}$ | PM leak $Ca^{2+}$ channel conductance | 25 | pS |
| $g_K$ | PM delayed rectifier $K^+$ channel conductance | 7800 | pS |
| $g_{K(ATP)}$ | PM ATP-regulated $K^+$ channel conductance | 800 | pS |
| $g_{IP3R}$ | IP3 receptor calcium conductance | 58000 | pS |
| $g_{RR}$ | Ryanodine receptor calcium conductance | 26800 | pS |
| $g_{CaNR}$ | ER non-regulated $Ca^{2+}$ channel conductance | 1050 | pS |

Table 1: *Variables and Parameters continued*
| Name | Description | Value | Units |
| --- | --- | --- | --- |
| Conductance |  |  |  |
| $g_{leak}$ | ER leak conductance | 11000 | pS |
| Eq. potential |  |  |  |
| $V_{Ca}$ | PM $Ca^{2+}$ equilibrium potential | 100 | mV |
| $V_K$ | PM $K^+$ equilibrium potential | -75 | mV |
| $P_{leak}$ | ER leak current equilibrium potential | -76 | mV |
| Threshold and gating slope of PM V-gated channels |  |  |  |
| vm | voltage-dependent $Ca_V$ channel threshold | -20 | mV |
| sm | voltage-dependent $Ca_V$ channel gating slope | 7.5 | mV |
| vn | delayed rectifier $K^+$ channel threshold | -15 | mV |
| sn | delayed rectifier $K^+$ channel gating slope | 6.0 | mV |
| vp | $V_{ER}$-regulated $Ca^{2+}$ channel threshold | -22 | mV |
| sp | $V_{ER}$-regulated $Ca^{2+}$ channel gating slope | 0.1 | mV |
| Kinetic parameters of IP3 and Ryanodine receptors |  |  |  |
| dact | Calcium activation constant for IP3R | 80 | nmol l<sup>-1</sup> |
| dip3 | IP3 activation constant for IP3R | 100 | nmol l<sup>-1</sup> |
| dinh | Calcium inhibition constant for IP3R | 500 | nmol l<sup>-1</sup> |
| krr | Calcium activation constant for RyR | 2.75 | μmol l<sup>-1</sup> |
| rrinh | Calcium inhibition constant for RyR | 4.5 | μmol l<sup>-1</sup> |

Table 1: *Variables and Parameters continued*
| Name | Description | Value | Units |
| --- | --- | --- | --- |
| Kinetic parameters of calcium-ATPase pumps |  |  |  |
| $v_{PM}$ | $v_{max}$ plasma membrane pump | 0.4 | $\mu\text{mol l}^{-1}$ |
| $K_{m(PM)}$ | Dissociation constant plasma membrane pump | 0.4 | $\mu\text{mol l}^{-1}$ |
| $v_{ER}$ | $v_{max}$ endoplasmic reticulum pump | 1.35 | $\mu\text{mol l}^{-1}$ |
| $K_{m(ER)}$ | Dissociation constant endoplasmic reticulum pump | 0.1 | $\mu\text{mol l}^{-1}$ |
| Kinetic parameters for cytosolic IP3 |  |  |  |
| $v_{IP3}$ | Rate of IP3 synthesis | 2.8 | $\mu\text{mol l}^{-1}$ |
| $K_{m(Ca)}$ | Calcium activating constant | 1.1 | $\mu\text{mol l}^{-1}$ |
| $v_D$ | Rate of IP3 degradation | 1.2 | $\text{s}^{-1}$ |
| Other parameters |  |  |  |
| tau | Time constant of activation of the delayed rectifier $K^+$ channel | 0.3 | |
| gamma | Charge to mass conversion factor | 5.18 | $\mu\text{mol C}^{-1}$ |

### 2.2 Voltage across the ER membrane

The equations for the electrical activity of the ER membrane are based on the Li and Rinzel model of the ER (196), with the modifications indicated in Eq. 3 in the Introduction.

The voltage across the ER membrane (P in the equations) depends on three calcium currents and a leak current according to the expression:

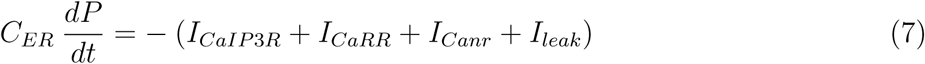

where *C*_*ER*_ represents the capacitance of the ER membrane.

The Ca^2+^ current *I*_*CaIP3R*
_ across the IP3R is

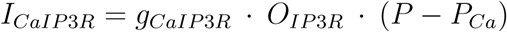

where *O*_*IP3R*_ represents the open probability of the IP3R channel. The expression for *O*_*IP3R*_ incorporates the fast activation of the channel by Ca^2+^ (*a*) and IP3 (*b*) and the slow inactivation by elevated concentrations of Ca^2+^ (*h*)

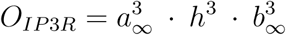

The activation functions *a*
_∞_ and *b*
_∞_ are

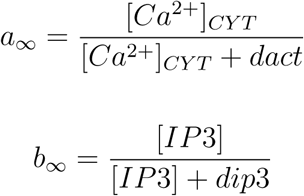

The rate of change of the gating variable *h* is

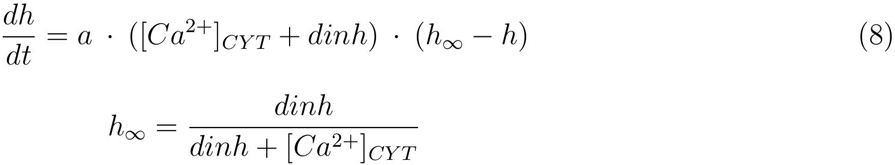

The current *I*_*CaRR*_ across the RyR channel is also activated by small elevations of [Ca^2+^]_*CYT*_ and inhibited by high concentrations of Ca^2+^. The expression for the current is

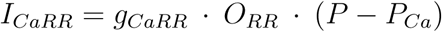

where *O*_*RR*
_ is given by the expression:

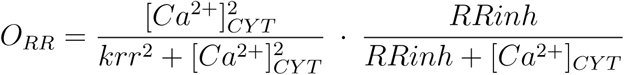

The third Ca^2+^ current included in the model, *I*_*Canr*_, is a non-regulated current that depends only on the electrochemical gradient for Ca^2+^ across the ER membrane. This current provides the observed leak of calcium from the lumen of the ER, responsible for the depletion of the ER when the SERCA pump is inhibited. The expression for the current is

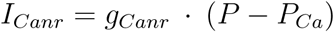

Finally, *I*_*leak*_ is a Hodgkin and Huxley type leak current (142). This leak current integrates all the counterion flows, glutamate, chloride (200), but mainly potassium (201, 202), that take place in response to the uptake and release of calcium from the lumen of the ER. The expression for the current is

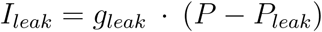

Given the predominant role of K^+^ in the leak current, the equilibrium potential *P*_*leak*_ is assumed to be close to the equilibrium potential for K^+^ in the ER membrane, that lacking better information is taken to be similar to the equilibrium potential *V*_*K*_ across the PM. The equilibrium potential for Ca^2+^, *P*_*Ca*_, is given by the Nernst equation

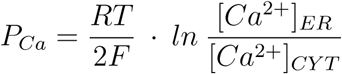

The values used in the simulations for the ionic conductances and other parameters in the equations are given in Table 1.

### 2.3 Calcium balance equations

The cell in this model is assumed to have 5 *μ*m of radius and thus a surface of 314 *μ*m^2^ and a volume of 523 f liter. Beta cells are known to have a great calcium buffering capacity in both the cytosol and the lumen of the ER. Bertram et al. (172) introduced the concept of effective volume, *V*_*eff*_, to deal with calcium buffering in their model of a *β*-cell. For a ratio *f*_*i*_ of free to total calcium in a cellular compartment of volume *V*_*i*_, the effective volume of the compartment is defined as:

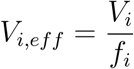

Using this concept, the model cell is assumed to have a cytosolic effective volume, *V*_*c,eff*_ of 2.7 pico liters. This corresponds to a proportion of free to total calcium of the order of ten percent. The effective volume of the lumen of the ER, *V*_*er,eff*
_, is assumed to be 0.5 pico liters, i.e. 5.4 times smaller than the volume of the cytosol.

With these assumptions, the change of cytosolic calcium concentration is related to the calcium fluxes across the PM (*J*_*PM*_) and the ER membrane (*J*_*ER*_) by the expression:

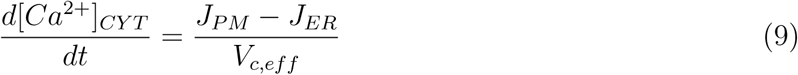

And the change of the calcium concentration in the lumen of the ER:

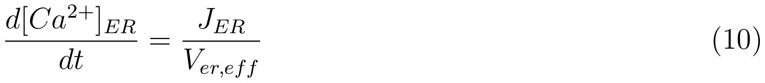

The flux of calcium across the plasma membrane, *J*_*PM*_, is due to the calcium currents already described and to the extrusion of calcium by a calcium-ATPase pump:

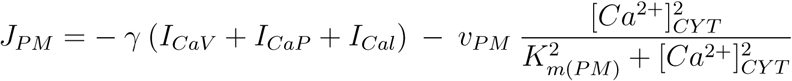

A similar equation describes the calcium flux, *J*_*ER*_, across the ER membrane:

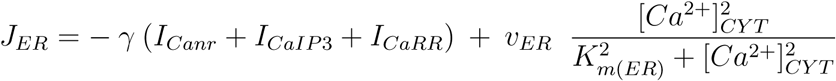

where *v*_*PM*_, *v*_*ER*_ and *K*_*m*(*PM*)_, *K*_*m*(*ER*)_ represent the maximum velocity and the Michaelis-Menten constant for the PM and ER ATPase pumps. A second order kinetics for these pumps has been assumed. The parameter γ transforms the flow of electrical charge into flow of mass, assuming one mole of calcium has zF, i.e. 193000, Coulombs.

The values of the parameters used in the simulations are shown in Table 1.

### 2.4 Dynamics of IP3

The final equation of the model describes the dynamics of inositol thisphosphate in the cytosol. The cytosolic concentration of IP3, [*IP3*], is determined by the rate of synthesis and the rate of degradation of the phospholipid according to the expression:

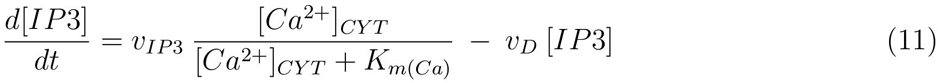

The rate of synthesis v_*IP*3_ is set to reflect the activity of the different forms of PLC activated by glucose as well as by neural and hormonal factors such as ACh and GLP-1. All PLC isoforms are regulated by calcium (42). This regulation is included in the model through the *K*_*m*(*Ca*)_ parameter.

The degradation of IP3 in cells can proceed in two different ways. IP3 can be either dephosphorylated by IP3 5-phosphatase or phosphorilated by IP3 kinase (203). The activity of the two enzymes is integrated in the model into the single rate of degradation *v*_*D*_. This process is assumed to be independent of calcium.

The model equations have been integrated numerically using a step adaptable Gear method implemented in the free software package XPPAUT developed by Bard Ermentrout (204).

## 3 Results

### 3.1 Biphasic response to glucose

Pancreatic *β*-cells display a biphasic response to glucose (11). Upon the elevation of blood glucose, there is first a short acute secretory phase believed to be required for changing the metabolic program of the liver from main glucose producer to glucose sink. This acute initial response is followed by a reduced but sustained secretion of insulin that stabilizes the normal levels of glucose in the bloodstream (205, 206). The electrical counterpart of the two phases are the continuous and the bursting patterns of action potential firing (14). The biphasic structure of the *β*-cell response is critical for the maintenance of normoglycemia. The loss of the first phase is one of the earliest detectable abnormalities in the development of T2DM (207–209).

An important test of the relevance of a theoretical model of *β*-cells is to see if it can reproduce this biphasic response. In pancreatic *β*-cells, the elevation of glucose induces the closing of K_*ATP*_ channels and stimulates the production of IP3. I have simulated these effects in the *β*-cell model by changing the value of the *g*_*K*(*ATP*)_ (conductance of the K_*ATP*_ channel) and v_*IP*3_ (rate of IP3 synthesis) parameters in a two step integration. The result of the simulation can be seen in Figure 1.

**Figure 1:**
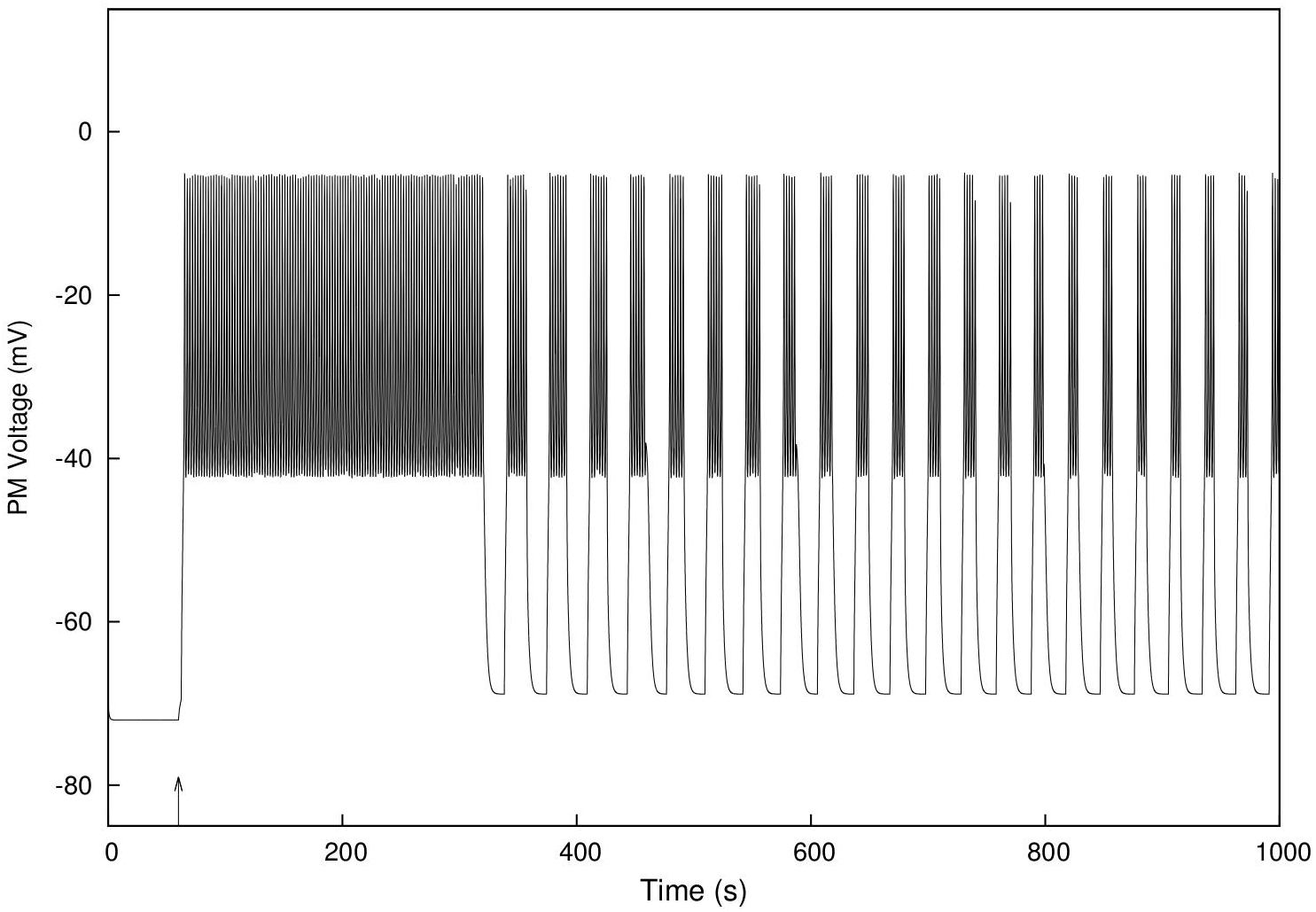
Biphasic response of the model cell to glucose. Equations 5 to 11 were integrated in two steps, starting from the initial conditions shown in Table 2. The parameters had the values shown in Table 1, except for *g_K(ATP)_*, the conductance of the K_*ATP*_ channels, and v_*IP*3_, the rate of synthesis of IP3. For the initial step of the integration, up until the arrow, *g*_*K*(*ATP)*_ was set to 1600 fS and v_*IP*3_ to 1.6 *μ*mol l^−1^ *s*^−1^. These values correspond in the model to open K_*ATP*_ channels and no induction of PLC, the conditions of non-stimulated cells. At the arrow, the parameters were changed to 800 fS and 2.8 *μ*mol l^−1^ *s*^−1^ to simulate the elevation of glucose in the cell environment, and the integration was continued until 1000 seconds. The panel shows the value of the plasma membrane potential during the whole integration. It can be observed that, upon the change of parameters, the PM becomes depolarized and starts firing action potentials. The firing is initially continuous but after five minutes it is periodically interrupted by intervals of PM hyperpolarization. The model cell therefore exhibits a biphasic response, with a continuous and a bursting phase, comparable with the biphasic response observed experimentally in pancreatic *β*-cells. In the model, the bursts last between 50 and 40 seconds and are separated by intervals of electrical inactivity of 30 to 50 seconds. The frequency of bursting is of 1.4 to 1.7 bursts per minute.

The integration was started with parameter values corresponding to a non-stimulated cell, open K_*ATP*_ channels and no activation of IP3 synthesis. In these conditions, the plasma membrane is hyperpolarized (Figure 1, left). At the time indicated by the arrow, the parameters were changed to levels that simulate a moderate elevation of glucose and the integration continued. The change caused the rapid depolarization of the plasma membrane from the resting −70 mV to a value of about −40 mV, at which point the cell started firing action potentials, fast oscillations of the PM potential between −40 mV and −5 mV. The action potentials were first continuous, but after 5 minutes the PM became periodically hyperpolarized interrupting the firing of action potentials. This generated a bursting-like electrical response (Figure 1, right).

The model cell therefore is capable of reproducing the biphasic response observed in *β*-cells. The duration of the initial continuous phase is of 5 minutes, similar to the duration measured in vivo (14, 210). The bursting phase in the model has a frequency of about 2 bursts per minute, comparable with the bursting frequencies of glucose stimulated islets in vivo (121, 122) and in culture (9, 10).

The evolution of the electric potential of the ER membrane during the simulation is shown in Figure 2B. The ER membrane was initially hyperpolarized (-60 mV) and became depolarized upon the simulated elevation of glucose. The membrane then remained depolarized for about 5 minutes, i.e. during the whole initial phase, until it started to oscillate periodically between −50 and −10 mV. Figure 3 shows a closed up detail of the simulation with the oscillations of the V*_PM_* overimposed to those of the ER membrane potential.

**Figure 2:**
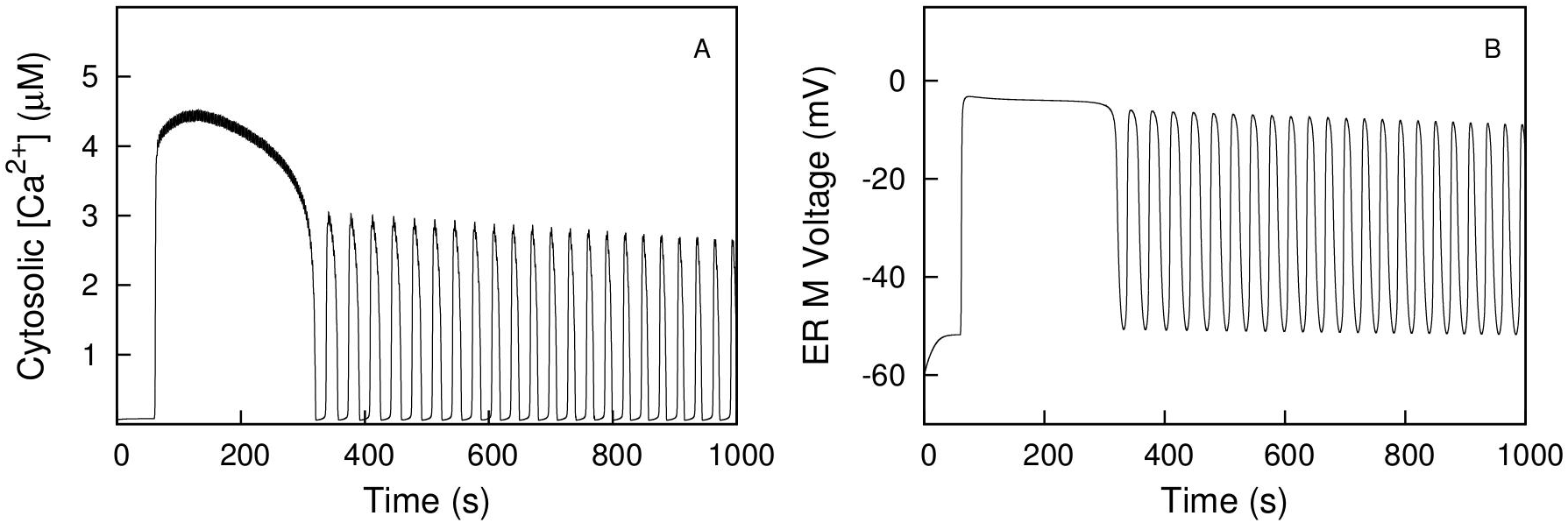
Biphasic response of model cell to glucose. Cytosolic calcium concentration (Panel A) and electric potential across the ER membrane (Panel B). Data from the simulation described in Figure 1. Both [Ca^2+^] and V_*ER*_ display the same biphasic pattern exhibited by the PM voltage.

**Figure 3:**
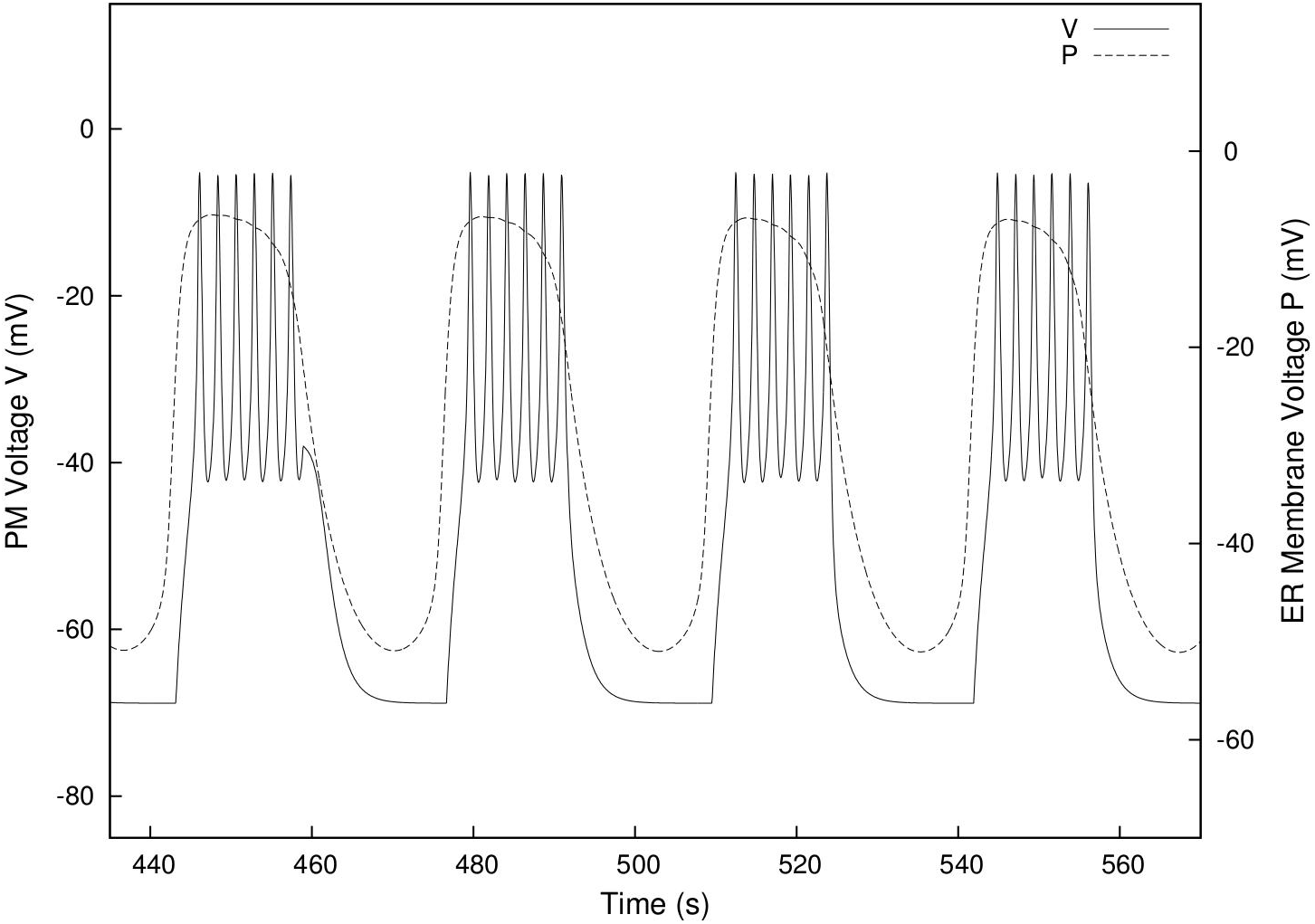
Detail of electrical bursting by the model cell. Value of the voltage across the PM (V) and the ER membrane (P). Data from the simulation described in Figure 1. Note the synchrony of the electrical bursts of the plasma membrane and the oscillations of the ER membrane potential. A similar synchrony also takes place with the oscillations of the cytosolic concentration of calcium (not shown).

**Table 2:** Initial Conditions

|  |  |  |
| --- | --- | --- |
| V | Plasma membrane potential | -70 mV |
| n | Delayed rectifier $K^+$ channel gating variable | 0.001 |
| P | ER membrane potential | -60 mV |
| h | IP3R channel gating variable | 0.6 |
| $[Ca^{2+}]_{CYT}$ | Cytosolic calcium concentration | 0.1 μmol l<sup>-1</sup> |
| $[Ca^{2+}]_{ER}$ | ER calcium concentration | 1080 μmol l<sup>-1</sup> |
| [IP3] | Cytosolic concentration of IP3 | 0.01 μmol l<sup>-1</sup> |

The concentration of cytosolic calcium during the simulation is shown in Figure 2A. It has also a biphasic structure with an initial sustained elevation followed by periodic oscillations, as has been observed experimentally in *β*-cells (120). The concentration of calcium in the lumen of the ER decreases during the simulation from an initial value of 1 mmol l ^−1^ to a value of 600 *μ*mol l ^−1^ after the 1000 s of integration.

### 3.2 Origin of the biphasic response

The effective control of the blood levels of glucose requires the secretion of insulin in two differentiated phases. As mentioned above, the loss of the initial acute phase of secretion is an early marker of a developing diabetes. In spite of its clinical importance, the mechanism responsible for the two different patterns of secretion is still poorly understood (211). It is therefore important to analyze in detail how the biphasic response originates in the current model, to see if the mechanism operating in the model cell can help to understand the generation of the biphasic response in vivo.

To better dissect the process, I have studied the dynamics of the ER in a reduced model in which the ER membrane has been isolated from the effects of the electrical activity of the plasma membrane. In this context, it is easier to analyze the specific responses of the ER to external stimuli. This autonomous system consists of equations 7 to 11 of the general model. By removing the equations 5 and 6 that deal with the PM potential, the reduced system becomes de facto a model of a non-excitable cell.

Using these equations, I have analyzed how the change in two parameters alters the dynamic state of the ER membrane. One of these parameters is v_ER_, the rate of calcium uptake by the SERCA pump, a component of the ER membrane that is known to be regulated by several signaling systems and is also the target of pharmacological agents such as the inhibitor thapsigargin. The other parameter is v_*IP*3_, the rate of synthesis of IP3. As discussed in the Introduction, this second messenger is regulated by glucose and several neurohormonal factors and plays a central role in the control of insulin secretion. The changes of these two parameters therefore reflect the effects of the main physiological stimuli that operate on *β*-cells. I have performed this study first in a closed system, a cell that does not allow the flow of calcium or any other ion across the PM, and then in an open system where calcium can leave the cell.

The behavior of the closed system is shown in Figures 4 and 5.

**Figure 4:**
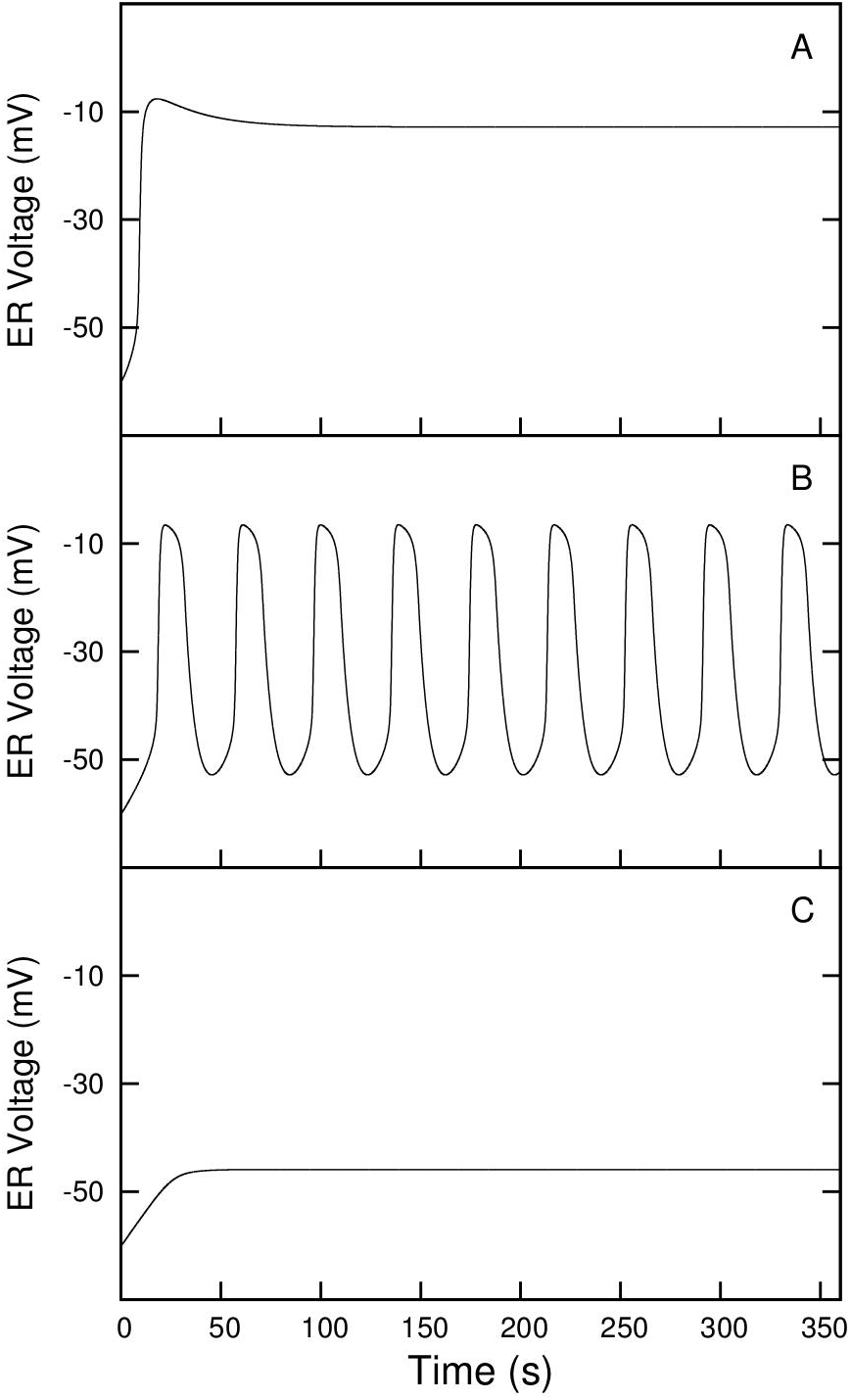
Dynamics of the ER membrane potential in a closed cell. A model system consisting of equations 7 to 11 has been used to simulate a non-excitable, closed cell in which the plasma membrane has no activity (v_*PM*_ = 0). The equations were integrated for the time indicated in the figure, using the parameter values in Table 1 except for v_*PM*_, v_*ER*_ and v_*IP*3_. The integration was started from the initial conditions shown in Table 2. Panel A shows the evolution of the ER membrane potential when the max rate of calcium uptake into the ER (v_*ER*_) is 3.7 *μ*mol l ^−1^ *s*^−1^ and the rate of IP3 synthesis (v_*IP*3_) is 2.5 *μ*mol l ^−1^ *s*^−1^. As can be observed, the ER membrane potential, initially at −60 mV, becomes rapidly depolarized and settles at −10 mV, a stable point of the dynamic system for these parameter values. In panel B the parameters v_*ER*_ and v_*IP*3_ had the values 4.3 *μ*mol l ^−1^ *s*^−1^ and 2.5 *μ*mol 1 ^−1^ *s*^−1^ respectively. In these conditions, the voltage across the ER membrane displays periodic oscillations between −50 and −10 mV. This behavior indicates that the system approaches a stable limit cycle similar to the one shown in Figure 8. Panel C shows the evolution of the ER membrane potential when v_*ER*_ and v_*IP*3_ had the values 4.9 *μ*mol l ^−1^ *s*^−1^ and 2.5 *μ*mol l ^−1^ *s*^−1^. With these parameter values, the ER membrane becomes slightly depolarized and settles to a stable point of −50 mV. The whole set of v_*ER*_ and v_*IP*3_ values that generate each of the three stable conditions are shown in Figure 5.

**Figure 5:**
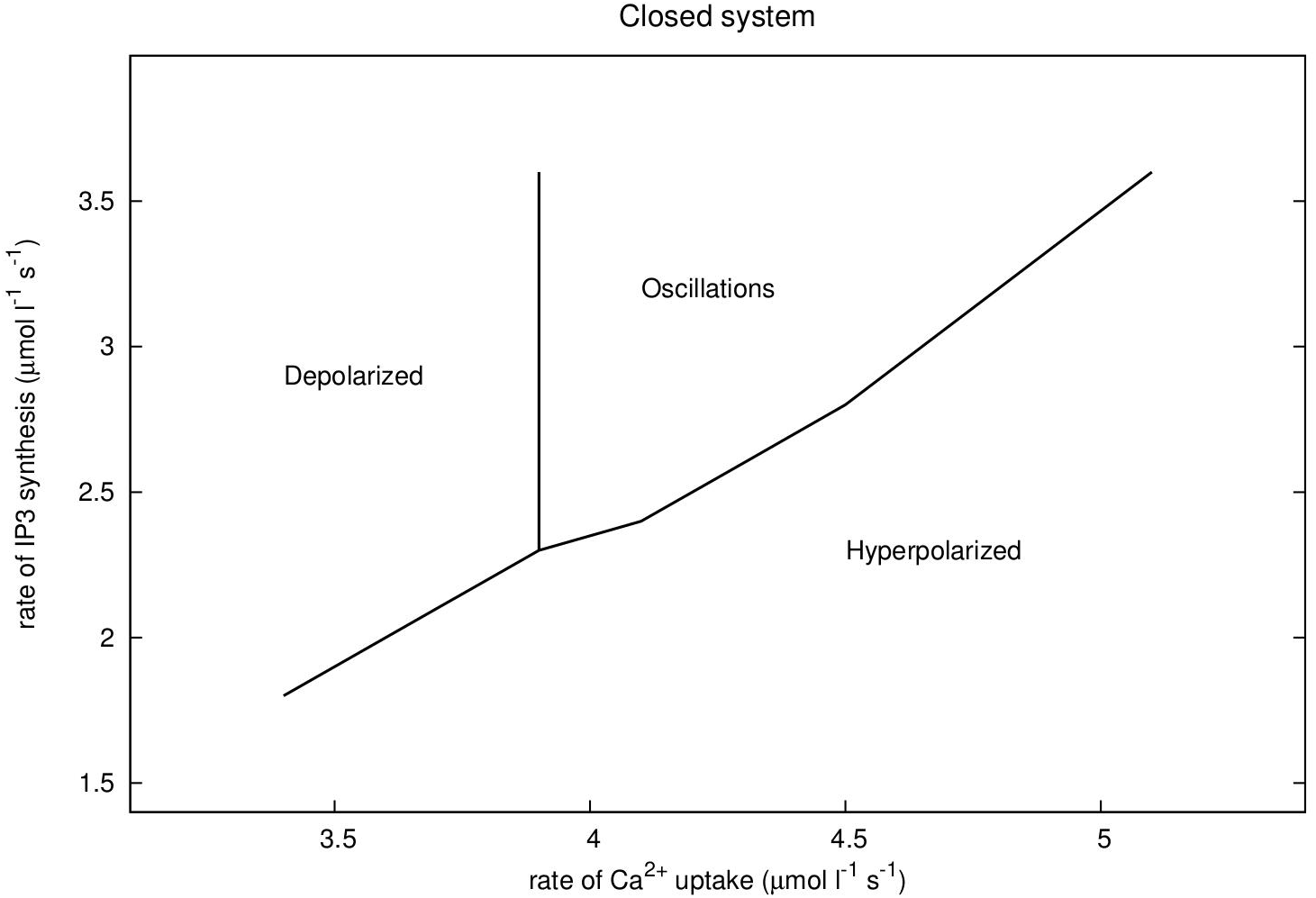
Stability diagram of the ER membrane potential in a closed cell. Equations 7 to 11, in conditions that simulate a non-excitable closed cell (v_*PM*_ = 0), were integrated as described in the legend to Figure 4, using different values of v_*ER*_ and v_*IP*3_. The rate of Ca^2+^ uptake into the ER (v_*ER*_), was varied between 3.4 and 5.5 *μ*mol l ^−1^ *s*^−1^. v_*IP*3_, the rate of IP3 synthesis, was given values between 1.7 and 3.6 *μ*mol l ^−1^ *s*^−1^. For each pair of values, the type of stable condition reached by the system was recorded. Three different stable states were identified (see Figure 4). The diagram shows the sets of v_*ER*_, v_*IP*3_ points that generate each state. The area to the left, labeled Depolarized, represents the values of v_*ER*_, v_*IP*3_ that cause the ER membrane to evolve towards a depolarized stable point, following a trajectory like the one shown in Figure 4, Panel A. The central area, labeled Oscillations, corresponds to the values of the parameters for which the system shows an oscillatory behavior as in Figure 4, Panel B. Finally, the area labeled Hyperpolarized corresponds to the set of parameter values that cause the system to remain in a hyperpolarized stable state as in Figure 4, Panel C. It can be observed that at low levels of IP3 activation, the system only has two possible stable states, a hyperpolarized or a depolarized ER membrane. However, when the rate of IP3 synthesis increases, a new equilibrium state emerges, a limit cycle that causes periodic oscillations of the ER membrane.

Figure 4 shows the three possible behaviors the system may display depending on the value of the parameters v_*ER*_ and v_*IP*3_. At low levels of SERCA pump activity (low v_*ER*_), the elevation of IP3 causes the depolarization of the ER membrane to a stable value of −10 mV (Figure 4, Panel A).

At high levels of SERCA activity (high v_*ER*_), the system reaches a different stable point around −50 mV. The ER membrane basically remains hyperpolarized (Figure 4, Panel C).

Finally, for a set of intermediate v_*ER*_ and v_*IP*3_ values, the system displays a periodic oscillatory behavior ((Figure 4, Panel B). The ER membrane potential oscillates between −50 mV and −10 mV. This is the type of behavior that generates electrical bursting in the whole *β*-cell model.

Figure 5 shows the stability diagram of the closed system in the v_*ER*_ v_*IP*3_ plane. The panel shows the set of v_*ER*_, v_*IP*3_ values that correspond to each of the three equilibrium states that can be reached by the dynamical system, labeled Hyperpolarized, Depolarized and Oscillations. The diagram helps to understand how the change in one parameter affects the behavior of the model cell. The horizontal v_ER_ axis reflects the rate of calcium uptake into the ER. Any change on the SERCA pump activity causes a horizontal displacement in the diagram. When a line is crossed the behavior of the system changes, for instance from being hyperpolarized to become depolarized. Similarly, the vertical v_*IP*3_ axis indicates the balance between IP3 synthesis and degradation. In this case, an important change takes place in the structure of the dynamical system. At low levels of IP3 activation, an elevation of the rate of IP3 synthesis (the v_*IP*3_ parameter) only may cause a transition from the Hyperpolarization to the Depolarization area. The system only has two possible stable states. However, above a threshold value of v*_IP_*_3_ (2.3 *μ*mol l ^−1^ *s*^−1^), a new stable state emerges, labeled Oscillations in the diagram. The system now displays a new behavior, periodic oscillations of the ER membrane potential. This reflects the emergence of a new stable condition in the system, a limit cycle similar to the cycle in the phase-plane analysis shown in Figure 8.

The nonlinear dynamics of the ER in the current model therefore explains how the periodic oscillations of the electric potential of the ER membrane, responsible for electrical bursting, appear. However, the closed system does not provide an explanation for the generation of the biphasic response. To understand this response it is necessary to analyze an open system.

The open system I have analyzed consists basically on the same non-excitable cell used before but with a calcium pump added to the plasma membrane that transports Ca^2+^ out of the cell. In this cell model, the total amount of calcium inside the cell is not constant like in the closed cell but decreases continuously during the simulation. The stability diagram in these conditions (Figure 6) has the same three stable states seen in the closed system but is no longer static, it suffers a leftward displacement in the course of the integration. This generates a new area, labeled Biphasic in the diagram. For values of v_*ER*_ and v_*IP3*_ in this area the system evolves as shown in Figure 7. The ER membrane potential evolves initially to the stable state of this area and thus becomes depolarized. However, when the intracellular level of calcium reaches a critical point, the stability of the system changes from the depolarized fixed point to the stable limit cycle that causes the periodic oscillations of the ER membrane potential. This is the process that generates the biphasic response in the simplified as well as in the whole *β*-cell model (compare Figure 7 with (Figure 2B).

**Figure 6:**
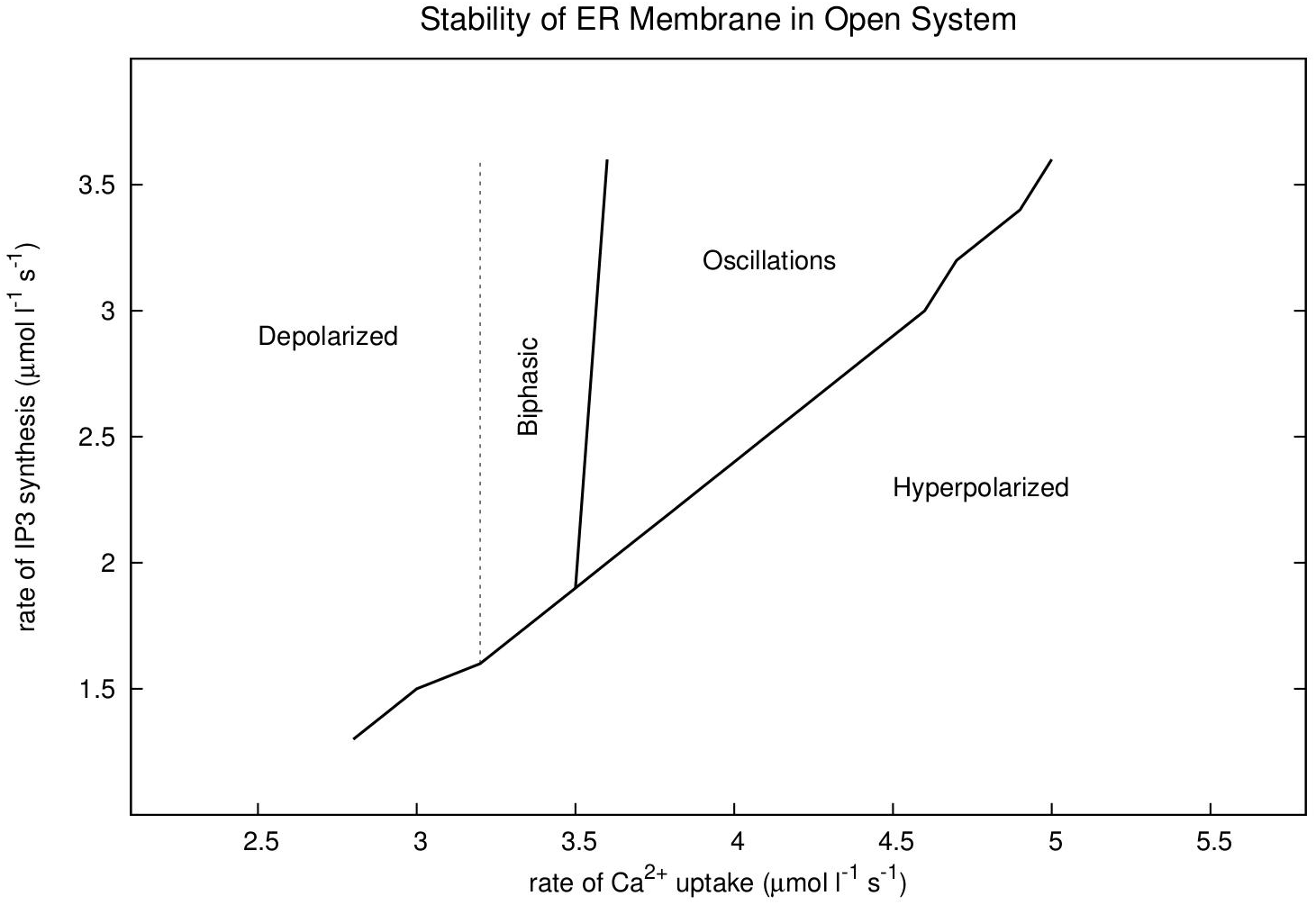
Stability diagram of the ER membrane potential in an open cell. Equations 7 to 11, in conditions that simulate a non-excitable open cell (v_*PM*_ = 0.4 *μ*mol l ^−1^ *s*^−1^), were used to run a series of integrations like those described in the legend of Figure 5. The diagram shows the stability areas found in the integrations. The open system has the same three areas that the closed cell model. They are labeled Depolarized, Oscillations and Hyperpolarized and correspond to the set of v_*ER*_, v_*IP*3_ points that generated trajectories similar to those shown in Figure 4, Panels A, B and C respectively. In addition, a new behavior appeared in the simulations. In the area labeled Biphasic, the trajectory of the system was initially the same as the trajectories in the Depolarized area. The electric potential of the ER membrane changed from the initial hyperpolarization of −60 mV to a depolarized value of −10 mV and remained at this value for several minutes but eventually the trajectory continued to produce an oscillatory behavior similar to the behavior of the system in the Oscillations area. An example of this type of trajectory can be seen in Figure 7. A phase-plane analysis of the trajectory is shown in Figure 8. For the set of parameters in the Biphasic area therefore the reduced open system displays a biphasic response comparable to the response seen in the whole cell model (compare Figure 7 with Figure 2B).

**Figure 7:**
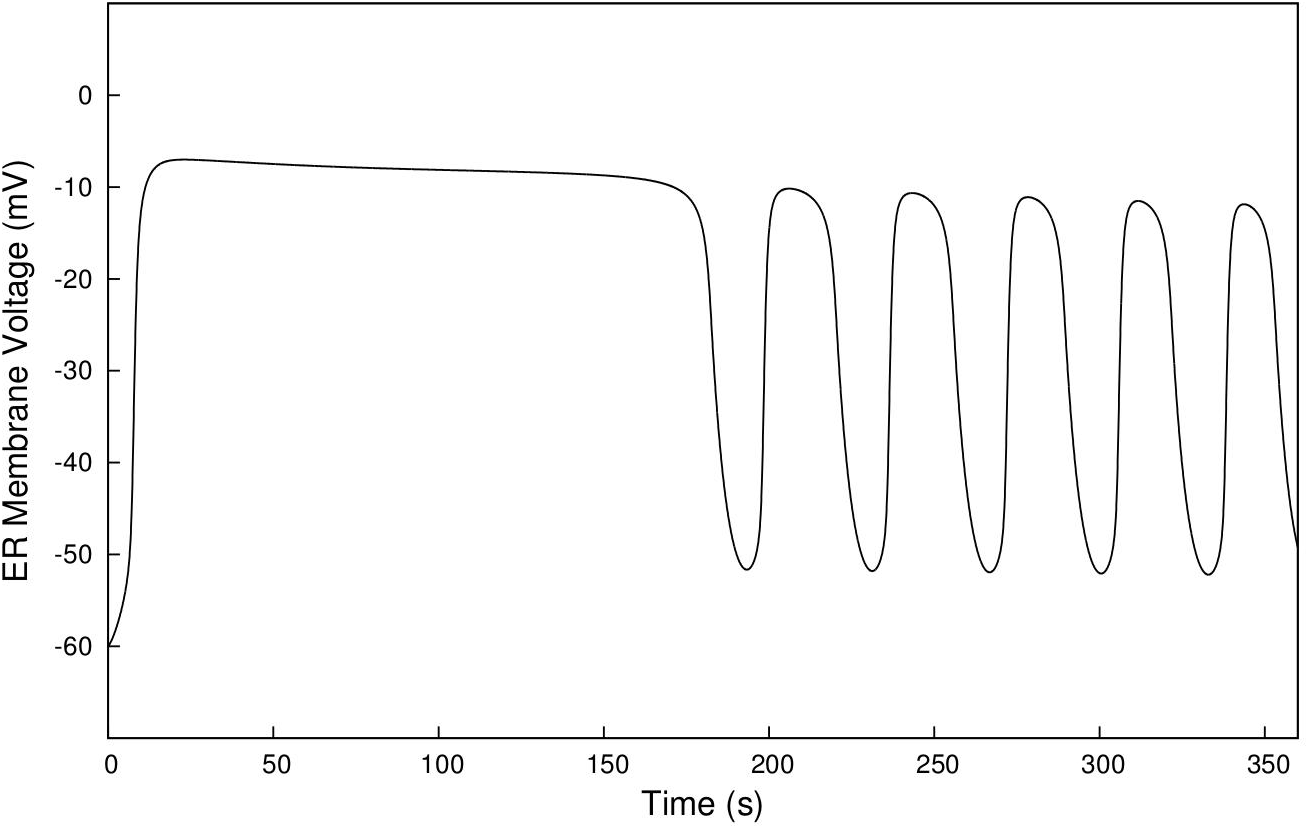
Biphasic pattern of the ER membrane potential in an open cell. Equations 7 to 11 were integrated for the time indicated in the figure, using the parameter values in Table 1 except for v_*ER*_ and v_*IP*3_. The value of the max velocity of the SERCA pump (v_*ER*_) was 3.4 *μ*mol l ^−1^ *s*^−1^ and the value of the rate of IP3 synthesis (v_*IP*3_) was 2.5 *μ*mol l ^−1^ *s*^−1^. The integration was started from the initial conditions shown in Table II. The figure displays the evolution of the ER membrane potential during the simulation. It can be observed that the ER membrane depolarizes rapidly to a value of −10 mV and remains depolarized for about 3 minutes. Then the electric potential of the ER membrane starts to oscillate between −50 mV and −10 mV.

This biphasic response is also illustrated in the phase plane analysis shown in Figure 8, where the orbit of the ER membrane potential (P) in the h, P plane can be observed. The trajectory of the system goes first from the initial point at 0.6, −60 mV to the depolarized stable point in 0.1, −5 mV. The trajectory remains there for a few minutes, but eventually continues to the stable limit cycle where it remains for the rest of the integration. The cycling around the limit cycle generates the periodic oscillations of the ER membrane potential that appear in the second part of the biphasic response.

**Figure 8:**
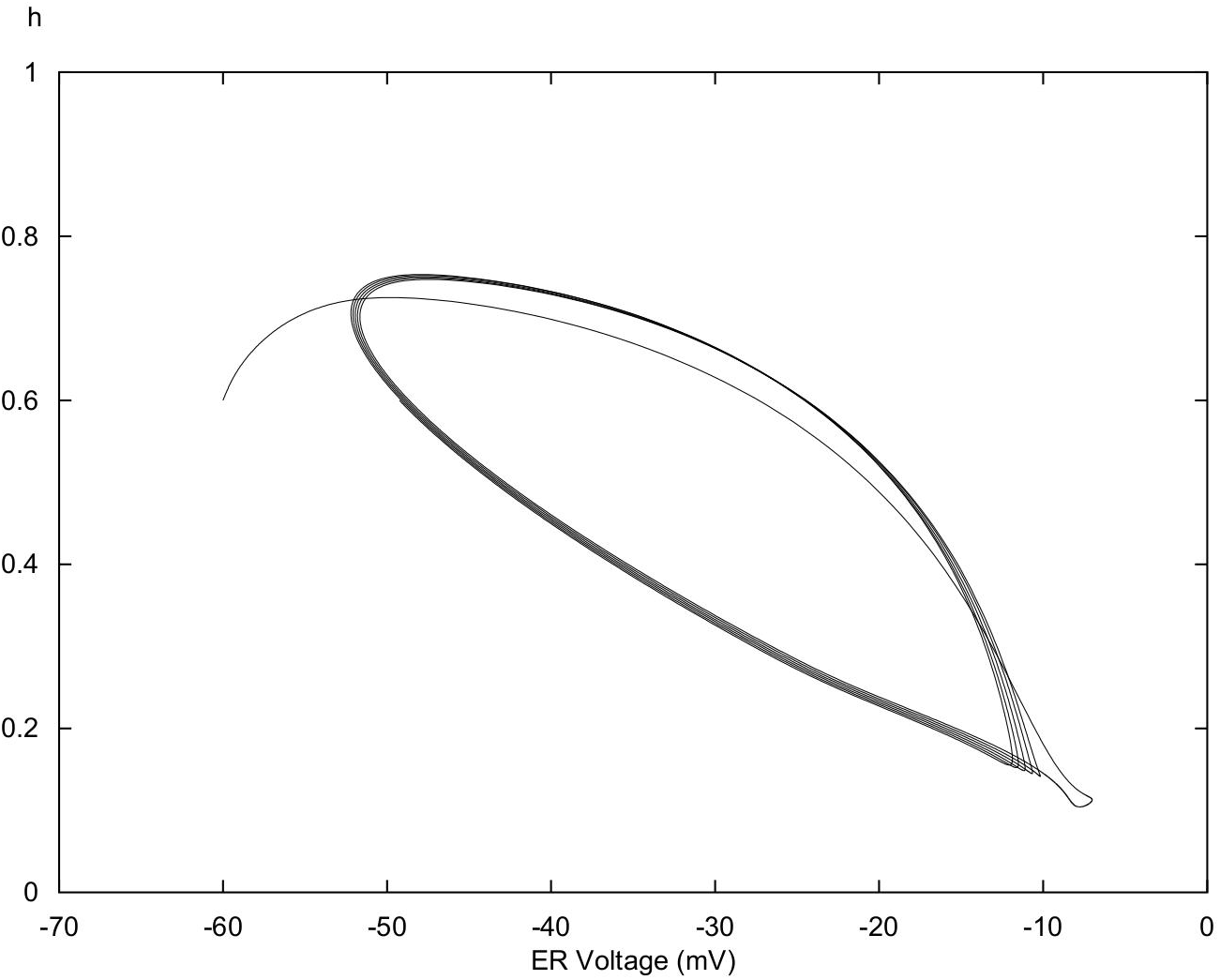
Phase portrait of a biphasic orbit of the ER membrane potential in an open cell. Equations 7 to 11 were integrated as described in the legend of Figure 7. The panel shows the evolution of the integration in the h, P plane. It can be observed that the trajectory goes from the initial values 0.6, −60 mV to the point 0.1, −5 mV. This is initially a stable point, but after a few minutes (Figure 7) the point is no longer stable and the trajectory continues to the limit cycle where it remains for the rest of the integration. During this second phase, the ER membrane potential oscillates between a minimum value of −50 mV and a maximum value of −10 mV.

### 3.3 Effect of thapsigargin

A second aspect to consider when evaluating a theoretical *β*-cell model is how the model cell responds to regulatory factors and pharmacological agents that have a well defined effect on pancreatic *β*-cells. Most significant among these are processes that affect the intracellular dynamics of calcium through their action on the SERCA pumps that transport Ca^2+^ into the ER or by modulating the conductance of the ER membrane calcium channels.

The ATP-dependent SERCA pumps that transport Ca^2+^ actively from the cytosol into the lumen of the ER play a central role in the process of insulin secretion in *β*-cells (212). A reduced SERCA pump activity has been observed in *β*-cells derived from animal models of diabetes such as the db/db mouse (213) and the Goto-Kakizaki rat (214). It is believed that the defective Ca^2+^ uptake may represent the primary cause for the development of T2DM in these animals (212, 213). In humans, it has been found that the SERCA3 locus possibly contributes to the genetic susceptibility to Type 2 diabetes (215).

An important tool in the study of the physiological role of SERCA pumps has been the irreversible inhibitor thapsigargin (216–218). In *β*-cells, thapsigargin causes calcium depletion of the ER and induces a nonselective cation current in the PM that depolarizes the membrane (166, 167). Glucose stimulation of thapsigargin-treated cells results in a sustained elevation of cytosolic calcium (23) and the continuous firing of action potentials (167, 219). The treated cells do not display the oscillations of [Ca^2+^]_*CYT*_ nor the electrical bursting observed in control untreated cells. In glucose-stimulated *β*-cells from db/db diabetic mice it has been observed also that the gradual decrease of SERCA pump activity is associated with the loss of periodic oscillations of cytosolic calcium (213).

The treatment with thapsigargin also affects the secretory response of *β*-cells. The short term effect is a potentiation of the stimulatory action of glucose, as can be expected from the activation of continuous firing. However, with prolonged exposures the cells lose the capacity to respond to glucose (213). Even, prolonged treatments with thapsigargin have been reported to induce apoptosis in *β*-cells (220).

I have analyzed the effect of a simulated exposure of the model *β*-cell to thapsigargin in both non-stimulated and glucose stimulated conditions.

The result of the treatment of a glucose-stimulated model cell is shown in Figure 9A. The control response of the cell without thapsigargin is shown in Figure 9B. In the conditions used in these simulations, low initial [Ca^2+^]_*ER*_, the control response consists only on periodic electrical bursting without the continuous initial phase shown in Figure 1 that requires a higher initial ER calcium concentration. The cytosolic concentration of calcium in the control cell oscillated in parallel with the electrical oscillations of the PM as shown in Figure 2A. By contrast, the response of the thapsigargin-treated cell was a continuous firing of action potentials (Figure 9A) accompanied by the depolarization of the ER membrane and a sustained elevation of cytosolic calcium (not shown). The pattern of the response to thapsigargin of the model cell in glucose-stimulated conditions is therefore the same observed experimentally in *β*-cells (23, 167).

**Figure 9:**
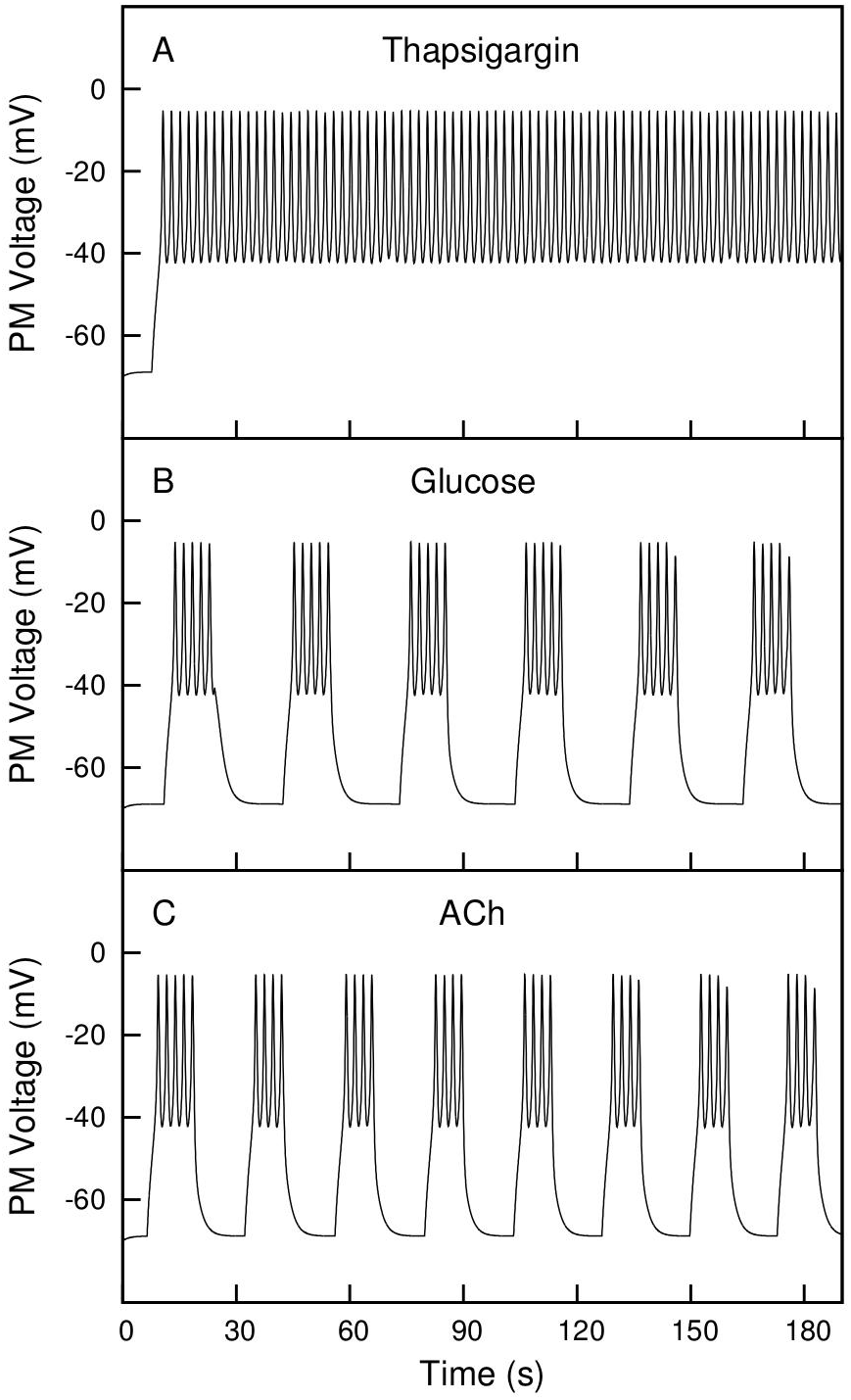
Effect of simulated Thapsigargin and ACh treatments. Equations 5 to 11 were integrated for 190 seconds, using the parameters shown in Table 1, unless otherwise indicated. The integrations were started from the initial conditions shown in Table 1, except for the concentration of calcium in the ER that was 600 *μ*mol l ^−1^. Panel B shows a simulation of normal control cells in glucose-stimulated conditions (g_*K(ATP)*_: 800 fS; v_*IP*3_: 2.8 *μ*mol l ^−1^ *s*^−1^). The cells displayed regular bursting with a frequency of 2 bursts per minute. Panel A shows the effect of reducing the rate of calcium uptake into the ER from 1.35 *μ*mol l ^−1^ *s*^−1^ to 1.0 *μ*mol l ^−1^ *s*^−1^ to simulate the treatment with the SERCA inhibitor thapsigargin. In these conditions the PM fired continuous action potentials. The simulation of ACh treatment is shown in panel C. The rate of IP3 synthesis was elevated from 2.8 *μ*mol l ^−1^ *s*^−1^ to 4 *μ*mol l ^−1^ *s*^−1^. The elevation of v_*IP*3_ caused an increase of the bursting frequency from 2 to 2.7 bursts per minute.

With parameter values corresponding to a cell growing in a low concentration of glucose, the simulated thapsigargin treatment caused the depolarization of the ER membrane and an elevation of the cytosolic concentration of calcium, but had no significant effect on the electrical activity of the plasma membrane that became slightly depolarized to a level, −56 mV, insufficient to trigger the firing of action potentials. This response agrees with the experimental observation that in the absence of glucose thapsigargin does not stimulate electrical activity nor insulin secretion in *β*-cells (167, 213).

The behavior of pancreatic *β*-cells with reduced SERCA activity, due either to the exposure to thapsigargin or to genetic defects as in the db/db diabetic mice, is consistent with the theoretical predictions of the stability analysis described in the previous section. The decreased SERCA activity in thapsigargin-treated cells, a reduction of the rate of calcium uptake v*_ER_*, displaces the state of the system to the left in the stability diagram in Figure 5, from the Oscillations area to the Depolarized area. This changes the functional program of the cell from displaying periodic bursting to the continuous firing of action potentials.

A similar leftward displacement has been observed during the progressive loss of SERCA activity in db/db mice. Roe et al. reported that as the pumping activity in *β*-cells decreases in the early weeks after birth, the biphasic response to glucose is triggered by increasingly lower concentrations of glucose. This result is also consistent with the stability diagram in Figure 6 that predicts that as v_*ER*_ decreases the required rate of IP3 production (v_*IP*3_) for the transition between a resting (Hyperpolarized) to an activated (Oscillations) state also decreases. What means that the triggering level of IP3 can be achieved at lower concentrations of glucose as experimentally observed.

An important finding in these experimental studies, outside the scope of the present theoretical study, is that the Depolarized area in the stability diagrams seems to be associated with the triggering of apoptosis and that this probably is what eventually causes the development of hyperglycemia and T2DM in these animal models of diabetes.

### 3.4 Regulation of the IP3 pathway

The IP3 signaling pathway plays a central role in the stimulus-response coupling in *β*-cells and also mediates the modulatory effects of incretins and neural factors such as GLP-1 and ACh (59, 221). I have analyzed the effect of simulating the glucose and neuro-hormonal activation of the IP3 pathway by raising the value of v_*IP*3_, the parameter that represents the rate of IP3 synthesis in the model *β*-cell.

In low-glucose conditions (open K_*ATP*_ channels and low IP3 synthesis), the conditions of pancreatic islets in fasted animals, the exposure of *β*-cells to either ACh (222, 223) or GLP-1 does not activate the secretion of insulin. To simulate these type of experiments, I integrated the model equations as described in the legend of Figure 9. First, the parameters *g*_*K(ATP)*_ and v_*IP*3_ were set to 1600 fS and 1.6 μmol l ^−1^ *s*^−1^, that correspond to non-stimulated cells in low-glucose conditions. With these parameter values, the plasma and ER membranes remained hyperpolarized during the whole integration with values of −70 mV and −54 mV respectively. This is the typical response of cells at rest. Elevating the value of v*_IP_*_3_ to simulate the activation of the IP3 pathway caused the appearance of oscillations of V*_ER_* the electric potential across the ER membrane, for values of v*_IP3_* of 2.2 *μ*mol 1^−1^ *s*^−1^and above. V*_ER_* oscillated between −55 mV and −10 mV, but these oscillations were unable to stimulate the firing of action potentials by the plasma membrane, the indication in the model of active insulin secretion. In simulated low-glucose conditions, the model cell is therefore as unresponsive to the activation of the IP3 pathway as pancreatic *β*-cells.

At moderate but still non-stimulatory concentrations of glucose, the conditions in blood shortly after food ingestion, the activation of the IP3 pathway does induce the secretion of insulin (224). To test this response in the model cell, I run a new series of simulations starting with parameter values, 1000 pS for *g_K(ATP)_* and 2.0 *μ*mol l ^−1^ *s*^−1^ for v*_IP3_*, that correspond to a higher but still non-stimulatory concentration of glucose. In these conditions both the ER and the plasma membranes remained hypepolarized at −48 mV and −70 mV during the whole integration. However, when the value of v_*IP*3_ was increased to 2.2 *μ*mol 1^−1^ *s*^−1^and above, to simulate the activation of the IP3 pathway at this level of glucose, the plasma membrane began firing action potentials indicating the activation of insulin release. This shows that the model cell has a threshold of K_*ATP*_ conductance below which it can not be induced to secrete insulin by activating the IP3 pathway. This seems to be also the situation with pancreatic *β*-cells. It has been proposed that the early action specially of the incretins shortly after food ingestion prevents that blood glucose reaches a significant elevation that could cause hyperglycemia.

At stimulatory levels of glucose, a further elevation of glucose concentration in the media or the stimulation of the *β*-cells with incretins or neural factors cause an increase of the frequency of bursting and the elevation of the rate of insulin secretion. In these conditions, the model also replicates the electrical response of *β*-cells, as can be seen in Figure 9. The elevation of the rate of IP3 synthesis from 2.8 *μ*mol l ^−1^ *s*^−1^, the level assumed to have been stimulated by glucose, to 4.0 *μ*mol 1 ^−1^ *s*^−1^, the new level reached by the addition of ACh, increases the frequency of bursting from 2 to 2.7 bursts per minute (compare panels B and C in Figure 9).

In the three conditions analyzed therefore the behavior of the model cell replicates the main features of the physiological response of *β*-cells to the parasympathetic and incretin hormone regulation of the IP3 signaling pathway.

### 3.5 A Model of Type 2 Diabetes

The two most common forms of diabetes are insulin-dependent diabetes mellitus (IDDM or type 1 diabetes) and non-insulin-dependent diabetes mellitus (NIDDM or type 2 diabetes). Type 1 diabetes is an autoimmune disease characterized by the destruction of the pancreatic *β*-cells (225, 226). Type 2 diabetes mellitus (T2DM) is caused initially by a secretory malfunction of *β*-cells that produces hyperglycemia and, as the disease progresses, loss of *β*-cell mass due to the activation of apoptosis and the consequent cell death (227, 228). A variety of pathological processes culminate in the development of type 2 diabetes (229, 230). In the most common, associated with obesity, the primary defect seems to be the development of insulin resistance in muscle, liver and other organs. The reduced uptake of glucose by these tissues causes a stark elevation of glucose in the blood that is usually compensated by an increased rate of insulin secretion. However, in some cases and for reasons still not well understood, *β*-cells fail to secrete enough insulin and the individual develops hyperglycemia. Chronic hyperglycemia is believed to cause the activation of apoptosis that compounds the initial secretory defects and precipitates the development of T2DM (231). In some instances though, in particular in non-obese patients, the primary cause of diabetes is the malfunction of the insulin secretory process itself, without any evidence of insulin resistance (232). The identification of the genetic and environmental factors that predispose *β*-cells to lose their functionality is of crucial importance for the diagnostic and treatment of T2DM. A great progress has been made recently. More than 80 genetic loci that confer T2DM susceptibility have been identified in genetic studies (233, 234), although it has been estimated that these mutant loci account for less than ten percent of the overall T2DM risk (235). Furthermore, several mechanisms have been proposed to mediate the activation of *β*-cell apoptosis in diabetic subjects. They include glucose toxicity (236), lipotoxicity (237), mitochondrial dysfunction (238), oxidative stress (239), islet amyloid polypeptide deposition (240–242) and ER stress (243, 244). However, all these mechanisms, perhaps with the exception of islet amyloidosis (245), are secondary to the failure of *β*-cells to compensate for the acute elevation of blood glucose. The primary cause of the insulin secretory defect that triggers the development of T2DM still remains elusive (246).

There are several clinical strategies to treat T2DM. One of the oldest is the use of sulfonyl ureas (SU) (247). These drugs bind to the regulatory subunit of the K*_ATP_* channel (SUR1) and close the channel. As a consequence, *β*-cells become electrically active and secrete insulin independently of glucose. In this way, the treatment compensates the secretory defect in T2DM patients. An important problem with this approach is that the induction of insulin secretion takes place even at low levels of blood glucose and can cause severe hypoglycemia if not properly controlled (248).

Alternative to the use of SU drugs are strategies aimed at the activation of incretin receptor pathways, by means of GIP or GLP-1 analogs (249, 250) or by using inhibitors of the DPP-4 protease that degrades GIP and GLP-1 (251). As indicated in the previous section, the activation of the GLP-1 receptor only potentiates insulin secretion in the presence of elevated glucose, therefore this approach does not have the risk of causing hypoglycemia. Furthermore, the treatment also stimulates *β*-cell proliferation thus compensating for the loss of *β*-cell mass associated with T2DM (252, 253). The negative effect of this type of treatments is that the chronic activation of GLP-1 receptors leads in some cases to the development of pancreatic and thyroid cancer (254).

I have used the mathematical model to explore the potential role of the ER in the genesis of T2DM and to see the effects on the model *β*-cell of simulating the therapeutic treatments described above.

In the model, the electrical activity of the ER membrane plays a major role in the generation of the secretory response of *β*-cells. Its malfunction is therefore likely to have an effect comparable to the secretory failure in T2DM *β*-cells. To test the idea, I have simulated the ER malfunction by preventing the elevation of v*_IP3_*, the rate of IP3 synthesis, when the model cell is exposed to glucose. The result of this simulation is shown in Figure 10. In panel A, it can be observed the control response of a normal cell to a simulated elevation of glucose. The numerical integration reproduced the characteristic biphasic response previously shown in Figure 1. Panel B shows the result of keeping v*_IP3_* at the non-stimulated value of 1.6 *μ*mol l ^−1^ *s*^−1^ to mimic a defective response from the ER. In these conditions, the glucose-induced closing of the K_*ATP*_ channels (*g*_*K(ATP)*_ reduced from 1600 fS to 800 fS) was unable to depolarize the PM to the threshold required to trigger the ring of action potentials. Even at more extreme conditions (*g*_*K(ATP)*_ set to 400 fS), simulating a higher elevation of glucose in the growing media, the PM remained hyperpolarized (V_PM_: −60 mV) during the whole integration. The model cell with a defective ER response behaves therefore like a defective T2DM *β*-cell unable to secrete insulin in response to an elevation of glucose.

**Figure 10:**
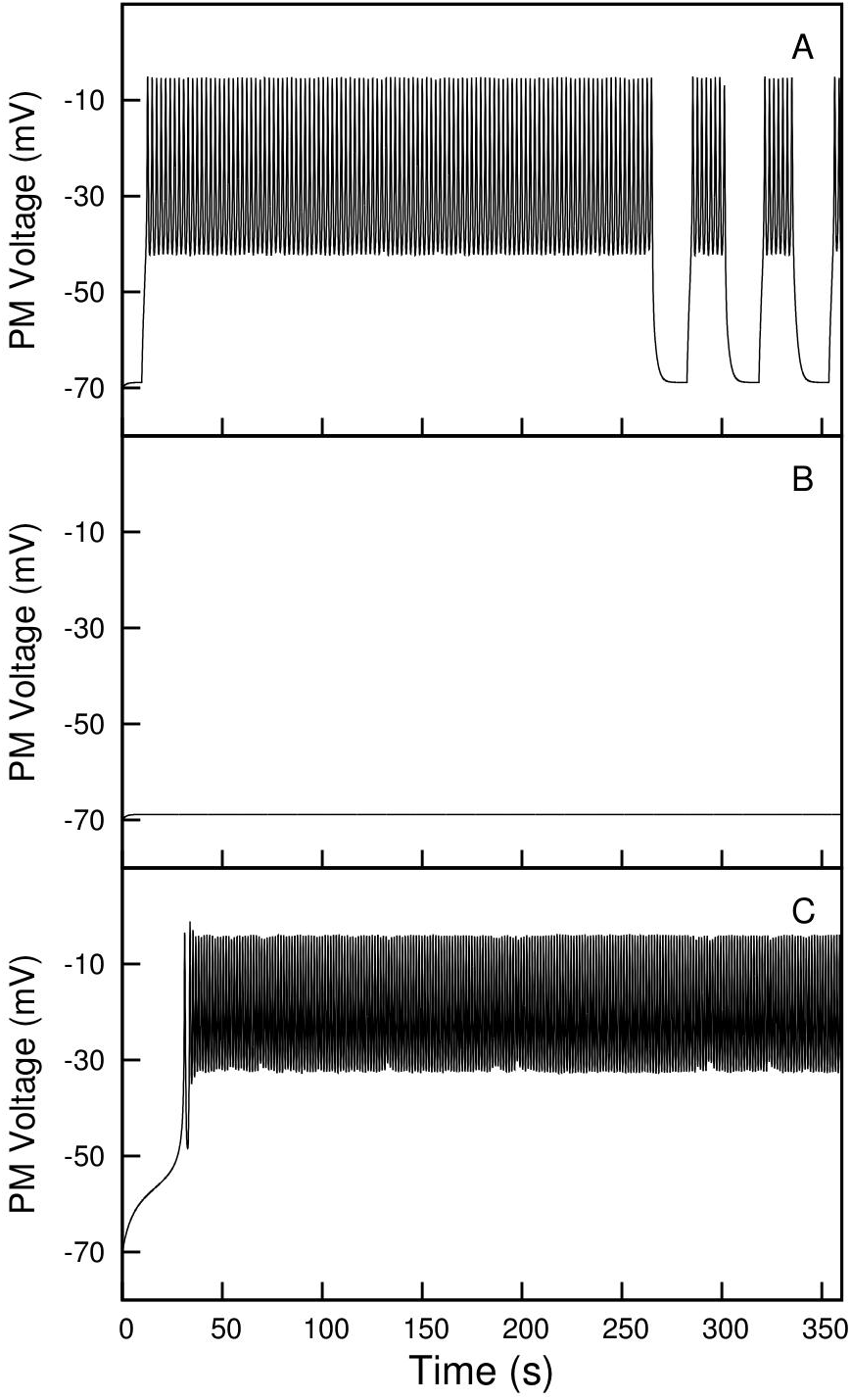
0: A model of Type 2 Diabetes. Response to glucose of a normal cell (Panel A), a cell with a defective ER signaling system (Panel B) and a defective cell treated with sulphonylurea (Panel C). Equations 5 to 11 were numerically integrated for 360 seconds, starting from the initial conditions in Table 2 and with the parameter values shown in Table 1 except otherwise indicated. Panel A shows the result of the integration in conditions simulating the glucose activation of a normal cell. ( v_*IP*3_: 2.8*μ*mol l ^−1^ *s*^−1^ and *g*_*K(ATP)*_: 800 fS.) A normal biphasic response like the one shown in figure 1 can be observed. In the simulation in panel B, the parameter v_*IP*3_ was reduced to 1.6 *μ*mol l ^−1^ *s*^−1^ while keeping *g*_*K(ATP)*_ at 800 fS to simulate the exposure to glucose of a defective cell unable to trigger the electrical response of the ER. With these parameter values, the electric potential of the plasma membrane remained at −70 mV during the whole integration. The simulated defective cell, like T2DM *β*-cells, was unresponsive to glucose. Panel C shows the reactivation of the secretory activity in the diabetic-like model cell by simulating a sulfonylurea treatment. The parameter v_*IP*3_ was maintained at the non-functional value of 1.6 *μ*mol 1^−1^ *s*^−1^ and the K_*ATP*_ conductance was reduced to 360 fS to simulate the inhibitory effect of SU on the K_*ATP*_ channels. The integration with these parameter values caused the continuous firing of action potentials, as has been reported for SU-treated *β*-cells. The treatment with an incretin analog of a glucose-exposed defective cell can be simulated by elevating the value of v_IP3_, the rate of IP3 synthesis, above the basal level of 1.6 *μ*mol l ^−1^ *s*^−1^ to reflect the activation of the IP3 pathway by this type of drugs. A simulation with a v_*IP*3_ value of 2.8*μ*mol l ^−1^ *s*^−1^ and a conductance *g*_*K(ATP)*_ equal to 800 fS produces the same result shown in Panel A, a result that was obtained integrating with exactly these parameters. The family of incretin-related drugs therefore restores a normal secretory activity in the diabetic-like model cell, as they do in type 2 diabetic *β*-cells.

In the model, the electrical activity of the ER membrane plays a major role in the generation of the secretory response of *β*-cells. Its malfunction is therefore likely to have an effect comparable to the secretory failure in T2DM *β*-cells. To test the idea, I have simulated the ER malfunction by preventing the elevation of v_*IP*3_, the rate of IP3 synthesis, when the model cell is exposed to glucose. The result of this simulation is shown in Figure 10. In panel A, it can be observed the control response of a normal cell to a simulated elevation of glucose. The numerical integration reproduced the characteristic biphasic response previously shown in Figure 1. Panel B shows the result of keeping v_*IP*3_ at the non-stimulated value of 1.6 *μ*mol l ^−1^ *s*^−1^ to mimic a defective response from the ER. In these conditions, the glucose-induced closing of the K_*ATP*_ channels (*g_K(ATP)_* reduced from 1600 fS to 800 fS) was unable to depolarize the PM to the threshold required to trigger the firing of action potentials. Even at more extreme conditions (*g_K(ATP)_* set to 400 fS), simulating a higher elevation of glucose in the growing media, the PM remained hyperpolarized (V*_PM_*: −60 mV) during the whole integration. The model cell with a defective ER response behaves therefore like a defective T2DM *β*-cell unable to secrete insulin in response to an elevation of glucose.

Sulfonylureas restore the secretory activity in diabetic *β*-cells by closing the K*_ATP_* channels. This effect can be emulated in the diabetic-like model cell by reducing the value of *g_K(ATP)_*. Figure 10C shows the result of a simulation in which *g_K(ATP)_* was lowered to 360 fS while keeping v*_IP_*_3_ at the low inactivated value of 1.6*μ*mol l ^−1^ *s*^−1^. In these conditions, the PM displayed a continuous firing of action potentials. A simulated SU treatment therefore is able to restore the secretory capacity of the model cell too.

To simulate the effect of incretins in the diabetic-like model cell, the value of v*_IP_*_3_ should be elevated to mimic the activation of IP3 synthesis caused by this type of drugs. A simulation with elevated v*_IP_*_3_ (2.8 *μ*mol l^−1^ *s*^−1^) and reduced K_*ATP*_ conductance (800 fS) produces the same response shown in Figure 10A, as the model does not discriminate whether the v*_IP_*_3_ elevation is caused by glucose or by GLP-1 receptor activation. So, as with sulfonylureas, the simulation of a treatment with a GLP-1 analog also restores the secretory activity in the unresponsive, diabetic-like model cell.

These results therefore are consistent with the hypothesis that a primary cause for the development of T2DM may be the malfunction of the electrical response of the ER membrane.

## 4 Discussion

In this paper I present a minimal mathematical model of an insulin secreting pancreatic *β*-cell. The model cell contains an excitable ER membrane that modulates the electrical activity of the PM. The interaction of these two excitable membranes, ER and PM, generates an electrical response (Figure 1) comparable to the response observed in glucose stimulated *β*-cells (9, 10, 15, 120). In both, the theoretical model and live *β*-cells, the response is biphasic with an initial phase characterized by continuous firing followed by rhythmic bursts of action potentials in the second pulsatile phase.

### 4.1 Dynamic states of the ER membrane

The main assumption on which this model is built is that the electrical activity of the ER membrane is the driving force that determines the nature of the PM response in stimulated *β*-cells. Consequently, to understand how the continuous and rhythmic patterns of action potential firing are generated requires to investigate the causes of the complex electrical behavior of the ER membrane in the model cell.

The study of the intracellular dynamics of calcium has already established that the ER constitutes a nonlinear dynamical system, what Li and Rinzel have called the calcium excitability of the ER (193, 198, 255). This nonlinearity, due fundamentally to the biophysical properties of the IP3 receptor (44, 46), is responsible for the complex behavior of calcium in eukaryotic cells. A behavior that includes cytosolic oscillations of calcium concentration and propagating calcium waves. As described in the Introduction, the electrical activity of the ER membrane is closely linked to the dynamics of intracellular calcium and is therefore equally nonlinear.

A well known characteristic of nonlinear systems is that small changes in the value of some parameter may cause dramatic alterations in the behavior of the system (256). In pancreatic *β*-cells, two parameters, v*_ER_*, the maximal velocity of the SERCA pump, and v*_IP_*_3_, the rate of IP3 synthesis, are controlled by the main signaling systems that convey the effects of nutrients and regulatory factors to the secretory activity of the cells. This makes them good candidates to analyze how changes in stimulatory conditions affect the dynamics of the ER.

A result of this analysis has been the identification of three different dynamic states of the ER membrane that explain how the biphasic electrical response of the PM and the pattern of electrical bursting are generated.

I started the analysis in a closed model cell with an inert plasma membrane. In this reduced model, the variation of v*_ER_* and v*_IP_*_3_ within a physiological range of values uncovered three different behaviors of the ER membrane (Figures 4 and 5). For a set of parameter values, the ER membrane remained hyperpolarized reflecting a non-stimulated, resting cell (Figure 4C). The behavior is caused by the evolution of the system to a steady state determined by a stable fixed point. A second set of values caused the membrane to settle in a depolarized state (Figure 4A), indicating the presence of a second stable fixed point in the dynamic system. This second steady state corresponds in the whole cell model to a hyperactive cell that fires continuous action potentials as in the first phase of the response to glucose (Figure 1). Finally, in an area of the parameter map a new dynamic state emerges, a limit cycle, responsible for an oscillatory behavior of the ER membrane potential (Figure 4B). In the whole cell model, the oscillatory state of the ER membrane generates the electrical bursting seen in the second phase of the response of the cell to glucose (Figure 1).

The closed cell model explains how the two firing patterns of the biphasic response are generated but it does not provide an explanation for the transition between the two phases. To understand how this transition takes place, it is necessary to study the behavior of the ER membrane in an open cell model.

The electric potential of the ER membrane in a still non-excitable but open cell displays the same three behaviors observed in the closed cell, hyperpolarization, depolarization and cyclic oscillations. However, the stability diagram of the open cell (Figure 6) is not static as in the closed model but varies with the changing levels of intracellular calcium. As the cell loses calcium, the stability diagram suffers a leftward displacement. Parameter points that at the beginning of the simulation reside in a stable depolarized area become at some time point part of the oscillatory area. This displacement or change of the stability conditions of the system is what causes the transition from the initial continuous firing phase to the second bursting phase in the whole *β*-cell model.

The nonlinearity or excitability of the ER provides therefore a unified explanation for the whole electrical response of *β*-cells to glucose, the continuous firing of the initial phase and the pulsatile firing of the bursting phase, as well as the mechanism that causes the transition from one phase to the other.

A second important insight gained with this analysis is the correlation of the dynamic states of the ER and the long-term fate of *β*-cells. Each dynamic state seems associated with a different developmental program, quiescent cells, proliferating cells or apoptosis.

The hyperpolarized state corresponds with non-stimulated cells, cells with no secretory activity living in quiescent-like conditions. Cells in this state have a considerable long-term stability with low rates of cell division and cell death.

The state with a depolarized ER membrane corresponds to hyperactive cells with an acute level of insulin secretion. This is the normal state of *β*-cells for a brief period of time after the elevation of blood glucose when the cells fire a continuous train of action potentials. In some conditions though, such as the prolonged exposure to very high glucose or when the SERCA pump has low activity due to the action of inhibitors or to genetic defects as in diabetic animal models (257), this hyperactive state remains for long time intervals. In all these situations, a persistent depolarized state is associated with the activation of apoptosis and cell death (258).

The oscillatory state of the ER membrane potential corresponds with cells displaying electrical bursting. Electrical bursting is induced by moderate elevations of glucose in the blood stream, usually in collaboration with incretins like GLP-1 secreted by the intestine and by ACh released in the pancreas by parasympathetic nerve terminals. The activation of the bursting state has been associated with the induction of cell proliferation. In fact, part of the therapeutic effect of incretin analogs is attributed to the activation of *β*-cell proliferation that compensates for the loss of cell mass that takes place in type 2 diabetes (75, 253).

The correlation of the dynamic states of the ER membrane with particular functional and developmental programs suggests a key role for the ER as a central processor that integrates all environmental information, in the context of a genetic background, and adopts a dynamic state that determines the immediate functional response of the cell as well as long-term effects like cell proliferation, cell differentiation or cell death.

### 4.2 On the Origin of Type 2 Diabetes Mellitus

In spite of the extensive investigation on the causes of type 2 diabetes, the primary events that trigger the progressive degradation of *β*-cells that ends up in T2DM have not been elucidated yet. Using the mathematical model presented here, I have been able to reproduce the phenotype of T2DM *β*-cells by simulating a cell with a defective ER response. This diabetic-like model cell is unable to generate a normal electrical response to a simulated elevation of glucose, an indication that it has lost the normal secretory activity (Figure 10 B). Interestingly, a simulated exposure of the diabetic model cell to the standard drugs used to treat T2DM, sulphonyl ureas and incretin analogs, restores the electrical response of the cell and thus its ability to secrete insulin (Figure 10 A and C). These results suggest that in actual *β*-cells a malfunction of the ER response may indeed be one of the earliest events that lead to type 2 diabetes.

Some evidence from animal models of diabetes supports this point of view (259, 260). In both, obese (213, 261) and non-obese (214, 257, 262) diabetic rodents, it has been observed that the progression of the diabetic state is accompanied by a gradual loss of SERCA pump activity and thus a defective electrical response of the ER. The reduced SERCA activity initially causes insulin hypersecretion (257) as would be expected from the stability diagram in Figure 6, but eventually the damage on the ER disrupts the secretory process causing hyperglycemia and the development of diabetes.

Genetic analysis in humans also provides support for the hypothesis. Some variants of the SERCA and TRPM5 loci, two genes involved in the ER-dependent response to glucose, have been found associated with an increased risk of developing type 2 diabetes (118, 215).

An indirect evidence for the involvement of a defective ER in the development of T2DM in humans is the therapeutic effectiveness of sulphonyl ureas and incretin analogs in most patients with T2DM. SUs bind to the same two sites occupied by ATP in the SUR1 subunit of the K_*ATP*_ channel (247). The reactivation of insulin secretion by these drugs therefore indicates the presence of a fully functional K_*ATP*_ channel in T2DM *β*-cells. The molecular defect must reside on one of the alternative pathways activated by glucose. That this pathway most likely involves the ER is suggested by the successful restoration of normal insulin secretion by incretin analogs in most T2DM patients. These drugs elevate IP3 independently of glucose and thus can compensate for a defective ER activation by the sugar in diabetic cells (249, 250).

An explanation of type 2 diabetes in terms of the malfunction of the electrical response of the ER, a nonlinear system in which small changes in a multitude of parameters can alter dramatically its behavior, would explain the difficulty to single out critical genetic and environmental factors that confer susceptibility to the disease.

An impressive amount of work has been dedicated to investigate the genetic basis of T2DM (263–265). In genome-wide association studies, more than 80 common variants associated with T2DM have been identified (234, 264, 266, 267). Most of the identified loci though are located at intergenic or intronic regions, making it difficult to study its functional role (268). Furthermore, each variant has a relative low effect and together account for only a small percentage of T2DM heritability (234, 268, 269). These limitations have motivated and exhaustive search for rare or low frequency variants with larger functional effects. With this approach, several genes whose loss of function is associated with T2DM have been identified (270–272). Unfortunately, some of these genes are involved in functions, such as melatonin signaling (270) or adipocyte differentiation (271), that are difficult to correlate with the development of diabetes.

The initial expectations that the genetic analysis of T2DM would clarify the etiology of the disease and produce genetic markers for risk prediction and early diagnosis have so far failed to materialize. The genetic analysis would benefit from a strategy that incorporates the study of the electrical activity of the ER in *β*-cells from healthy persons and from individuals that carry the genetic variants that predispose to develop diabetes. This approach may unravel the actual role the susceptibility genes play in *β*-cells and improve the understanding of the sequence of events that lead to T2DM. The identification of ER abnormalities at the initial stages of type 2 diabetes may also provide an effective tool for early diagnosis and risk assessment.

### 4.3 Physiological role of the ER membrane potential

What distinguishes this model from previous ones is the change of the slow variable assumed to drive electrical bursting. While in previous models this slow variable was the intracellular concentration of calcium, either in the cytosol ([Ca^2+^]_*CYT*_) or in the lumen of the ER ([Ca^2+^]_*ER*_), here it is postulated to be the electric potential of the ER membrane, V_*ER*_. With this change of variable, the model is able to explain the electrical response of pancreatic *β*-cells to glucose, including the generation of electrical bursting, and even provides a potential explanation for the development of type 2 diabetes.

Although the hypothesis is successful at theoretical level, its relevance needs to be tested in actual cells. It is thus important to evaluate what is the evidence hinting to the presence in *β*-cells of V_*ER*
_-regulated PM channels activated by the depolarization of the ER membrane as proposed in the model.

A good candidate for a PM channel regulated by the depolarization of the ER membrane is TRPM5. As indicated in the Introduction, TRPM5 is a store regulated cation channel that is expressed in the PM of *β*-cells. The channel is required for the normal secretion of insulin. Mutant *β*-cells lacking TRPM5 have a defective secretion of insulin and are unable to generate electrical bursting (114, 115).

In the initial studies on TRPM5, it was found that the channel was activated by the IP3- dependent release of calcium from the ER (110, 273) although the activation did not seem to be caused by the depletion of lumenal calcium (113, 274, 275). It was therefore proposed that TRPM5 was activated by the elevation of the cytosolic concentration of calcium (112). However, the experimental evidence is not fully consistent with this hypothesis. In patch-clamp studies with isolated channels it has been found that TRPM5 is indeed opened by calcium but at concentrations of the order of 50 *μ*M, way above the usual values found in the cytosol (111, 275). At physiological concentrations of calcium (less than 1 *μ*M), the channel was completely unresponsive in this recording configuration (274, 276). Only when a whole cell recording mode is used does TRPM5 appear regulated by physiological levels of calcium (113, 276). Interestingly, in this whole cell configuration, the activation of the channel has the same bell-shaped dependence on calcium that the IP3R (113, 114), as if the opening of the channel depended on the current of calcium through IP3Rs rather than on the cytosolic concentration of calcium.

In another set of experiments, Zhang et al. (273) found that the elevation of cytosolic calcium using the SERCA pump inhibitor thapsigargin did not activate the TRPM5 channel whereas, conversely, the activation of the release of calcium from the ER while preventing the elevation of its concentration in the cytosol did cause the opening of the channel. Furthermore, Prawitt et al. (113) have reported that the activation of TRPM5 depends on the rate of calcium release from the ER rather than on the actual concentration of calcium in the cytosol. All these results could be better explained by a mechanism of regulation of TRPM5 that depends on the depolarization of the ER membrane instead of on the concentration of cytosolic calcium.

TRPM5 and perhaps other members of the TRP channel family such as TRPM4 and TRPC2 may therefore provide the molecular basis for the hypothetical V_*ER*
_-regulated PM channel proposed in this theoretical study. The loss of electrical bursting in Trpm5 −/− cells (114) is consistent with TRPM5 playing in pancreatic *β*-cells the same role played by the V_*ER*_-regulated channel in the model cell.

The electric potential across the ER membrane may indeed be involved in the regulation of insulin secretion in pancreatic *β*-cells. But the regulatory role of V_*ER*_ is unlikely to be restricted to just this type of cells. A wide variety of physiological processes, from fertilization to cell death, display a similarly complex dynamics of calcium (185, 277) and are thus candidates to be regulated by the electrical activity of the ER membrane.

During fertilization, the transition of the oocyte from a quiescent state, arrested metaphase of the second meiotic division in mammals, to a state of mitotic proliferation is associated in all animal species studied with waves of Ca^2+^ that transverse the egg’s cytosol (278–281). The oscillations of calcium are essential for the initiation of embryogenesis. Experimental manipulations that prevent the oscillations also block the activation of the egg (282, 283). The calcium waves are triggered in mammals by the elevation of IP3 by the sperm-specific phospholipase C zeta (PLCζ) (284, 285). The injection of (PLCζ) in the oocyte is sufficient to induce the same calcium oscillations seen in fertilization and to activate the early steps of embryonic development (286, 287).

Although fertilization is always triggered by oscillations of calcium in the oocyte, the pattern of the calcium waves varies considerably between species and genera. In mammals, the oscillations consists in an initial transient that lasts several minutes followed by regular spikes with a frequency of the order of 5 to 10 per hour (285, 288). In amphibians and fish, a single wave transverses the egg from the animal to the vegetal hemisphere (280, 289, 290). In insects, in particular in Drosophila, there is a single rapid transient wave that extends usually from the posterior to the anterior pole of the egg (291). All these different patterns accomplish a common set of transformations in the egg that include resumption of meiosis, cortical granule exocytosis, activation of maternal mRNA translation, and cytoskeletal rearrangements (292, 293). The search for the mechanisms that link the oscillations of calcium with all these cellular transformations is a very active area of research.

The subsequent embryonic development is also associated with complex oscillations of cytosolic calcium (294, 295). The dynamics of calcium plays a central role in the key steps that lead to the formation of the embryo, including axis determination, the establishment of the germ layer, and organogenesis (296–298). In at least one case, during the development of the chick retina, the oscillations of calcium in the embryo have been found associated with parallel oscillations of the electric potential across the ER membrane (178).

Another example of a physiological process controlled by the dynamics of calcium is provided by the immune system. The antigen-dependent activation of T lymphocytes, a process that involves the control of gene expression, cell proliferation and cell differentiation, depends on several signaling pathways (299), the most important being the activation of PLC and the consequent IP3-dependent release of calcium from the ER. The release causes oscillations of cytosolic calcium (300, 301) and the activation of store-operated currents in the plasma membrane of the lymphocyte (302). The activation of B lymphocytes also depends on calcium signals (303). Interestingly, the particular dynamics of calcium, single transients, repetitive oscillations or sustained plateau, determines the type of transcriptional factor activated in both T and B lymphocytes (304–306). The central role of the calcium signaling in the immune response is emphasized by the fact that a defective calcium response in either T or B lymphocytes causes immunosuppression (303, 307).

In the nervous system, learning and memory are tightly linked to the dynamics of calcium (184, 308, 309). Pioneer studies with simple invertebrate models of learning provided the first evidence that the cAMP pathway, and thus the release of calcium from the ER, is involved in synaptic plasticity. For instance, the activation of the cAMP pathway in Aplysia by serotonin secreting interneurons was found to potentiate the secretion of neurotransmitter by stimulus activated sensory neurons (310) in much the same way that GLP-1 and other incretins potentiate the secretion of insulin in glucose-stimulated pancreatic *β*-cells (221). Genetic studies with Drosophila showed the involvement of the cAMP signaling pathway in learning and memory also in this organism (311, 312).

In higher vertebrates, learning and memory is assumed to depend on structural and functional modifications of dendritic spines (313) as originally proposed by Ramón y Cajal in the late nineteenth century (314). Dendritic spines are mushroom-like structures protruding from the dendritic shafts of neurons that serve as the anchoring point of presynaptic axonic terminals in most excitatory synapses (315). Inside the spines there is a projection of the ER that seems to play an important role in synaptic plasticity (316–319). When spines are activated by stimulating the presynaptic terminal with trains of electrical pulses, they experience modifications that include changes of size and number as well as variations of synaptic strength known as long-term potentiation (LTP) and long-term depression (LTD) (320–322). The primary event that triggers synaptic modifications in these experimental conditions is the elevation of calcium inside the dendritic spines. Calcium enters the spines through the N-methyl D aspartate (NMDA) type of glutamate receptor (323) and is also released from the projections of the ER inside the spines. The internal release is induced by calcium itself and by IP3 produced by activated metabotropic receptors (324, 325). Low elevations of calcium are believed to cause LTD whereas high intra-spinal concentrations seem to activate LTP (323). Interestingly, the elevation of calcium in the spines generates calcium waves that propagate through the dendritic shaft in both directions, reaching the nucleus on the soma side. These waves are believed to be involved in the reinforcement of Hebbian as well as hetorosynaptic plasticity (326, 327).

Inhibitory GABAergic synapses also display experience-dependent plasticity (I-LTP and I-LTD) that is involved in learning and memory (328, 329). This form of synaptic plasticity is usually heterosynaptic, requiring the activation of nearby glutamatergic excitatory synapses. A variety of regulatory mechanisms both pre- and postsynaptic have been observed (328). In many of them, such as for instance the endocannabinoid-mediated induction of I-LTD (330) or the induction of I-LTP by the brain-derived neurotrophic factor (BDNF) neurotophin (331), the regulation has been shown to be mediated by cAMP or IP3, indicating the involvement of the ER in the process.

This is only a limited sample of the many physiological situations in which the dynamics of calcium, and consequently of the electrical activity of the ER membrane, have been found to determine the normal physiological response and developmental fate of cells.

In addition to the regulatory activity in normal cells, the dynamics of the ER seems to play a pivotal role in the development of several pathological conditions as already discussed with respect to type 2 diabetes.

### 4.4 The ER membrane potential in Alzheimer’s disease

A pathological process with a parallelism with type 2 diabetes is Alzheimer’s Disease (AD), a neurodegenerative disorder that causes loss of memory and senile dementia (332). AD is characterized by synapse loss, predominantly in associative neocortical structures of the brain, the extracellular deposition of senile plaques composed mainly of misfolded amyloid-beta (A*β*) peptides, and the intracellular accumulation of hyper-phosphorilated tau protein, a component of the cytoskeleton (332, 333). The accumulation of A*β* deposits in the brain is caused by the disruption of the proteolytic cleavage of the amyloid precursor protein (APP) (334). This process is reminiscent of the extracellular accumulation of deposits of *β* amyloid fibrils in the islets of Langerhans of the pancreas of T2DM individuals, caused by the misfolding of the human islet amyloid polypeptide (hiAPP) in diabetic *β*-cells (242).

An explanation for the origin of AD, known as the amyloid hypothesis (335), proposes that the accumulation of insoluble A*β* aggregates in senile plaques represents the primary influence that drives AD pathogenesis (336, 337). The hypothesis overlooks the fact that senile plaques are often found in the brain of elderly individuals with no cognitive disorder and also that strategies that appear to be effective in removing the amyloid deposits have so far failed to produce any significant improvement in treated subjects (332, 338).

Competing or complementing the amyloid explanation is the calcium hypothesis, according to which the dysregulation of calcium signaling in dendritic spines is the primary cause of synaptic loss and neural degeneration in AD brains (339–341). Several lines of evidence support this point of view.

On the first place, the study of familial Alzheimers (FAD) has provided important insights on the role of calcium dynamics on the early development of AD. FAD is a rare form of Alzheimers caused by mutations on the genes encoding presenilin-1 (PSEN1), presenilin-2 (PSEN2) or amyloid precursor protein (APP) (342). The other forms of AD, known as sporadic or late-onset Alzheimers, also have a strong genetic component with more than 20 AD-related genes identified so far (343, 344) but their exact etiology remains unknown.

Presenilins (PSs) form the catalitic subunit of *γ*-secretase, one of the proteases involved in the generation of the toxic A*β* peptide from APP. The FAD mutations in PSs cause an increased production of A*β* that potentiates the toxic effects of the peptide, a mechanism consistent with the amyloid hypothesis (345). But in addition, presenilins have a second role in the ER as a Ca^2+^-leak channel (346, 347). Many FAD mutations in PS1 and PS2 cause the loss of this channel function. The phenotype of these mutations is an elevated [Ca^2+^]_*ER*_ that causes an abnormally high release of calcium from the ER when RyR or IP3R are activated (346, 348, 349).

Another effect of FAD mutations that also contributes to potentiate the release of calcium from the ER is the upregulation of the expression and activity of RyR2 (350, 351) and IP3R (341, 352). Interestingly, upregulation of RyR2 has also been observed in individuals with mild cognitive impairment, a condition considered a precursor of AD (353).

FAD mutations also affect the intracellular dynamics of calcium by reducing the activity of the SERCA pump (354). A reduction that has also been observed in *β*-cells from animal models of type 2 diabetes (213), and that increases the electrical excitability of the ER membrane (see Figure 6).

The balance of all the effects of mutated presenilins on calcium channels and pumps is an enhanced release of calcium from the ER that is likely to produce a sustained depolarization of the ER membrane, a condition that, as discussed in Section 4.1, is associated with an increased rate of apoptosis.

In addition to the effects on the ER, FAD mutations also cause changes in the PM. In particular, they reduce the level of store operated currents normally activated by the release of calcium from the ER (341, 355, 356), apparently by downregulating the activity of STIM2 (357, 358) and of two members of the TRP family, TRPC3 and TRPC6 (359). The reduction of the store operated inward currents are likely to have significant effects on the excitability of the postsynaptic PM.

A second line of evidence for the involvement of the ER in the development of AD comes from studies on the GLP-1 signaling pathway in the central nervous system (340, 360). The use of analogs of GLP-1, the same analogs used to treat type 2 diabetes (361), has been shown to reduce plaque deposition and neurological deterioration in patients with AD as well as in animal models of the disease (362–365). In fact, a connection between the pathophysiology of Alzheimers and type 2 diabetes has been proposed (366–368). The connection could well be a similar electrical malfunction of the ER. A malfunction that seems to be corrected, at least in part, in both *β*-cells and neurons by the activation of IP3-dependent Ca^2+^ release from the ER through the pharmacological stimulation of the GLP-1 signaling pathway.

The study of the familial form of Alzheimer’s disease has revealed that changes in the electrical excitability of the ER membrane and PM, as well as conditions that alter the normal electrical interactions between them, can be one of the primary causes of this particular form of the disease. Whether the same mechanisms also play a fundamental role in the onset of the sporadic forms of Alzheimer’s disease is a question that may be worth investigating.

A hallmark of the malfunction of the ER seems to be a significant hereditary component that nevertheless results very difficult to identify in genetic analysis. A pathology with these characteristics is Autism Spectrum Disorder (ASD), a series of neurodevelopmental abnormalities that cause dramatic behavioral changes in the persons affected. In concordance studies with identical twins and genetic studies in affected families, it has been established that ASD has a strong genetic component (369, 370). However, in common with type 2 diabetes, the elevated number of loci found associated with ASD susceptibility, more than 500 (371–373), has not so far provided meaningful information on the origin of the disease nor a genetic marker for the early diagnosis of the disorder.

Due to the neurodevelopmental nature of ASD, involving the maturation of neural circuits during prenatal and early postnatal development, it is believed that the diffierent forms of autism are caused by defects in the synaptogenesis and synaptic trimming that take place in these formative period, particularly in the cerebelum (374, 375). These are processes that depend critically on the dynamics of calcium in the synaptic terminals and dendritic spines (376), raising the possibility that an important factor in the genesis of autism related disorders may be a defective electrical activity of the ER.

### 4.5 Concluding Remarks

The theoretical analysis I have presented here illustrates the capacity of a nonlinear system to generate complex cellular behaviors. Even in the most simplified version of the model, the ER membrane displays three different dynamic states that generate three different functional programs in the cell. In actual cells, where the number of parameters and variables susceptible to be modified by environmental and genetic signals is orders of magnitude higher than in the minimal models studied here, the electrical activity of the ER is likely to be able to adopt a multiplicity of dynamic substates, each with a particular spatio-temporal pattern of oscillations, each encoding a particular functional program.

The progress in superresolution (SR) imaging technology has revealed the ER as a very complex and dynamic organelle, with an extremely sophisticated ultrastructure formed by cisternae and interconnected tubular matrices and tubular network arrays experiencing rapid transformations that reflect at the morphological level the rich variety of substates the organelle can adopt (182, 377).

Furthermore, the ER membrane interacts with practically all subcellular compartments in the cell from the PM to the nucleus. The ER gets in close apposition with the PM in areas known as ER-PM junctions (378, 379). This proximity seems necessary for the regulation of the conductance of store operated channels such as Orai and members of the TRP family and thus for the control of the excitability of the PM by the ER.

The ER interacts with mitochondria in specialized areas known as mitochondria-associated ER membranes (MAM). In these areas, the ER membrane and the mitochondrial outer membrane become as close as 10 to 30 nm, anchored by specific tether proteins (380). The interaction of the two organelles in the MAM areas is critical for many cellular activities that include mitochondrial biogenesis and transport as well as the initiation of apoptosis (381, 382). A key molecular component involved in the interaction is the voltage-dependent anionic channel (VDAC), a porin-like channel in the outer membrane of the mitochondria that forms a macromolecular complex with the IP3R channel in the ER membrane (383, 384). The activity of VDAC seems regulated by the release of calcium through the IP3R but the exact mechanisms governing gating and ion selectivity still remain controversial (385). The difficulty in resolving this issue may be due to the channel being regulated by the electrical activity of the ER membrane. Disruptions of the ER-mitochondrial interactions have been found in several neurodegenerative disorders that include Alzheimer’s disease, Parkinson and amyotrophic lateral sclerosis with associated frontotemporal dementia (ALS/FTD) (386, 387).

Lysosomes and endosomes also establish tight contacts with the ER membrane in specialized areas known as membrane contact sites (MCSs). These interactions have been found to play important roles in organelle biogenesis and in autophagy and cell survival (388, 389).

The interactions of the ER membrane with the nuclear matrix are reviewed in (390–393).

With the complex set of dynamic states and subcellular interactions, the ER is ideally suited to drive the coordinate orchestration of all the complex transformations that take place in cells during fertilization, embryonic development and in the different physiological activities of the adult life.

Transformations that involve the rearrangement of the cytoskeleton and chromatin structure, changes in the program of gene expression, the activation of secretion and other cellular processes such as mRNA translation, and even the initiation of programmed cell death.

A missing piece of information in this whole picture is the nature of the mechanism that mediates the complex signaling between the different subcellular compartments of the cell, between the plasma membrane and the nucleus, between different dendritic spines and between the spines and the nucleus in neurons, between components of the secretory pathway and mitochondria, to mention just a few examples. The excitability of the ER membrane provides a plausible mechanism to mediate this flow of information inside the cell as conjectured by Ruska in the 1950s (183).

In contrast with the impressive progress in characterizing the fine structure of the ER, the number of studies aimed at the evaluation and characterization of the electrical activity of the ER membrane is very limited. The activity, nevertheless, has been observed in the form of cytosolic calcium waves and oscillations. Calcium waves cross the oocyte during fecundation. There are calcium waves in the dendrites of excited neurons that communicate the stimulated synapses with the nucleus. Calcium oscillations are produced in the cytosol of *β*-cells when they are induced to secrete insulin. Underneath these calcium waves there are electrical oscillations of the ER membrane, as shown in this theoretical analysis. Many of the effects traditionally attributed to calcium may in fact be caused by these electrical signals. There are therefore good reasons to address the study of the electrical activity of the ER membrane, a study that is likely to produce important advances in our understanding of the role this versatile organelle plays in the physiology of cells both in health and disease.

## References

1. Saltiel, A. R., 2016. Insulin Signaling in the Control of Glucose and Lipid Homeostasis. Handbook of Experimental Pharmacology 233:51–71.

2. Rorsman, P., and M. Braun, 2013. Regulation of Insulin Secretion in Human Pancreatic Islets. Annual Review of Physiology 75:155–179.

3. Newsholme, P., V. Cruzat, F. Arfuso, and K. Keane, 2014. Nutrient Regulation of Insulin Secretion and Action. The Journal of Endocrinology 221:R105–120.

4. Nessa, A., S. A. Rahman, and K. Hussain, 2016. Hyperinsulinemic Hypoglycemia - The Molecular Mechanisms. Frontiers in Endocrinology 7:29.

5. Shah, P., S. A. Rahman, H. Demirbilek, M. Güemes, and K. Hussain, 2016. Hyperinsulinaemic Hypoglycaemia in Children and Adults. The Lancet. Diabetes & Endocrinology.

6. Stumvoll, M., B. J. Goldstein, and T. W. van Haeften, 2005 Apr 9-15. Type 2 Diabetes: Principles of Pathogenesis and Therapy. Lancet (London, England) 365:1333–1346.

7. Marchetti, P., R. Lupi, S. Del Guerra, M. Bugliani, L. Marselli, and U. Boggi, 2010. The Beta-Cell in Human Type 2 Diabetes. Advances in Experimental Medicine and Biology 654:501–514.

8. Rutter, G. A., T. J. Pullen, D. J. Hodson, and A. Martinez-Sanchez, 2015. Pancreatic *β*-Cell Identity, Glucose Sensing and the Control of Insulin Secretion. The Biochemical Journal 466:203–218.

9. Dean, P. M., and E. K. Matthews, 1968. Electrical Activity in Pancreatic Islet Cells. Nature 219:389–390.

10. Dean, P. M., and E. K. Matthews, 1970. Glucose-Induced Electrical Activity in Pancreatic Islet Cells. The Journal of Physiology 210:255–264.

11. Curry, D. L., L. L. Bennett, and G. M. Grodsky, 1968. Dynamics of Insulin Secretion by the Perfused Rat Pancreas. Endocrinology 83:572–584.

12. Fritsche, A., N. Stefan, E. Hardt, S. Schützenauer, H. Häring, and M. Stumvoll, 2000. A Novel Hyperglycaemic Clamp for Characterization of Islet Function in Humans: Assessment of Three Different Secretagogues, Maximal Insulin Response and Reproducibility. European Journal of Clinical Investigation 30:411–418.

13. Cheng, K., S. Andrikopoulos, and J. E. Gunton, 2013. First Phase Insulin Secretion and Type 2 Diabetes. Current Molecular Medicine 13:126–139.

14. Meissner, H. P., and I. J. Atwater, 1976. The Kinetics of Electrical Activity of Beta Cells in Response to a “Square Wave” Stimulation with Glucose or Glibenclamide. Hormone and Metabolic Research = Hormon- Und Stoffwechselforschung = Hormones Et Metabolisme 8:11–16.

15. Ashcroft, F. M., and P. Rorsman, 1989. Electrophysiology of the Pancreatic Beta-Cell. Progress in Biophysics and Molecular Biology 54:87–143.

16. Atwater, I., E. Rojas, and A. Scott, 1979. Simultaneous Measurements of Insulin Release and Electrical Activity from Single Microdissected Mouse Islets of Langerhans [Proceedings]. The Journal of Physiology 291:57P.

17. Rosario, L. M., I. Atwater, and A. M. Scott, 1986. Pulsatile Insulin Release and Electrical Activity from Single Ob/Ob Mouse Islets of Langerhans. Advances in Experimental Medicine and Biology 211:413–425.

18. Gilon, P., R. M. Shepherd, and J. C. Henquin, 1993. Oscillations of Secretion Driven by Oscillations of Cytoplasmic Ca2+ as Evidences in Single Pancreatic Islets. The Journal of Biological Chemistry 268:22265–22268.

19. Barbosa, R. M., A. M. Silva, A. R. Tomè, J. A. Stamford, R. M. Santos, and L. M. Rosàrio, 1998. Control of Pulsatile 5-HT/Insulin Secretion from Single Mouse Pancreatic Islets by Intracellular Calcium Dynamics. The Journal of Physiology 510 ( Pt 1):135–143.

20. Gilon, P., M. A. Ravier, J.-C. Jonas, and J.-C. Henquin, 2002. Control Mechanisms of the Oscillations of Insulin Secretion in Vitro and in Vivo. Diabetes 51 Suppl 1:S144–151.

21. Valdeolmillos, M., R. M. Santos, D. Contreras, B. Soria, and L. M. Rosario, 1989. Glucose-Induced Oscillations of Intracellular Ca2+ Concentration Resembling Bursting Electrical Activity in Single Mouse Islets of Langerhans. FEBS letters 259:19–23.

22. Santos, R. M., L. M. Rosario, A. Nadal, J. Garcia-Sancho, B. Soria, and M. Valdeolmillos, 1991. Widespread Synchronous [Ca2+]i Oscillations Due to Bursting Electrical Activity in Single Pancreatic Islets. Pflügers Archiv: European Journal of Physiology 418:417–422.

23. Roe, M. W., R. J. Mertz, M. E. Lancaster, J. F. Worley, and I. D. Dukes, 1994. Thapsigargin Inhibits the Glucose-Induced Decrease of Intracellular Ca2+ in Mouse Islets of Langerhans. The American Journal of Physiology 266:E852–862.

24. Komatsu, M., T. Schermerhorn, M. Noda, S. G. Straub, T. Aizawa, and G. W. Sharp, 1997. Augmentation of Insulin Release by Glucose in the Absence of Extracellular Ca2+: New Insights into Stimulus-Secretion Coupling. Diabetes 46:1928–1938.

25. Henquin, J. C., M. Nenquin, M. A. Ravier, and A. Szollosi, 2009. Shortcomings of Current Models of Glucose-Induced Insulin Secretion. Diabetes, Obesity & Metabolism 11 Suppl 4:168–179.

26. Prentki, M., F. M. Matschinsky, and S. R. M. Madiraju, 2013. Metabolic Signaling in Fuel-Induced Insulin Secretion. Cell Metabolism 18:162–185.

27. Ashcroft, F. M., and P. Rorsman, 2013. K(ATP) Channels and Islet Hormone Secretion: New Insights and Controversies. Nature Reviews. Endocrinology 9:660–669.

28. Ashcroft, F. M., D. E. Harrison, and S. J. Ashcroft, 1984 Nov 29-Dec 5. Glucose Induces Closure of Single Potassium Channels in Isolated Rat Pancreatic Beta-Cells. Nature 312:446–448.

29. Cook, D. L., and C. N. Hales, 1984 Sep 20-26. Intracellular ATP Directly Blocks K+ Channels in Pancreatic β-Cells. Nature 311:271–273.

30. Detimary, P., G. Van den Berghe, and J. C. Henquin, 1996. Concentration Dependence and Time Course of the Effects of Glucose on Adenine and Guanine Nucleotides in Mouse Pancreatic Islets. The Journal of Biological Chemistry 271:20559–20565.

31. Henquin, J. C., 2000. Triggering and Amplifying Pathways of Regulation of Insulin Secretion by Glucose. Diabetes 49:1751–1760.

32. Sato, Y., T. Aizawa, M. Komatsu, N. Okada, and T. Yamada, 1992. Dual Functional Role of Membrane Depolarization/Ca2+ Influx in Rat Pancreatic B-Cell. Diabetes 41:438–443.

33. Gembal, M., P. Gilon, and J. C. Henquin, 1992. Evidence That Glucose Can Control Insulin Release Independently from Its Action on ATP-Sensitive K+ Channels in Mouse B Cells. The Journal of Clinical Investigation 89:1288–1295.

34. Best, L., A. P. Yates, and S. Tomlinson, 1992. Stimulation of Insulin Secretion by Glucose in the Absence of Diminished Potassium (86Rb+) Permeability. Biochemical Pharmacology 43:2483–2485.

35. Komatsu, M., T. Schermerhorn, T. Aizawa, and G. W. Sharp, 1995. Glucose Stimulation of Insulin Release in the Absence of Extracellular Ca2+ and in the Absence of Any Increase in Intracellular Ca2+ in Rat Pancreatic Islets. Proceedings of the National Academy of Sciences of the United States of America 92:10728–10732.

36. Seghers, V., M. Nakazaki, F. DeMayo, L. Aguilar-Bryan, and J. Bryan, 2000. Sur1 Knockout Mice. A Model for K(ATP) Channel-Independent Regulation of Insulin Secretion. The Journal of Biological Chemistry 275:9270–9277.

37. Szollosi, A., M. Nenquin, L. Aguilar-Bryan, J. Bryan, and J.-C. Henquin, 2007. Glucose Stimulates Ca2+ Influx and Insulin Secretion in 2-Week-Old Beta-Cells Lacking ATP-Sensitive K+ Channels. The Journal of Biological Chemistry 282:1747–1756.

38. Ravier, M. A., M. Nenquin, T. Miki, S. Seino, and J.-C. Henquin, 2009. Glucose Controls Cy-tosolic Ca2+ and Insulin Secretion in Mouse Islets Lacking Adenosine Triphosphate-Sensitive K+ Channels Owing to a Knockout of the Pore-Forming Subunit Kir6.2. Endocrinology 150:33–45.

39. Miki, T., K. Nagashima, F. Tashiro, K. Kotake, H. Yoshitomi, A. Tamamoto, T. Gonoi, T. Iwanaga, J. Miyazaki, and S. Seino, 1998. Defective Insulin Secretion and Enhanced Insulin Action in KATP Channel-Deficient Mice. Proceedings of the National Academy of Sciences of the United States of America 95:10402–10406.

40. Shiota, C., O. Larsson, K. D. Shelton, M. Shiota, A. M. Efanov, M. Hoy, J. Lindner, S. Koop-tiwut, L. Juntti-Berggren, J. Gromada, P.-O. Berggren, and M. A. Magnuson, 2002. Sulfonylurea Receptor Type 1 Knock-out Mice Have Intact Feeding-Stimulated Insulin Secretion despite Marked Impairment in Their Response to Glucose. The Journal of Biological Chemistry 277:37176–37183.

41. Düfer, M., D. Haspel, P. Krippeit-Drews, L. Aguilar-Bryan, J. Bryan, and G. Drews, 2004. Oscillations of Membrane Potential and Cytosolic Ca(2+) Concentration in SUR1(−/−) Beta Cells. Diabetologia 47:488–498.

42. Gresset, A., J. Sondek, and T. K. Harden, 2012. The Phospholipase C Isozymes and Their Regulation. Sub-Cellular Biochemistry 58:61–94.

43. Berridge, M. J., 2007. Inositol Trisphosphate and Calcium Oscillations. Biochemical Society Symposium 1–7.

44. Mak, D.-O. D., and J. K. Foskett, 2014. Inositol 1,4,5-Trisphosphate Receptors in the Endoplasmic Reticulum: A Single-Channel Point of View. Cell Calcium.

45. Kaftan, E. J., B. E. Ehrlich, and J. Watras, 1997. Inositol 1,4,5-Trisphosphate (InsP3) and Calcium Interact to Increase the Dynamic Range of InsP3 Receptor-Dependent Calcium Signaling. The Journal of General Physiology 110:529–538.

46. Foskett, J. K., C. White, K.-H. Cheung, and D.-O. D. Mak, 2007. Inositol Trisphosphate Receptor Ca2+ Release Channels. Physiological Reviews 87:593–658.

47. Putney, J. W., 2011. The Physiological Function of Store-Operated Calcium Entry. Neurochemical Research 36:1157–1165.

48. Wollheim, C. B., and T. J. Biden, 1986. Second Messenger Function of Inositol 1,4,5-Trisphosphate. Early Changes in Inositol Phosphates, Cytosolic Ca2+, and Insulin Release in Carbamylcholine-Stimulated RINm5F Cells. The Journal of Biological Chemistry 261:8314–8319.

49. Santos, R. M., and E. Rojas, 1989. Muscarinic Receptor Modulation of Glucose-Induced Electrical Activity in Mouse Pancreatic B-Cells. FEBS letters 249:411–417.

50. Kaneko, Y. K., and T. Ishikawa, 2015. Diacylglycerol Signaling Pathway in Pancreatic β-Cells: An Essential Role of Diacylglycerol Kinase in the Regulation of Insulin Secretion. Biological & Pharmaceutical Bulletin 38:669–673.

51. Zawalich, W. S., and K. C. Zawalich, 2001. Effects of Protein Kinase C Inhibitors on Insulin Secretory Responses from Rodent Pancreatic Islets. Molecular and Cellular Endocrinology 177:95–105.

52. Carpenter, L., C. J. Mitchell, Z. Z. Xu, P. Poronnik, G. W. Both, and T. J. Biden, 2004. PKC Alpha Is Activated but Not Required during Glucose-Induced Insulin Secretion from Rat Pancreatic Islets. Diabetes 53:53–60.

53. Cantley, J., J. G. Burchfield, G. L. Pearson, C. Schmitz-Peiffer, M. Leitges, and T. J. Biden, 2009. Deletion of PKCepsilon Selectively Enhances the Amplifying Pathways of Glucose-Stimulated Insulin Secretion via Increased Lipolysis in Mouse Beta-Cells. Diabetes 58:1826–1834.

54. Gylfe, E., 1991. Carbachol Induces Sustained Glucose-Dependent Oscillations of Cytoplasmic Ca2+ in Hyperpolarized Pancreatic Beta Cells. Pflügers Archiv: European Journal of Physiology 419:639–643.

55. Gagerman, E., J. Sehlin, and I. B. Taljedal, 1980. Effects of Acetylcholine on Ion Fluxes and Chlorotetracycline Fluorescence in Pancreatic Islets. The Journal of Physiology 300:505–513.

56. Henquin, J. C., M. C. Garcia, M. Bozem, M. P. Hermans, and M. Nenquin, 1988. Muscarinic Control of Pancreatic B Cell Function Involves Sodium-Dependent Depolarization and Calcium Influx. Endocrinology 122:2134–2142.

57. Mears, D., and C. L. Zimliki, 2004. Muscarinic Agonists Activate Ca2+ Store-Operated and ‐Independent Ionic Currents in Insulin-Secreting HIT-T15 Cells and Mouse Pancreatic Beta-Cells. The Journal of Membrane Biology 197:59–70.

58. Swayne, L. A., A. Mezghrani, A. Varrault, J. Chemin, G. Bertrand, S. Dalle, E. Bourinet, P. Lory, R. J. Miller, J. Nargeot, and A. Monteil, 2009. The NALCN Ion Channel Is Activated by M3 Muscarinic Receptors in a Pancreatic Beta-Cell Line. EMBO reports 10:873–880.

59. Gilon, P., and J. C. Henquin, 2001. Mechanisms and Physiological Significance of the Cholinergic Control of Pancreatic Beta-Cell Function. Endocrine Reviews 22:565–604.

60. Nakajima, K., S. Jain, I. Ruiz de Azua, S. M. McMillin, M. Rossi, and J. Wess, 2013. Minireview: Novel Aspects of M3 Muscarinic Receptor Signaling in Pancreatic β-Cells. Molecular Endocrinology (Baltimore, Md.) 27:1208–1216.

61. Best, L., and W. J. Malaisse, 1983. Stimulation of Phosphoinositide Breakdown in Rat Pancreatic Islets by Glucose and Carbamylcholine. Biochemical and Biophysical Research Communications 116:9–16.

62. Montague, W., N. G. Morgan, G. M. Rumford, and C. A. Prince, 1985. Effect of Glucose on Polyphosphoinositide Metabolism in Isolated Rat Islets of Langerhans. The Biochemical Journal 227:483–489.

63. Kelley, G. G., K. C. Zawalich, and W. S. Zawalich, 1994. Calcium and a Mitochondrial Signal Interact to Stimulate Phosphoinositide Hydrolysis and Insulin Secretion in Rat Islets. Endocrinology 134:1648–1654.

64. Dunlop, M. E., and R. G. Larkins, 1984. The Role of Calcium in Phospholipid Turnover Following Glucose Stimulation in Neonatal Rat Cultured Islets. The Journal of Biological Chemistry 259:8407–8411.

65. Wollheim, C. B., and T. J. Biden, 1986. Signal Transduction in Insulin Secretion: Comparison between Fuel Stimuli and Receptor Agonists. Annals of the New York Academy of Sciences 488:317–333.

66. Zawalich, W. S., and K. C. Zawalich, 1996. Regulation of Insulin Secretion by Phospholipase C. American Journal of Physiology - Endocrinology and Metabolism 271:E409–E416.

67. Cerasi, E., 1975. Mechanisms of Glucose Stimulated Insulin Secretion in Health and in Diabetes: Some Re-Evaluations and Proposals. Diabetologia 11:1–13.

68. Rabinovitch, A., V. Grill, A. E. Renold, and E. Cerasi, 1976. Insulin Release and Cyclic AMP Accumulation in Response to Glucose in Pancreatic Islets of Fed and Starved Rats. The Journal of Clinical Investigation 58:1209–1216.

69. Dyachok, O., O. Idevall-Hagren, J. Sagetorp, G. Tian, A. Wuttke, C. Arrieumerlou, E. Gylfe Akusjarvi, and A. Tengholm, 2008. Glucose-Induced Cyclic AMP Oscillations Regulate Pulsatile Insulin Secretion. Cell Metabolism 8:26–37.

70. Tian, G., S. Sandler, E. Gylfe, and A. Tengholm, 2011. Glucose‐ and Hormone-Induced cAMP Oscillations in αߚ and β-Cells Within Intact Pancreatic Islets. Diabetes 60:1535–1543.

71. Tengholm, A., 2012. Cyclic AMP Dynamics in the Pancreatic β-Cell. Upsala Journal of Medical Sciences 117:355–369.

72. Ramos, L. S., J. H. Zippin, M. Kamenetsky, J. Buck, and L. R. Levin, 2008. Glucose and GLP-1 Stimulate cAMP Production via Distinct Adenylyl Cyclases in INS-1E Insulinoma Cells. The Journal of General Physiology 132:329–338

73. Holz, G. G., and G. Holz, 2004 Nov-Dec. New Insights Concerning the Glucose-Dependent Insulin Secretagogue Action of Glucagon-like Peptide-1 in Pancreatic Beta-Cells. Hormone and Metabolic Research = Hormon- Und Sto wechselforschung = Hormones Et Metabolisme 36:787–794

74. Drucker, D. J., 2006. The Biology of Incretin Hormones. Cell Metabolism 3:153–165.

75. Yabe, D., and Y. Seino, 2011. Two Incretin Hormones GLP-1 and GIP: Comparison of Their Actions in Insulin Secretion and Cell Preservation. Progress in Biophysics and Molecular Biology 107:248–256.

76. Doyle, M. E., and J. M. Egan, 2007. Mechanisms of Action of Glucagon-like Peptide 1 in the Pancreas. Pharmacology & Therapeutics 113:546–593

77. Ahren, B., 2009. Islet G Protein-Coupled Receptors as Potential Targets for Treatment of Type 2 Diabetes. Nature Reviews. Drug Discovery 8:369–385.

78. Ussher, J. R., and D. J. Drucker, 2014. Cardiovascular Actions of Incretin-Based Therapies. Circulation Research 114:1788–1803.

79. Nauck, M., 2016. Incretin Therapies: Highlighting Common Features and Differences in the Modes of Action of Glucagon-like Peptide-1 Receptor Agonists and Dipeptidyl Peptidase-4 Inhibitors. Diabetes, Obesity & Metabolism 18:203–216.

80. Shigeto, M., C. Y. Cha, P. Rorsman, and K. Kaku, 2017. A Role of PLC/PKC-Dependent Pathway in GLP-1-Stimulated Insulin Secretion. Journal of Molecular Medicine (Berlin, Germany).

81. Béguin, P., K. Nagashima, M. Nishimura, T. Gonoi, and S. Seino, 1999. PKA-Mediated Phosphorylation of the Human K(ATP) Channel: Separate Roles of Kir6.2 and SUR1 Subunit Phosphorylation. The EMBO journal 18:4722–4732.

82. Light, P. E., J. E. Manning Fox, M. J. Riedel, and M. B. Wheeler, 2002. Glucagon-like Peptide-1 Inhibits Pancreatic ATP-Sensitive Potassium Channels via a Protein Kinase A- and ADP-Dependent Mechanism. Molecular Endocrinology (Baltimore, Md.) 16:2135–2144.

83. Gloerich, M., and J. L. Bos, 2010. Epac: Defining a New Mechanism for cAMP Action. Annual Review of Pharmacology and Toxicology 50:355–375.

84. Schmidt, M., F. J. Dekker, and H. Maarsingh, 2013. Exchange Protein Directly Activated by cAMP (Epac): A Multidomain cAMP Mediator in the Regulation of Diverse Biological Functions. Pharmacological Reviews 65:670–709.

85. Kang, G., O. G. Chepurny, and G. G. Holz, 2001. cAMP-Regulated Guanine Nucleotide Exchange Factor II (Epac2) Mediates Ca2+-Induced Ca2+ Release in INS-1 Pancreatic Beta-Cells. The Journal of Physiology 536:375–385

86. Dzhura, I., O. G. Chepurny, G. G. Kelley, C. A. Leech, M. W. Roe, E. Dzhura, P. Afshari, S. Malik, M. J. Rindler, X. Xu, Y. Lu, A. V. Smrcka, and G. G. Holz, 2010. Epac2-Dependent Mobilization of Intracellular Ca2+ by Glucagon-like Peptide-1 Receptor Agonist Exendin-4 Is Disrupted in β-Cells of Phospholipase C-ε Knockout Mice. The Journal of Physiology 588:4871–4889

87. Dzhura, I., O. G. Chepurny, C. A. Leech, M. W. Roe, E. Dzhura, X. Xu, Y. Lu, F. Schwede, H.-G. Genieser, A. V. Smrcka, and G. G. Holz, 2011 May-Jun. Phospholipase C‐ Links Epac2 Activation to the Potentiation of Glucose-Stimulated Insulin Secretion from Mouse Islets of Langerhans. Islets 3:121–128.

88. Shigeto, M., R. Ramracheya, A. I. Tarasov, C. Y. Cha, M. V. Chibalina, B. Hastoy, K. Philippaert, T. Reinbothe, N. Rorsman, A. Salehi, W. R. Sones, E. Vergari, C. Weston, J. Gorelik, M. Katsura, V. O. Nikolaev, R. Vennekens, M. Zaccolo, A. Galione, P. R. V. Johnson, K. Kaku, G. Ladds, and P. Rorsman, 2015. GLP-1 Stimulates Insulin Secretion by PKC-Dependent TRPM4 and TRPM5 Activation. The Journal of Clinical Investigation 125:4714–4728

89. Calcraft, P. J., M. Ruas, Z. Pan, X. Cheng, A. Arredouani, X. Hao, J. Tang, K. Rietdorf, Teboul L., K.-T. Chuang, P. Lin, R. Xiao, C. Wang, Y. Zhu, Y. Lin, C. N. Wyatt, J. Parrington J., Ma, A. M. Evans, A. Galione, and M. X. Zhu, 2009. NAADP Mobilizes Calcium from Acidic Organelles through Two-Pore Channels. Nature 459:596–600.

90. Galione, A., 2015. A Primer of NAADP-Mediated Ca(2+) Signalling: From Sea Urchin Eggs to Mammalian Cells. Cell Calcium 58:27–47.

91. Masgrau, R., G. C. Churchill, A. J. Morgan, S. J. H. Ashcroft, and A. Galione, 2003. NAADP: A New Second Messenger for Glucose-Induced Ca2+ Responses in Clonal Pancreatic Beta Cells. Current biology: CB 13:247–251

92. Kim, B.-J., K.-H. Park, C.-Y. Yim, S. Takasawa, H. Okamoto, M.-J. Im, and U.-H. Kim, 2008. Generation of Nicotinic Acid Adenine Dinucleotide Phosphate and Cyclic ADP-Ribose by Glucagon-like Peptide-1 Evokes Ca2+ Signal That Is Essential for Insulin Secretion in Mouse Pancreatic Islets. Diabetes 57:868–878.

93. Naylor, E., A. Arredouani, S. R. Vasudevan, A. M. Lewis, R. Parkesh, A. Mizote, D. Rosen, J. M. Thomas, M. Izumi, A. Ganesan, A. Galione, and G. C. Churchill, 2009. Identification of a Chemical Probe for NAADP by Virtual Screening. Nature Chemical Biology 5:220–226.

94. Arredouani, A., M. Ruas, S. C. Collins, R. Parkesh, F. Clough, T. Pillinger, G. Coltart, K. Rietdorf, A. Royle, P. Johnson, M. Braun, Q. Zhang, W. Sones, K. Shimomura, A. J. Morgan, A. M. Lewis, K.-T. Chuang, R. Tunn, J. Gadea, L. Teboul, P. M. Heister, P. W. Tynan, E. A. Bellomo, G. A. Rutter, P. Rorsman, G. C. Churchill, J. Parrington, and A. Galione, 2015. Nicotinic Acid Adenine Dinucleotide Phosphate (NAADP) and Endolysosomal Two-Pore Channels Modulate Membrane Excitability and Stimulus-Secretion Coupling in Mouse Pancreatic Cells. The Journal of Biological Chemistry 290:21376–21392.

95. Cane, M. C., J. Parrington, P. Rorsman, A. Galione, and G. A. Rutter, 2016. The Two Pore Channel TPC2 Is Dispensable in Pancreatic β-Cells for Normal Ca(2+) Dynamics and Insulin Secretion. Cell Calcium 59:32–40.

96. Parekh, A. B., 2006. On the Activation Mechanism of Store-Operated Calcium Channels. Pflügers Archiv: European Journal of Physiology 453:303–311

97. Prakriya, M., and R. S. Lewis, 2015. Store-Operated Calcium Channels. Physiological Reviews 95:1383–1436.

98. Putney, J. W., 1986. A Model for Receptor-Regulated Calcium Entry. Cell Calcium 7:1–12.

99. Hoth, M., and R. Penner, 1992. Depletion of Intracellular Calcium Stores Activates a Calcium Current in Mast Cells. Nature 355:353–356.

100. Prakriya, M., S. Feske, Y. Gwack, S. Srikanth, A. Rao, and P. G. Hogan, 2006. Orai1 Is an Essential Pore Subunit of the CRAC Channel. Nature 443:230–233.

101. Roos, J., P. J. DiGregorio, A. V. Yeromin, K. Ohlsen, M. Lioudyno, S. Zhang, O. Safrina, J. A. Kozak, S. L. Wagner, M. D. Cahalan, G. Velicelebi, and K. A. Stauderman, 2005. STIM1, an Essential and Conserved Component of Store-Operated Ca2+ Channel Function. The Journal of Cell Biology 169:435–445

102. Zhang, S. L., Y. Yu, J. Roos, J. A. Kozak, T. J. Deerinck, M. H. Ellisman, K. A. Stauderman, and M. D. Cahalan, 2005. STIM1 Is a Ca2+ Sensor That Activates CRAC Channels and Migrates from the Ca2+ Store to the Plasma Membrane. Nature 437:902–905.

103. Choi, S., J. Maleth, A. Jha, K. P. Lee, M. S. Kim, I. So, M. Ahuja, and S. Muallem, 2014. The TRPCs-STIM1-Orai Interaction. Handbook of Experimental Pharmacology 223:1035–1054.

104. Flockerzi, V., 2007. An Introduction on TRP Channels. Handbook of Experimental Pharmacology 1–19.

105. Minke, B., and Z. Selinger, 1996. Role of Drosophila TRP in Inositide-Mediated Ca2+ Entry. Molecular Neurobiology 12:163–180.

106. Minke, B., and M. Parnas, 2006. Insights on TRP Channels from in Vivo Studies in Drosophila. Annual Review of Physiology 68:649–684.

107. Tamarina, N. A., A. Kuznetsov, and L. H. Philipson, 2008. Reversible Translocation of EYFP-Tagged STIM1 Is Coupled to Calcium Influx in Insulin Secreting Beta-Cells. Cell Calcium 44:533–544.

108. Jacobson, D. A., and L. H. Philipson, 2007. TRP Channels of the Pancreatic Beta Cell. Handbook of Experimental Pharmacology 409–424.

109. Sabourin, J., L. Le Gal, L. Saurwein, J.-A. Haefliger, E. Raddatz, and F. Allagnat, 2015. Store-Operated Ca2+ Entry Mediated by Orai1 and TRPC1 Participates to Insulin Secretion in Rat β-Cells. The Journal of Biological Chemistry 290:30530–30539.

110. Pørez, C. A., L. Huang, M. Rong, J. A. Kozak, A. K. Preuss, H. Zhang, M. Max, and R. F. Margolskee, 2002. A Transient Receptor Potential Channel Expressed in Taste Receptor Cells. Nature Neuroscience 5:1169–1176.

111. Zhang, Z., Z. Zhao, R. Margolskee, and E. Liman, 2007. The Transduction Channel TRPM5 Is Gated by Intracellular Calcium in Taste Cells. The Journal of Neuroscience: The Offcial Journal of the Society for Neuroscience 27:5777–5786

112. Liman, E. R., 2014. TRPM5. Handbook of Experimental Pharmacology 222:489–502.

113. Prawitt, D., M. K. Monteilh-Zoller, L. Brixel, C. Spangenberg, B. Zabel, A. Fleig, and R. Penner, 2003. TRPM5 Is a Transient Ca2+-Activated Cation Channel Responding to Rapid Changes in [Ca2+]I. Proceedings of the National Academy of Sciences of the United States of America 100:15166–15171.

114. Colsoul, B., A. Schraenen, K. Lemaire, R. Quintens, L. Van Lommel, A. Segal, G. Owsianik, K. Talavera, T. Voets, R. F. Margolskee, Z. Kokrashvili, P. Gilon, B. Nilius, F. C. Schuit, and R. Vennekens, 2010. Loss of High-Frequency Glucose-Induced Ca2+ Oscillations in Pancreatic Islets Correlates with Impaired Glucose Tolerance in Trpm5−/− Mice. Proceedings of the National Academy of Sciences of the United States of America 107:5208–5213.

115. Brixel, L. R., M. K. Monteilh-Zoller, C. S. Ingenbrandt, A. Fleig, R. Penner, T. Enklaar, B. U. Zabel, and D. Prawitt, 2010. TRPM5 Regulates Glucose-Stimulated Insulin Secretion. Pflügers Archiv: European Journal of Physiology 460:69–76

116. Uchida, K., K. Dezaki, B. Damdindorj, H. Inada, T. Shiuchi, Y. Mori, T. Yada, Y. Minokoshi, and M. Tominaga, 2011. Lack of TRPM2 Impaired Insulin Secretion and Glucose Metabolisms in Mice. Diabetes 60:119–126.

117. Krishnan, K., Z. Ma, A. Bjorklund, and M. S. Islam, 2014. Role of Transient Receptor Potential Melastatin-like Subtype 5 Channel in Insulin Secretion from Rat β-Cells. Pancreas 43:597–604.

118. Ketterer, C., K. Mussig, M. Heni, K. Dudziak, E. Randrianarisoa, R. Wagner, F. Machicao, N. Stefan, J. J. Holst, A. Fritsche, H.-U. Haring, and H. Staiger, 2011. Genetic Variation within the TRPM5 Locus Associates with Prediabetic Phenotypes in Subjects at Increased Risk for Type 2 Diabetes. Metabolism: Clinical and Experimental 60:1325–1333.

119. Drews, G., P. Krippeit-Drews, and M. Dufer, 2010. Electrophysiology of Islet Cells. Advances in Experimental Medicine and Biology 654:115–163.

120. Mart n, F., and B. Soria, 1996. Glucose-Induced [Ca2+]i Oscillations in Single Human Pancreatic Islets. Cell Calcium 20:409–414.

121. Sanchez-Andres, J. V., A. Gomis, and M. Valdeolmillos, 1995. The Electrical Activity of Mouse Pancreatic Beta-Cells Recorded in Vivo Shows Glucose-Dependent Oscillations. The Journal of Physiology 486 (Pt 1):223–228.

122. Fernandez, J., and M. Valdeolmillos, 2000. Synchronous Glucose-Dependent [Ca(2+)](i) Oscillations in Mouse Pancreatic Islets of Langerhans Recorded in Vivo. FEBS letters 477:33–36.

123. Hellman, B., 2009. Pulsatility of Insulin Release-a Clinically Important Phenomenon. Upsala Journal of Medical Sciences 114:193–205.

124. Satin, L. S., P. C. Butler, J. Ha, and A. S. Sherman, 2015. Pulsatile Insulin Secretion, Impaired Glucose Tolerance and Type 2 Diabetes. Molecular Aspects of Medicine 42:61–77.

125. O’Rahilly, S., R. C. Turner, and D. R. Matthews, 1988. Impaired Pulsatile Secretion of Insulin in Relatives of Patients with Non-Insulin-Dependent Diabetes. The New England Journal of Medicine 318:1225–1230.

126. Polonsky, K. S., 1999. Evolution of Beta-Cell Dysfunction in Impaired Glucose Tolerance and Diabetes. Experimental and Clinical Endocrinology & Diabetes: Official Journal, German Society of Endocrinology [and] German Diabetes Association 107 Suppl 4:S124–127.

127. Ashcroft, F. M., and P. Rorsman, 2012. Diabetes Mellitus and the Cell: The Last Ten Years. Cell 148:1160–1171.

128. Lechner, H. A., D. A. Baxter, J. W. Clark, and J. H. Byrne, 1996. Bistability and Its Regulation by Serotonin in the Endogenously Bursting Neuron R15 in Aplysia. Journal of Neurophysiology 75:957–962.

129. Tse, A., and B. Hille, 1992. GnRH-Induced Ca2+ Oscillations and Rhythmic Hyperpolarizations of Pituitary Gonadotropes. Science (New York, N.Y.) 255:462–464.

130. Liu, X., and A. E. Herbison, 2008. Small-Conductance Calcium-Activated Potassium Channels Control Excitability and Firing Dynamics in Gonadotropin-Releasing Hormone (GnRH) Neurons. Endocrinology 149:3598–3604.

131. Chagnac-Amitai, Y., and B. W. Connors, 1989. Synchronized Excitation and Inhibition Driven by Intrinsically Bursting Neurons in Neocortex. Journal of Neurophysiology 62:1149–1162.

132. Paladini, C. A., and J. Roeper, 2014. Generating Bursts (and Pauses) in the Dopamine Midbrain Neurons. Neuroscience 282:109–121.

133. Dragicevic, E., J. Schiemann, and B. Liss, 2015. Dopamine Midbrain Neurons in Health and Parkinson’s Disease: Emerging Roles of Voltage-Gated Calcium Channels and ATP-Sensitive Potassium Channels. Neuroscience 284:798–814.

134. Sherman, A., 1996. Contributions of Modeling to Understanding Stimulus-Secretion Coupling in Pancreatic Beta-Cells. The American Journal of Physiology 271:E362–372

135. Bertram, R., A. Sherman, and L. S. Satin, 2010. Electrical Bursting, Calcium Oscillations, and Synchronization of Pancreatic Islets. Advances in Experimental Medicine and Biology 654:261–279.

136. Fridlyand, L. E., N. Tamarina, and L. H. Philipson, 2010. Bursting and Calcium Oscillations in Pancreatic Beta-Cells: Specific Pacemakers for Specific Mechanisms. American Journal of Physiology. Endocrinology and Metabolism 299:E517–532.

137. Felix-Mart nez, G. J., and J. R. God nez-Fernandez, 2014. Mathematical Models of Electrical Activity of the Pancreatic β-Cell: A Physiological Review. Islets 6.

138. Berggren, P.-O., and C. J. Barker, 2008. A Key Role for Phosphorylated Inositol Compounds in Pancreatic Beta-Cell Stimulus-Secretion Coupling. Advances in Enzyme Regulation 48:276–294.

139. Atwater, I., C. M. Dawson, B. Ribalet, and E. Rojas, 1979. Potassium Permeability Activated by Intracellular Calcium Ion Concentration in the Pancreatic Beta-Cell. The Journal of Physiology 288:575–588

140. Atwater, I., C. M. Dawson, A. Scott, G. Eddlestone, and E. Rojas, 1980. The Nature of the Oscillatory Behaviour in Electrical Activity from Pancreatic Beta-Cell. Hormone and Metabolic Research. Supplement Series Suppl 10:100–107

141. Chay, T. R., and J. Keizer, 1983. Minimal Model for Membrane Oscillations in the Pancreatic Beta-Cell. Biophysical Journal 42:181–190.

142. Hodgkin, A. L., and A. F. Huxley, 1952. A Quantitative Description of Membrane Current and Its Application to Conduction and Excitation in Nerve. The Journal of Physiology 117:500–544

143. Chay, T. R., 1990. Electrical Bursting and Intracellular Ca2+ Oscillations in Excitable Cell Models. Biological Cybernetics 63:15–23.

144. Keizer, J., and P. Smolen, 1991. Bursting Electrical Activity in Pancreatic Beta Cells Caused by Ca(2+)- and Voltage-Inactivated Ca2+ Channels. Proceedings of the National Academy of Sciences of the United States of America 88:3897–3901.

145. Göpel, S. O., T. Kanno, S. Barg, L. Eliasson, J. Galvanovskis, E. Renström, and P. Rorsman, 1999. Activation of Ca(2+)-Dependent K(+) Channels Contributes to Rhythmic Firing of Action Potentials in Mouse Pancreatic Beta Cells. The Journal of General Physiology 114:759–770

146. Goforth, P. B., R. Bertram, F. A. Khan, M. Zhang, A. Sherman, and L. S. Satin, 2002. Calcium-Activated K+ Channels of Mouse Beta-Cells Are Controlled by Both Store and Cytoplasmic Ca2+: Experimental and Theoretical Studies. The Journal of General Physiology 120:307–322

147. Zhang, M., K. Houamed, S. Kupershmidt, D. Roden, and L. S. Satin, 2005. Pharmacological Properties and Functional Role of Kslow Current in Mouse Pancreatic Beta-Cells: SK Channels Contribute to Kslow Tail Current and Modulate Insulin Secretion. The Journal of General Physiology 126:353–363

148. Guéguinou, M., A. ChantÔme, G. Fromont, P. Bougnoux, C. Vandier, and M. Potier-Cartereau, 2014. KCa and Ca(2+) Channels: The Complex Thought. Biochimica Et Biophysica Acta 1843:2322–2333.

149. Kukuljan, M., A. A. Goncalves, and I. Atwater, 1991. Charybdotoxin-Sensitive K(Ca) Channel Is Not Involved in Glucose-Induced Electrical Activity in Pancreatic Beta-Cells. The Journal of Membrane Biology 119:187–195.

150. Houamed, K. M., I. R. Sweet, and L. S. Satin, 2010. BK Channels Mediate a Novel Ionic Mechanism That Regulates Glucose-Dependent Electrical Activity and Insulin Secretion in Mouse Pancreatic β-Cells. The Journal of Physiology 588:3511–3523.

151. Garcia, M. L., H. G. Knaus, P. Munujos, R. S. Slaughter, and G. J. Kaczorowski, 1995. Charybdotoxin and Its Effects on Potassium Channels. The American Journal of Physiology 269:C1–10

152. Lebrun, P., I. Atwater, M. Claret, W. J. Malaisse, and A. Herchuelz, 1983. Resistance to Apamin of the Ca2+-Activated K+ Permeability in Pancreatic B-Cells. FEBS letters 161:41–44

153. Tamarina, N. A., Y. Wang, L. Mariotto, A. Kuznetsov, C. Bond, J. Adelman, and L. H. Philipson, 2003. Small-Conductance Calcium-Activated K+ Channels Are Expressed in Pancreatic Islets and Regulate Glucose Responses. Diabetes 52:2000–2006.

154. Keizer, J., and G. Magnus, 1989. ATP-Sensitive Potassium Channel and Bursting in the Pancreatic Beta Cell. A Theoretical Study. Biophysical Journal 56:229–242.

155. Magnus, G., and J. Keizer, 1998. Model of Beta-Cell Mitochondrial Calcium Handling and Electrical Activity. I. Cytoplasmic Variables. The American Journal of Physiology 274:C1158–1173

156. Magnus, G., and J. Keizer, 1998. Model of Beta-Cell Mitochondrial Calcium Handling and Electrical Activity. II. Mitochondrial Variables. The American Journal of Physiology 274:C1174–1184

157. Smolen, P., and J. Keizer, 1992. Slow Voltage Inactivation of Ca2+ Currents and Bursting Mechanisms for the Mouse Pancreatic Beta-Cell. The Journal of Membrane Biology 127:9–19

158. Tornheim, K., 1997. Are Metabolic Oscillations Responsible for Normal Oscillatory Insulin Secretion? Diabetes 46:1375–1380.

159. Diederichs, F., 2006. Mathematical Simulation of Membrane Processes and Metabolic Fluxes of the Pancreatic Beta-Cell. Bulletin of Mathematical Biology 68:1779–1818.

160. Bertram, R., and A. Sherman, 2004. A Calcium-Based Phantom Bursting Model for Pancreatic Islets. Bulletin of Mathematical Biology 66:1313–1344.

161. Bertram, R., A. Sherman, and L. S. Satin, 2007. Metabolic and Electrical Oscillations: Partners in Controlling Pulsatile Insulin Secretion. American Journal of Physiology. Endocrinology and Metabolism 293:E890–900.

162. Larsson, O., H. Kindmark, R. Brandstrom, B. Fredholm, and P. O. Berggren, 1996. Oscillations in KATP Channel Activity Promote Oscillations in Cytoplasmic Free Ca2+ Concentration in the Pancreatic Beta Cell. Proceedings of the National Academy of Sciences of the United States of America 93:5161–5165.

163. Ainscow, E. K., and G. A. Rutter, 2002. Glucose-Stimulated Oscillations in Free Cytosolic ATP Concentration Imaged in Single Islet Beta-Cells: Evidence for a Ca2+-Dependent Mechanism. Diabetes 51 Suppl 1:S162–170.

164. Li, J., H. Y. Shuai, E. Gylfe, and A. Tengholm, 2013. Oscillations of Sub-Membrane ATP in Glucose-Stimulated Beta Cells Depend on Negative Feedback from Ca(2+). Diabetologia 56:1577–1586.

165. Yildirim, V., S. Vadrevu, B. Thompson, L. S. Satin, and R. Bertram, 2017. Upregulation of an Inward Rectifying K+ Channel Can Rescue Slow Ca2+ Oscillations in K(ATP) Channel Deficient Pancreatic Islets. PLoS computational biology 13:e1005686.

166. Worley, J. F., M. S. McIntyre, B. Spencer, and I. D. Dukes, 1994. Depletion of Intracellular Ca2+ Stores Activates a Maitotoxin-Sensitive Nonselective Cationic Current in Beta-Cells. The Journal of Biological Chemistry 269:32055–32058.

167. Worley, J. F., M. S. McIntyre, B. Spencer, R. J. Mertz, M. W. Roe, and I. D. Dukes, 1994. Endoplasmic Reticulum Calcium Store Regulates Membrane Potential in Mouse Islet Beta-Cells. The Journal of Biological Chemistry 269:14359–14362.

168. Chay, T. R., 1987. The Effect of Inactivation of Calcium Channels by Intracellular Ca2+ Ions in the Bursting Pancreatic Beta-Cells. Cell Biophysics 11:77–90.

169. Chay, T. R., 1996. Electrical Bursting and Luminal Calcium Oscillation in Excitable Cell Models. Biological Cybernetics 75:419–431.

170. Chay, T. R., 1997. Effects of Extracellular Calcium on Electrical Bursting and Intracellular and Luminal Calcium Oscillations in Insulin Secreting Pancreatic Beta-Cells. Biophysical Journal 73:1673–1688.

171. Tengholm, A., B. Hellman, and E. Gylfe, 1999. Glucose Regulation of Free Ca2+ in the Endoplasmic Reticulum of Mouse Pancreatic Beta Cells. Journal of Biological Chemistry 274:36883–36890.

172. Bertram, R., P. Smolen, A. Sherman, D. Mears, I. Atwater, F. Martin, and B. Soria, 1995. A Role for Calcium Release-Activated Current (CRAC) in Cholinergic Modulation of Electrical Activity in Pancreatic Beta-Cells. Biophysical Journal 68:2323–2332.

173. Roe, M. W., J. F. Worley, F. Qian, N. Tamarina, A. A. Mittal, F. Dralyuk, N. T. Blair, R. J. Mertz, L. H. Philipson, and I. D. Dukes, 1998. Characterization of a Ca2+ Release-Activated Nonselective Cation Current Regulating Membrane Potential and [Ca2+] i Oscillations in Transgenically Derived β-Cells. Journal of Biological Chemistry 273:10402–10410.

174. Fridlyand, L. E., N. Tamarina, and L. H. Philipson, 2003. Modeling of Ca2+ Flux in Pancreatic β-Cells: Role of the Plasma Membrane and Intracellular Stores. American Journal of Physiology - Endocrinology and Metabolism 285:E138–E154

175. Solovyova, N., and A. Verkhratsky, 2002. Monitoring of Free Calcium in the Neuronal Endo-plasmic Reticulum: An Overview of Modern Approaches. Journal of Neuroscience Methods 122:1–12.

176. Alvarez, J., and M. Montero, 2002 Nov-Dec. Measuring [Ca2+] in the Endoplasmic Reticulum with Aequorin. Cell Calcium 32:251–260.

177. Burdakov, D., O. H. Petersen, and A. Verkhratsky, 2005 Sep-Oct. Intraluminal Calcium as a Primary Regulator of Endoplasmic Reticulum Function. Cell Calcium 38:303–310.

178. Yamashita, M., M. Sugioka, and Y. Ogawa, 2006. Voltage‐ and Ca2+-Activated Potassium Channels in Ca2+ Store Control Ca2+ Release. The FEBS journal 273:3585–3597

179. Berridge, M. J., 2002 Nov-Dec. The Endoplasmic Reticulum: A Multifunctional Signaling Organelle. Cell Calcium 32:235–249.

180. Bootman, M. D., O. H. Petersen, and A. Verkhratsky, 2002 Nov-Dec. The Endoplasmic Reticulum Is a Focal Point for Co-Ordination of Cellular Activity. Cell Calcium 32:231–234.

181. Schuldiner, M., and B. Schwappach, 2013. From Rags to Riches - the History of the Endoplasmic Reticulum. Biochimica Et Biophysica Acta 1833:2389–2391.

182. Nixon-Abell, J., C. J. Obara, A. V. Weigel, D. Li, W. R. Legant, C. S. Xu, H. A. Pasolli, K. Harvey, H. F. Hess, E. Betzig, C. Blackstone, and J. Lippincott-Schwartz, 10 28, 2016. Increased Spatiotemporal Resolution Reveals Highly Dynamic Dense Tubular Matrices in the Peripheral ER. Science (New York, N.Y.) 354.

183. Ruska, H., G. A. Edwards, and R. Caesar, 1958. A Concept of Intracellular Transmission of Excitation by Means of the Endoplasmic Reticulum. Experientia 14:117–120.

184. Berridge, M. J., 2014. Calcium Regulation of Neural Rhythms, Memory and Alzheimer’s Disease. The Journal of Physiology 592:281–293

185. Berridge, M. J., 2016. The Inositol Trisphosphate/Calcium Signaling Pathway in Health and Disease. Physiological Reviews 96:1261–1296.

186. Iino, M., 1990. Biphasic Ca2+ Dependence of Inositol 1,4,5-Trisphosphate-Induced Ca Release in Smooth Muscle Cells of the Guinea Pig Taenia Caeci. The Journal of General Physiology 95:1103–1122

187. Bezprozvanny, I., J. Watras, and B. E. Ehrlich, 1991. Bell-Shaped Calcium-Response Curves of Ins(1,4,5)P3‐ and Calcium-Gated Channels from Endoplasmic Reticulum of Cerebellum. Nature 351:751–754.

188. Marshall, I. C., and C. W. Taylor, 1993. Biphasic Effects of Cytosolic Ca2+ on Ins(1,4,5)P3-Stimulated Ca2+ Mobilization in Hepatocytes. The Journal of Biological Chemistry 268:13214–13220.

189. Payne, R., B. Walz, S. Levy, and A. Fein, 1988. The Localization of Calcium Release by Inositol Trisphosphate in Limulus Photoreceptors and Its Control by Negative Feedback. Philosophical Transactions of the Royal Society of London. Series B, Biological Sciences 320:359–379.

190. Parker, I., and I. Ivorra, 1990. Inhibition by Ca2+ of Inositol Trisphosphate-Mediated Ca2+ Liberation: A Possible Mechanism for Oscillatory Release of Ca2+. Proceedings of the National Academy of Sciences of the United States of America 87:260–264.

191. Finch, E. A., T. J. Turner, and S. M. Goldin, 1991. Calcium as a Coagonist of Inositol 1,4,5-Trisphosphate-Induced Calcium Release. Science (New York, N.Y.) 252:443–446.

192. Ilyin, V., a I. Parker, 1994. Role of Cytosolic Ca2+ in Inhibition of InsP3-Evoked Ca2+ Release in Xenopus Oocytes. The Journal of Physiology 477 (Pt 3):503–509

193. Sneyd, J., and M. Falcke, 2005. Models of the Inositol Trisphosphate Receptor. Progress in Biophysics and Molecular Biology 89:207–245.

194. De Young, G. W., and J. Keizer, 1992. A Single-Pool Inositol 1,4,5-Trisphosphate-Receptor-Based Model for Agonist-Stimulated Oscillations in Ca2+ Concentration. Proceedings of the National Academy of Sciences of the United States of America 89:9895–9899.

195. Atri, A., J. Amundson, D. Clapham, and J. Sneyd, 1993. A Single-Pool Model for Intracellular Calcium Oscillations and Waves in the Xenopus Laevis Oocyte. Biophysical Journal 65:1727–1739.

196. Li, Y. X., and J. Rinzel, 1994. Equations for InsP3 Receptor-Mediated [Ca2+]i Oscillations Derived from a Detailed Kinetic Model: A Hodgkin-Huxley like Formalism. Journal of Theoretical Biology 166:461–473.

197. Tang, Y., J. L. Stephenson, and H. G. Othmer, 1996. Simplification and Analysis of Models of Calcium Dynamics Based on IP3-Sensitive Calcium Channel Kinetics. Biophysical Journal 70:246–263.

198. Li, Y. X., J. Keizer, S. S. Stojilkovic, and J. Rinzel, 1995. Ca2+ Excitability of the ER Membrane: An Explanation for IP3-Induced Ca2+ Oscillations. The American Journal of Physiology 269:C1079–1092

199. Keizer, J., Y. X. Li, S. Stojilkovic, and J. Rinzel, 1995. InsP3-Induced Ca2+ Excitability of the Endoplasmic Reticulum. Molecular Biology of the Cell 6:945–951.

200. al-Awqati, Q., 1995. Chloride Channels of Intracellular Organelles. Current Opinion in Cell Biology 7:504–508.

201. Takeshima, H., E. Venturi, and R. Sitsapesan, 2014. New and Notable Ion-Channels in the Sarcoplasmic/Endoplasmic Reticulum: Do They Support the Process of Intracellular Ca2+ Release? The Journal of Physiology

202. Kuum, M., V. Veksler, and A. Kaasik, 2015. Potassium Fluxes across the Endoplasmic Reticulum and Their Role in Endoplasmic Reticulum Calcium Homeostasis. Cell Calcium 58:79–85.

203. Shears, S. B., 1992. Metabolism of Inositol Phosphates. Advances in Second Messenger and Phosphoprotein Research 26:63–92.

204. Ermentrout, B., 2002. Simulating, Analyzing, and Animating Dynamical Systems: A Guide to XPPAUT for Researches and Students. SIAM, Philadelphia, first edition.

205. Del Prato, S., P. Marchetti, and R. C. Bonadonna, 2002. Phasic Insulin Release and Metabolic Regulation in Type 2 Diabetes. Diabetes 51 Suppl 1:S109–116.

206. Del Prato, S., 2003. Loss of Early Insulin Secretion Leads to Postprandial Hyperglycaemia. Diabetologia 46 Suppl 1:M2–8.

207. Cerasi, E., and R. Luft, 1967. The Plasma Insulin Response to Glucose Infusion in Healthy Subjects and in Diabetes Mellitus. Acta Endocrinologica 55:278–304.

208. Pratley, R. E., and C. Weyer, 2001. The Role of Impaired Early Insulin Secretion in the Pathogenesis of Type II Diabetes Mellitus. Diabetologia 44:929–945.

209. Del Prato, S., and A. Tiengo, 2001 May-Jun. The Importance of First-Phase Insulin Secretion: Implications for the Therapy of Type 2 Diabetes Mellitus. Diabetes/Metabolism Research and Reviews 17:164–174.

210. Grodsky, G. M., 1972. A Threshold Distribution Hypothesis for Packet Storage of Insulin and Its Mathematical Modeling. The Journal of Clinical Investigation 51:2047–2059

211. Henquin, J.-C., D. Dufrane, J. Kerr-Conte, and M. Nenquin, 2015. Dynamics of Glucose-Induced Insulin Secretion in Normal Human Islets. American Journal of Physiology. Endocrinology and Metabolism 309:E640–650

212. Zarain-Herzberg, A., G. García-Rivas, and R. Estrada-Avilés, 2014. Regulation of SERCA Pumps Expression in Diabetes. Cell Calcium 56:302–310.

213. Roe, M. W., L. H. Philipson, C. J. Frangakis, A. Kuznetsov, R. J. Mertz, M. E. Lancaster, B. Spencer, J. F. Worley, and I. D. Dukes, 1994. Defective Glucose-Dependent Endoplasmic Reticulum Ca2+ Sequestration in Diabetic Mouse Islets of Langerhans. The Journal of Biological Chemistry 269:18279–18282.

214. Váradi, A., E. Molnár, C. G. Ostenson, and S. J. Ashcroft, 1996. Isoforms of Endoplasmic Reticulum Ca(2+)-ATPase Are Differentially Expressed in Normal and Diabetic Islets of Langerhans. The Biochemical Journal 319 (Pt 2):521–527

215. Varadi, A., L. Lebel, Y. Hashim, Z. Mehta, S. J. Ashcroft, and R. Turner, 1999. Sequence Variants of the Sarco(Endo)Plasmic Reticulum Ca(2+)-Transport ATPase 3 Gene (SERCA3) in Caucasian Type II Diabetic Patients (UK Prospective Diabetes Study 48). Diabetologia 42:1240–1243.

216. Jackson, T. R., S. I. Patterson, O. Thastrup, and M. R. Hanley, 1988. A Novel Tumour Promoter, Thapsigargin, Transiently Increases Cytoplasmic Free Ca2+ without Generation of Inositol Phosphates in NG115-401L Neuronal Cells. The Biochemical Journal 253:81–86

217. Thastrup, O., P. J. Cullen, B. K. Drøbak, M. R. Hanley, and A. P. Dawson, 1990. Thapsigargin, a Tumor Promoter, Discharges Intracellular Ca2+ Stores by Specific Inhibition of the Endoplasmic Reticulum Ca2(+)-ATPase. Proceedings of the National Academy of Sciences of the United States of America 87:2466–2470.

218. Kirby, M. S., Y. Sagara, S. Gaa, G. Inesi, W. J. Lederer, and T. B. Rogers, 1992. Thapsigargin Inhibits Contraction and Ca2+ Transient in Cardiac Cells by Specific Inhibition of the Sarcoplasmic Reticulum Ca2+ Pump. The Journal of Biological Chemistry 267:12545–12551.

219. Gilon, P., A. Arredouani, P. Gailly, J. Gromada, and J.-C. Henquin, 1999. Uptake and Release of Ca2+ by the Endoplasmic Reticulum Contribute to the Oscillations of the Cytosolic Ca2+ Concentration Triggered by Ca2+ Influx in the Electrically Excitable Pancreatic B-Cell. Journal of Biological Chemistry 274:20197–20205.

220. Luciani, D. S., K. S. Gwiazda, T.-L. B. Yang, T. B. Kalynyak, Y. Bychkivska, M. H. Z. Frey, K. D. Jeffrey, A. V. Sampaio, T. M. Underhill, and J. D. Johnson, 2009. Roles of IP3R and RyR Ca2+ Channels in Endoplasmic Reticulum Stress and Beta-Cell Death. Diabetes 58:422–432.

221. Baggio, L. L., and D. J. Drucker, 2007. Biology of Incretins: GLP-1 and GIP. Gastroenterology 132:2131–2157.

222. Bloom, S. R., and A. V. Edwards, 1981. Pancreatic Endocrine Responses to Stimulation of the Peripheral Ends of the Vagus Nerves in Conscious Calves. The Journal of Physiology 315:31–41.

223. Garcia, M. C., M. P. Hermans, and J. C. Henquin, 1988. Glucose-, Calcium- and Concentration-Dependence of Acetylcholine Stimulation of Insulin Release and Ionic Fluxes in Mouse Islets. The Biochemical Journal 254:211–218

224. Persaud, S. J., P. M. Jones, D. Sugden, and S. L. Howell, 1989. The Role of Protein Kinase C in Cholinergic Stimulation of Insulin Secretion from Rat Islets of Langerhans. The Biochemical Journal 264:753–758

225. Todd, J. A., 2010. Etiology of Type 1 Diabetes. Immunity 32:457–467.

226. Wang, Z., Z. Xie, Q. Lu, C. Chang, and Z. Zhou, 2017. Beyond Genetics: What Causes Type 1 Diabetes. Clinical Reviews in Allergy & Immunology 52:273–286.

227. Prentki, M., and C. J. Nolan, 2006. Islet Beta Cell Failure in Type 2 Diabetes. The Journal of Clinical Investigation 116:1802–1812

228. Guillausseau, P.-J., T. Meas, M. Virally, M. Laloi-Michelin, V. Medeau, and J.-P. Kevorkian, 2008. Abnormalities in Insulin Secretion in Type 2 Diabetes Mellitus. Diabetes & Metabolism 34 Suppl 2:S43–48.

229. Cersosimo, E., C. Triplitt, L. J. Mandarino, and R. A. DeFronzo, 2000. Pathogenesis of Type 2 Diabetes Mellitus. In L. J. De Groot, G. Chrousos, K. Dungan, K. R. Feingold, A. Grossman, J. M. Hershman, C. Koch, M. Korbonits, R. McLachlan, M. New, J. Purnell, R. Rebar, F. Singer, and A. Vinik, editors, Endotext, MDText.com, Inc., South Dartmouth (MA).

230. Kahn, S. E., M. E. Cooper, and S. Del Prato, 2014. Pathophysiology and Treatment of Type 2 Diabetes: Perspectives on the Past, Present, and Future. Lancet (London, England) 383:1068–1083.

231. Lencioni, C., R. Lupi, and S. Del Prato, 2008. Beta-Cell Failure in Type 2 Diabetes Mellitus. Current Diabetes Reports 8:179–184.

232. Arner, P., T. Pollare, and H. Lithell, 1991. Different Aetiologies of Type 2 (Non-Insulin-Dependent) Diabetes Mellitus in Obese and Non-Obese Subjects. Diabetologia 34:483–487.

233. Sun, X., W. Yu, and C. Hu, 2014. Genetics of Type 2 Diabetes: Insights into the Pathogenesis and Its Clinical Application. BioMed Research International 2014:926713.

234. Wang, X., G. Strizich, Y. Hu, T. Wang, R. C. Kaplan, and Q. Qi, 2016. Genetic Markers of Type 2 Diabetes: Progress in Genome-Wide Association Studies and Clinical Application for Risk Prediction. Journal of Diabetes 8:24–35.

235. Alejandro, E. U., B. Gregg, M. Blandino-Rosano, C. Cras-Méneur, and E. Bernal-Mizrachi, 2015. Natural History of β-Cell Adaptation and Failure in Type 2 Diabetes. Molecular Aspects of Medicine 42:19–41.

236. Yki-Järvinen, H., 1992. Glucose Toxicity. Endocrine Reviews 13:415–431.

237. Lee, Y., H. Hirose, M. Ohneda, J. H. Johnson, J. D. McGarry, and R. H. Unger, 1994. Beta-Cell Lipotoxicity in the Pathogenesis of Non-Insulin-Dependent Diabetes Mellitus of Obese Rats: Impairment in Adipocyte-Beta-Cell Relationships. Proceedings of the National Academy of Sciences of the United States of America 91:10878–10882.

238. Supale, S., N. Li, T. Brun, and P. Maechler, 2012. Mitochondrial Dysfunction in Pancreatic *β* Cells. Trends in endocrinology and metabolism: TEM 23:477–487

239. Robertson, R. P., 2010. Antioxidant Drugs for Treating Beta-Cell Oxidative Stress in Type 2 Diabetes: Glucose-Centric versus Insulin-Centric Therapy. Discovery Medicine 9:132–137.

240. Haataja, L., T. Gurlo, C. J. Huang, and P. C. Butler, 2008. Islet Amyloid in Type 2 Diabetes, and the Toxic Oligomer Hypothesis. Endocrine Reviews 29:303–316.

241. Jurgens, C. A., M. N. Toukatly, C. L. Fligner, J. Udayasankar, S. L. Subramanian, S. Zraika, K. Aston-Mourney, D. B. Carr, P. Westermark, G. T. Westermark, S. E. Kahn, and R. L. Hull, 2011. β-Cell Loss and β-Cell Apoptosis in Human Type 2 Diabetes Are Related to Islet Amyloid Deposition. The American Journal of Pathology 178:2632–2640.

242. Fernández, M. S., 2014. Human IAPP Amyloidogenic Properties and Pancreatic β-Cell Death. Cell Calcium 56:416–427.

243. Back, S. H., and R. J. Kaufman, 2012. Endoplasmic Reticulum Stress and Type 2 Diabetes. Annual Review of Biochemistry 81:767–793.

244. Rabhi, N., E. Salas, P. Froguel, and J.-S. Annicotte, 2014. Role of the Unfolded Protein Response in *β* Cell Compensation and Failure during Diabetes. Journal of Diabetes Research 2014:795171.

245. Guardado-Mendoza, R., A. M. Davalli, A. O. Chavez, G. B. Hubbard, E. J. Dick, A. Majluf-Cruz, C. E. Tene-Perez, L. Goldschmidt, J. Hart, C. Perego, A. G. Comuzzie, M. E. Tejero, G. Finzi, C. Placidi, S. La Rosa, C. Capella, G. Halff, A. Gastaldelli, R. A. DeFronzo, and F. Folli, 2009. Pancreatic Islet Amyloidosis, Beta-Cell Apoptosis, and Alpha-Cell Proliferation Are Determinants of Islet Remodeling in Type-2 Diabetic Baboons. Proceedings of the National Academy of Sciences of the United States of America 106:13992–13997.

246. Kahn, S. E., S. Zraika, K. M. Utzschneider, and R. L. Hull, 2009. The Beta Cell Lesion in Type 2 Diabetes: There Has to Be a Primary Functional Abnormality. Diabetologia 52:1003–1012.

247. Thulé, P. M., and G. Umpierrez, 2014. Sulfonylureas: A New Look at Old Therapy. Current Diabetes Reports 14:473.

248. Klein-Schwartz, W., G. L. Stassinos, and G. K. Isbister, 2016. Treatment of Sulfonylurea and Insulin Overdose. British Journal of Clinical Pharmacology 81:496–504.

249. Knop, F. K., J. J. Holst, and T. Vilsbøll, 2008. Replacing SUs with Incretin-Based Therapies for Type 2 Diabetes Mellitus: Challenges and Feasibility. IDrugs: the investigational drugs journal 11:497–501

250. Hartman, I., E. Rojas, and D. Rodríguez-Molina, 2013 Jul-Aug. Incretin-Based Therapy for Type 2 Diabetes Mellitus: Pancreatic and Extrapancreatic E ects. American Journal of Therapeutics 20:384–393.

251. Pratley, R. E., and A. Salsali, 2007. Inhibition of DPP-4: A New Therapeutic Approach for the Treatment of Type 2 Diabetes. Current Medical Research and Opinion 23:919–931.

252. Butler, A. E., J. Janson, S. Bonner-Weir, R. Ritzel, R. A. Rizza, and P. C. Butler, 2003. Beta-Cell Deficit and Increased Beta-Cell Apoptosis in Humans with Type 2 Diabetes. Diabetes 52:102–110.

253. Baggio, L. L., and D. J. Drucker, 2006. Therapeutic Approaches to Preserve Islet Mass in Type 2 Diabetes. Annual Review of Medicine 57:265–281.

254. Gigoux, V., and D. Fourmy, 2013. Acting on Hormone Receptors with Minimal Side Effect on Cell Proliferation: A Timely Challenge Illustrated with GLP-1R and GPER. Frontiers in Endocrinology 4:50.

255. Sneyd, J., J. Keizer, and M. J. Sanderson, 1995. Mechanisms of Calcium Oscillations and Waves: A Quantitative Analysis. FASEB journal: official publication of the Federation of American Societies for Experimental Biology 9:1463–1472

256. Nayfeh, A. H., and D. T. Mook, 1995. Nonlinear Oscillations. John Wiley & Sons, Inc, New York.

257. Liang, K., W. Du, W. Zhu, S. Liu, Y. Cui, H. Sun, B. Luo, Y. Xue, L. Yang, L. Chen, and F. Li, 2011. Contribution of Different Mechanisms to Pancreatic Beta-Cell Hyper-Secretion in Non-Obese Diabetic (NOD) Mice during Pre-Diabetes. The Journal of Biological Chemistry 286:39537–39545.

258. Chami, M., D. Gozuacik, D. Lagorce, M. Brini, P. Falson, G. Peaucellier, P. Pinton, H. Lecoeur, M.-L. Gougeon, M. le Maire, R. Rizzuto, C. Bréchot, and P. Paterlini-Bréchot, 2001. Serca1 Truncated Proteins Unable to Pump Calcium Reduce the Endoplasmic Reticulum Calcium Concentration and Induce Apoptosis. The Journal of Cell Biology 153:1301–1314

259. Srinivasan, K., and P. Ramarao, 2007. Animal Models in Type 2 Diabetes Research: An Overview. The Indian Journal of Medical Research 125:451–472

260. King, A. J. F., 2012. The Use of Animal Models in Diabetes Research. British Journal of Pharmacology 166:877–894.

261. Liang, K., W. Du, J. Lu, F. Li, L. Yang, Y. Xue, B. Hille, and L. Chen, 2014. Alterations of the Ca2 Signaling Pathway in Pancreatic Beta-Cells Isolated from Db/Db Mice. Protein & Cell 5:783–794.

262. Al-Awar, A., K. Kupai, M. Veszelka, G. Szűcs, Z. Attieh, Z. Murlasits, S. Török, A. Pósa, and C. Varga, 2016. Experimental Diabetes Mellitus in Different Animal Models. Journal of Diabetes Research 2016:9051426.

263. Meigs, J. B., L. A. Cupples, and P. W. Wilson, 2000. Parental Transmission of Type 2 Diabetes: The Framingham Offspring Study. Diabetes 49:2201–2207.

264. Sladek, R., G. Rocheleau, J. Rung, C. Dina, L. Shen, D. Serre, P. Boutin, D. Vincent, A. Belisle, S. Hadjadj, B. Balkau, B. Heude, G. Charpentier, T. J. Hudson, A. Montpetit, A. V. Pshezhetsky, M. Prentki, B. I. Posner, D. J. Balding, D. Meyre, C. Polychronakos, and P. Froguel, 2007. A Genome-Wide Association Study Identifies Novel Risk Loci for Type 2 Diabetes. Nature 445:881–885.

265. Hemminki, K., X. Li, K. Sundquist, and J. Sundquist, 2010. Familial Risks for Type 2 Diabetes in Sweden. Diabetes Care 33:293–297.

266. Zeggini, E., M. N. Weedon, C. M. Lindgren, T. M. Frayling, K. S. Elliott, H. Lango, N. J. Timpson, J. R. B. Perry, N. W. Rayner, R. M. Freathy, J. C. Barrett, B. Shields, A. P. Morris, S. Ellard, C. J. Groves, L. W. Harries, J. L. Marchini, K. R. Owen, B. Knight, L. R. Cardon, M. Walker, G. A. Hitman, A. D. Morris, A. S. F. Doney, Wellcome Trust Case Control Consortium (WTCCC), M. I. McCarthy, and A. T. Hattersley, 2007. Replication of Genome-Wide Association Signals in UK Samples Reveals Risk Loci for Type 2 Diabetes. Science (New York, N.Y.) 316:1336–1341.

267. Kwak, S. H., and K. S. Park, 2013. Genetics of Type 2 Diabetes and Potential Clinical Implications. Archives of Pharmacal Research 36:167–177.

268. Kwak, S. H., and K. S. Park, 2016. Recent Progress in Genetic and Epigenetic Research on Type 2 Diabetes. Experimental & Molecular Medicine 48:e220.

269. McCarthy, M. I., 2010. Genomics, Type 2 Diabetes, and Obesity. The New England Journal of Medicine 363:2339–2350.

270. Bonnefond, A., N. Clement, K. Fawcett, L. Yengo, E. Vaillant, J.-L. Guillaume, A. Dechaume, F. Payne, R. Roussel, S. Czernichow, S. Hercberg, S. Hadjadj, B. Balkau, M. Marre, O. Lantieri, C. Langenberg, N. Bouatia-Naji, Meta-Analysis of Glucose and Insulin-Related Traits Consortium (MAGIC), G. Charpentier, M. Vaxillaire, G. Rocheleau, N. J. Wareham, R. Sladek, M. I. McCarthy, C. Dina, I. Barroso, R. Jockers, and P. Froguel, 2012. Rare MTNR1B Variants Impairing Melatonin Receptor 1B Function Contribute to Type 2 Diabetes. Nature Genetics 44:297–301.

271. Majithia, A. R., J. Flannick, P. Shahinian, M. Guo, M.-A. Bray, P. Fontanillas, S. B. Gabriel, GoT2D Consortium, NHGRI JHS/FHS Allelic Spectrum Project, SIGMA T2D Consortium, T2D-GENES Consortium, E. D. Rosen, and D. Altshuler, 2014. Rare Variants in PPARG with Decreased Activity in Adipocyte Differentiation Are Associated with Increased Risk of Type 2 Diabetes. Proceedings of the National Academy of Sciences of the United States of America 111:13127–13132.

272. Steinthorsdottir, V., G. Thorleifsson, P. Sulem, H. Helgason, N. Grarup, A. Sigurdsson, H. T. Helgadottir, H. Johannsdottir, O. T. Magnusson, S. A. Gudjonsson, J. M. Justesen, M. N. Harder, M. E. Jørgensen, C. Christensen, I. Brandslund, A. Sandbxk, T. Lauritzen, H. Vestergaard, A. Linneberg, T. J rgensen, T. Hansen, M. S. Daneshpour, M.-S. Fallah, A. B. Hreidarsson, G. Sigurdsson, F. Azizi, R. Benediktsson, G. Masson, A. Helgason, A. Kong, D. F. Gudbjartsson, O. Pedersen, U. Thorsteinsdottir, and K. Stefansson, 2014. Identification of Low-Frequency and Rare Sequence Variants Associated with Elevated or Reduced Risk of Type 2 Diabetes. Nature Genetics 46:294–298.

273. Zhang, Y., M. A. Hoon, J. Chandrashekar, K. L. Mueller, B. Cook, D. Wu, C. S. Zuker, and N. J. P. Ryba, 2003. Coding of Sweet, Bitter, and Umami Tastes: Different Receptor Cells Sharing Similar Signaling Pathways. Cell 112:293–301.

274. Hofmann, T., V. Chubanov, T. Gudermann, and C. Montell, 2003. TRPM5 Is a Voltage-Modulated and Ca(2+)-Activated Monovalent Selective Cation Channel. Current biology: CB 13:1153–1158

275. Liu, D., and E. R. Liman, 2003. Intracellular Ca2+ and the Phospholipid PIP2 Regulate the Taste Transduction Ion Channel TRPM5. Proceedings of the National Academy of Sciences of the United States of America 100:15160–15165.

276. Ullrich, N. D., T. Voets, J. Prenen, R. Vennekens, K. Talavera, G. Droogmans, and B. Nilius, 2005. Comparison of Functional Properties of the Ca2+-Activated Cation Channels TRPM4 and TRPM5 from Mice. Cell Calcium 37:267–278.

277. Berridge, M. J., 2009. Inositol Trisphosphate and Calcium Signalling Mechanisms. Biochimica Et Biophysica Acta 1793:933–940.

278. Miyazaki, S., 2006. Thirty Years of Calcium Signals at Fertilization. Seminars in Cell & Developmental Biology 17:233–243.

279. Swann, K., and Y. Yu, 2008. The Dynamics of Calcium Oscillations That Activate Mammalian Eggs. The International Journal of Developmental Biology 52:585–594

280. Webb, S. E., and A. L. Miller, 2013. Ca(2+) Signaling during Activation and Fertilization in the Eggs of Teleost Fish. Cell Calcium 53:24–31.

281. Sartain, C. V., and M. F. Wolfner, 2013. Calcium and Egg Activation in Drosophila. Cell Calcium 53:10–15.

282. Kline, D., and J. T. Kline, 1992. Repetitive Calcium Transients and the Role of Calcium in Exocytosis and Cell Cycle Activation in the Mouse Egg. Developmental Biology 149:80–89.

283. Miyazaki, S., M. Yuzaki, K. Nakada, H. Shirakawa, S. Nakanishi, S. Nakade, and K. Mikoshiba, 1992. Block of Ca2+ Wave and Ca2+ Oscillation by Antibody to the Inositol 1,4,5-Trisphosphate Receptor in Fertilized Hamster Eggs. Science (New York, N.Y.) 257:251–255.

284. Saunders, C. M., M. G. Larman, J. Parrington, L. J. Cox, J. Royse, L. M. Blayney, K. Swann, and F. A. Lai, 2002. PLC Zeta: A Sperm-Specific Trigger of Ca(2+) Oscillations in Eggs and Embryo Development. Development (Cambridge, England) 129:3533–3544.

285. Kashir, J., M. Nomikos, F. A. Lai, and K. Swann, 2014. Sperm-Induced Ca2+ Release during Egg Activation in Mammals. Biochemical and Biophysical Research Communications 450:1204–1211.

286. Cox, L. J., M. G. Larman, C. M. Saunders, K. Hashimoto, K. Swann, and F. A. Lai, 2002. Sperm Phospholipase Czeta from Humans and Cynomolgus Monkeys Triggers Ca2+ Oscillations, Activation and Development of Mouse Oocytes. Reproduction (Cambridge, England) 124:611–623.

287. Kouchi, Z., K. Fukami, T. Shikano, S. Oda, Y. Nakamura, T. Takenawa, and S. Miyazaki, 2004. Recombinant Phospholipase Czeta Has High Ca2+ Sensitivity and Induces Ca2+ Oscillations in Mouse Eggs. The Journal of Biological Chemistry 279:10408–10412.

288. Marangos, P., G. FitzHarris, and J. Carroll, 2003. Ca2+ Oscillations at Fertilization in Mammals Are Regulated by the Formation of Pronuclei. Development (Cambridge, England) 130:1461–1472.

289. Busa, W. B., and R. Nuccitelli, 1985. An Elevated Free Cytosolic Ca2+ Wave Follows Fertilization in Eggs of the Frog, Xenopus Laevis. The Journal of Cell Biology 100:1325–1329

290. Stricker, S. A., 1999. Comparative Biology of Calcium Signaling during Fertilization and Egg Activation in Animals. Developmental Biology 211:157–176.

291. York-Andersen, A. H., R. M. Parton, C. J. Bi, C. L. Bromley, I. Davis, and T. T. Weil, 2015. A Single and Rapid Calcium Wave at Egg Activation in Drosophila. Biology Open BIO201411296

292. Horner, V. L., and M. F. Wolfner, 2008. Transitioning from Egg to Embryo: Triggers and Mechanisms of Egg Activation. Developmental Dynamics: An Official Publication of the American Association of Anatomists 237:527–544

293. Krauchunas, A. R., and M. F. Wolfner, 2013. Molecular Changes during Egg Activation. Current Topics in Developmental Biology 102:267–292.

294. Whitaker, M., 2008. Calcium Signalling in Early Embryos. Philosophical Transactions of the Royal Society of London. Series B, Biological Sciences 363:1401–1418.

295. Miao, Y.-L., and C. J. Williams, 2012. Calcium Signaling in Mammalian Egg Activation and Embryo Development: The Influence of Subcellular Localization. Molecular Reproduction and Development 79:742–756.

296. Mikoshiba, K., 2011. Role of IP Receptor in Development. Cell Calcium 49:331–340.

297. Webb, S. E., R. A. Fluck, and A. L. Miller, 2011. Calcium Signaling during the Early Development of Medaka and Zebrafish. Biochimie 93:2112–2125.

298. Leclerc, C., I. Néant, and M. Moreau, 2011. Early Neural Development in Vertebrates Is Also a Matter of Calcium. Biochimie 93:2102–2111.

299. Pollizzi, K. N., and J. D. Powell, 2014. Integrating Canonical and Metabolic Signalling Programmes in the Regulation of T Cell Responses. Nature Reviews. Immunology 14:435–446.

300. Lewis, R. S., 2001. Calcium Signaling Mechanisms in T Lymphocytes. Annual Review of Immunology 19:497–521.

301. Gwack, Y., S. Feske, S. Srikanth, P. G. Hogan, and A. Rao, 2007. Signalling to Transcription: Store-Operated Ca2+ Entry and NFAT Activation in Lymphocytes. Cell Calcium 42:145–156.

302. Srikanth, S., and Y. Gwack, 2013. Orai1-NFAT Signalling Pathway Triggered by T Cell Receptor Stimulation. Molecules and Cells 35:182–194.

303. Baba, Y., and T. Kurosaki, 2016. Role of Calcium Signaling in B Cell Activation and Biology. Current Topics in Microbiology and Immunology 393:143–174.

304. Dolmetsch, R. E., R. S. Lewis, C. C. Goodnow, and J. I. Healy, 1997. Differential Activation of Transcription Factors Induced by Ca2+ Response Amplitude and Duration. Nature 386:855–858.

305. Lewis, R. S., 2003. Calcium Oscillations in T-Cells: Mechanisms and Consequences for Gene Expression. Biochemical Society Transactions 31:925–929.

306. Smedler, E., and P. Uhlen, 2014. Frequency Decoding of Calcium Oscillations. Biochimica Et Biophysica Acta 1840:964–969.

307. Feske, S., 2007. Calcium Signalling in Lymphocyte Activation and Disease. Nature Reviews. Immunology 7:690–702.

308. Kawamoto, E. M., C. Vivar, and S. Camandola, 2012. Physiology and Pathology of Calcium Signaling in the Brain. Frontiers in Pharmacology 3:61.

309. Baker, K. D., T. M. Edwards, and N. S. Rickard, 2013. The Role of Intracellular Calcium Stores in Synaptic Plasticity and Memory Consolidation. Neuroscience and Biobehavioral Reviews 37:1211–1239.

310. Mayford, M., S. A. Siegelbaum, and E. R. Kandel, 2012. Synapses and Memory Storage. Cold Spring Harbor Perspectives in Biology 4.

311. Byers, D., R. L. Davis, and J. A. Kiger, 1981. Defect in Cyclic AMP Phosphodiesterase Due to the Dunce Mutation of Learning in Drosophila Melanogaster. Nature 289:79–81.

312. Kahsai, L., and T. Zars, 2011. Learning and Memory in Drosophila: Behavior, Genetics, and Neural Systems. International Review of Neurobiology 99:139–167.

313. Bailey, C. H., E. R. Kandel, and K. M. Harris, 2015. Structural Components of Synaptic Plasticity and Memory Consolidation. Cold Spring Harbor Perspectives in Biology 7:a021758.

314. Cajal, S. R., 1894. La Fine Structure Des Centres Nerveux. Proceedings of the Royal Society of London 55:444–468

315. Tønnesen, J., G. Katona, B. Rozsa, and U. V. Nagerl, 2014. Spine Neck Plasticity Regulates Compartmentalization of Synapses. Nature Neuroscience 17:678–685.

316. Spacek, J., and K. M. Harris, 1997. Three-Dimensional Organization of Smooth Endoplasmic Reticulum in Hippocampal CA1 Dendrites and Dendritic Spines of the Immature and Mature Rat. The Journal of Neuroscience: The Official Journal of the Society for Neuroscience 17:190–203

317. Franks, K. M., and T. J. Sejnowski, 2002. Complexity of Calcium Signaling in Synaptic Spines. BioEssays: News and Reviews in Molecular, Cellular and Developmental Biology 24:1130–1144

318. Bardo, S., M. G. Cavazzini, and N. Emptage, 2006. The Role of the Endoplasmic Reticulum Ca2+ Store in the Plasticity of Central Neurons. Trends in Pharmacological Sciences 27:78–84.

319. Harris, K. M., and R. J. Weinberg, 2012. Ultrastructure of Synapses in the Mammalian Brain. Cold Spring Harbor Perspectives in Biology 4.

320. Whitlock, J. R., A. J. Heynen, M. G. Shuler, and M. F. Bear, 2006. Learning Induces Long-Term Potentiation in the Hippocampus. Science (New York, N.Y.) 313:1093–1097.

321. Lee, K. F. H., C. Soares, and J.-C. Beéïque, 2012. Examining Form and Function of Dendritic Spines. Neural Plasticity 2012:704103.

322. Segal, M., 2017. Dendritic Spines: Morphological Building Blocks of Memory. Neurobiology of Learning and Memory 138:3–9.

323. Lüscher, C., and R. C. Malenka, 2012. NMDA Receptor-Dependent Long-Term Potentiation and Long-Term Depression (LTP/LTD). Cold Spring Harbor Perspectives in Biology 4.

324. Bliss, T. V., and G. L. Collingridge, 1993. A Synaptic Model of Memory: Long-Term Potentiation in the Hippocampus. Nature 361:31–39.

325. Sweatt, J. D., 2004. Mitogen-Activated Protein Kinases in Synaptic Plasticity and Memory. Current Opinion in Neurobiology 14:311–317.

326. Barbara, J.-G., 2002. IP3-Dependent Calcium-Induced Calcium Release Mediates Bidirectional Calcium Waves in Neurones: Functional Implications for Synaptic Plasticity. Biochimica Et Biophysica Acta 1600:12–18.

327. Larkum, M. E., S. Watanabe, T. Nakamura, N. Lasser-Ross, and W. N. Ross, 2003. Synaptically Activated Ca2+ Waves in Layer 2/3 and Layer 5 Rat Neocortical Pyramidal Neurons. The Journal of Physiology 549:471–488

328. Castillo, P. E., C. Q. Chiu, and R. C. Carroll, 2011. Long-Term Plasticity at Inhibitory Synapses. Current Opinion in Neurobiology 21:328–338.

329. Rozov, A. V., F. F. Valiullina, and A. P. Bolshakov, 2017. Mechanisms of Long-Term Plasticity of Hippocampal GABAergic Synapses. Biochemistry. Biokhimiia 82:257–263.

330. Heifets, B. D., and P. E. Castillo, 2009. Endocannabinoid Signaling and Long-Term Synaptic Plasticity. Annual Review of Physiology 71:283–306.

331. Yoshii, A., and M. Constantine-Paton, 2010. Postsynaptic BDNF-TrkB Signaling in Synapse Maturation, Plasticity, and Disease. Developmental Neurobiology 70:304–322.

332. Vinters, H. V., 2015. Emerging Concepts in Alzheimer’s Disease. Annual Review of Pathology 10:291–319.

333. Querfurth, H. W., and F. M. LaFerla, 2010. Alzheimer’s Disease. The New England Journal of Medicine 362:329–344.

334. Haass, C., and D. J. Selkoe, 2007. Soluble Protein Oligomers in Neurodegeneration: Lessons from the Alzheimer’s Amyloid Beta-Peptide. Nature Reviews. Molecular Cell Biology 8:101–112.

335. Hardy, J., and D. J. Selkoe, 2002. The Amyloid Hypothesis of Alzheimer’s Disease: Progress and Problems on the Road to Therapeutics. Science (New York, N.Y.) 297:353–356.

336. Palop, J. J., and L. Mucke, 2010. Amyloid-Beta-Induced Neuronal Dysfunction in Alzheimer’s Disease: From Synapses toward Neural Networks. Nature Neuroscience 13:812–818.

337. Tu, S., S.-i. Okamoto, S. A. Lipton, and H. Xu, 2014. Oligomeric A β-Induced Synaptic Dysfunction in Alzheimer’s Disease. Molecular Neurodegeneration 9:48.

338. Karran, E., and J. Hardy, 2014. A Critique of the Drug Discovery and Phase 3 Clinical Programs Targeting the Amyloid Hypothesis for Alzheimer Disease. Annals of Neurology 76:185–205.

339. Segal, M., and E. Korkotian, 2014. Endoplasmic Reticulum Calcium Stores in Dendritic Spines. Frontiers in Neuroanatomy 8:64.

340. Popugaeva, E., E. Pchitskaya, and I. Bezprozvanny, 2017. Dysregulation of Neuronal Calcium Homeostasis in Alzheimer’s Disease - A Therapeutic Opportunity? Biochemical and Biophysical Research Communications 483:998–1004.

341. Briggs, C. A., S. Chakroborty, and G. E. Stutzmann, 2017. Emerging Pathways Driving Early Synaptic Pathology in Alzheimer’s Disease. Biochemical and Biophysical Research Communications 483:988–997.

342. Bergmans, B. A., and B. De Strooper, 2010. Gamma-Secretases: From Cell Biology to Therapeutic Strategies. The Lancet. Neurology 9:215–226.

343. Karch, C. M., C. Cruchaga, and A. M. Goate, 2014. Alzheimer’s Disease Genetics: From the Bench to the Clinic. Neuron 83:11–26.

344. Naj, A. C., G. D. Schellenberg, and Alzheimer’s Disease Genetics Consortium (ADGC), 2017. Genomic Variants, Genes, and Pathways of Alzheimer’s Disease: An Overview. American Journal of Medical Genetics. Part B, Neuropsychiatric Genetics: The Official Publication of the International Society of Psychiatric Genetics 174:5–26.

345. Supnet, C., and I. Bezprozvanny, 2011. Presenilins Function in ER Calcium Leak and Alzheimer’s Disease Pathogenesis. Cell Calcium 50:303–309.

346. Tu, H., O. Nelson, A. Bezprozvanny, Z. Wang, S.-F. Lee, Y.-H. Hao, L. Serneels, B. De Strooper, G. Yu, and I. Bezprozvanny, 2006. Presenilins Form ER Ca2+ Leak Channels, a Function Disrupted by Familial Alzheimer’s Disease-Linked Mutations. Cell 126:981–993.

347. Duggan, S. P., and J. V. McCarthy, 2016. Beyond γ-Secretase Activity: The Multifunctional Nature of Presenilins in Cell Signalling Pathways. Cellular Signalling 28:1–11.

348. Ito, E., K. Oka, R. Etcheberrigaray, T. J. Nelson, D. L. McPhie, B. Tofel-Grehl, G. E. Gibson, and D. L. Alkon, 1994. Internal Ca2+ Mobilization Is Altered in Fibroblasts from Patients with Alzheimer Disease. Proceedings of the National Academy of Sciences of the United States of America 91:534–538.

349. Stutzmann, G. E., A. Caccamo, F. M. LaFerla, and I. Parker, 2004. Dysregulated IP3 Signaling in Cortical Neurons of Knock-in Mice Expressing an Alzheimer’s-Linked Mutation in Presenilin1 Results in Exaggerated Ca2+ Signals and Altered Membrane Excitability. The Journal of Neuroscience: The Official Journal of the Society for Neuroscience 24:508–513

350. Chan, S. L., M. Mayne, C. P. Holden, J. D. Geiger, and M. P. Mattson, 2000. Presenilin-1 Mutations Increase Levels of Ryanodine Receptors and Calcium Release in PC12 Cells and Cortical Neurons. The Journal of Biological Chemistry 275:18195–18200.

351. Stutzmann, G. E., I. Smith, A. Caccamo, S. Oddo, F. M. Laferla, and I. Parker, 2006. Enhanced Ryanodine Receptor Recruitment Contributes to Ca2+ Disruptions in Young, Adult, and Aged Alzheimer’s Disease Mice. The Journal of Neuroscience: The Official Journal of the Society for Neuroscience 26:5180–5189

352. Cheung, K.-H., D. Shineman, M. Muller, C. Cardenas, L. Mei, J. Yang, T. Tomita, T. Iwatsubo, V. M.-Y. Lee, and J. K. Foskett, 2008. Mechanism of Ca2+ Disruption in Alzheimer’s Disease by Presenilin Regulation of InsP3 Receptor Channel Gating. Neuron 58:871–883.

353. Bruno, A. M., J. Y. Huang, D. A. Bennett, R. A. Marr, M. L. Hastings, and G. E. Stutzmann, 2012. Altered Ryanodine Receptor Expression in Mild Cognitive Impairment and Alzheimer’s Disease. Neurobiology of Aging 33:1001.e1–6.

354. Green, K. N., A. Demuro, Y. Akbari, B. D. Hitt, I. F. Smith, I. Parker, and F. M. LaFerla, 2008. SERCA Pump Activity Is Physiologically Regulated by Presenilin and Regulates Amyloid Beta Production. The Journal of Cell Biology 181:1107–1116

355. Leissring, M. A., Y. Akbari, C. M. Fanger, M. D. Cahalan, M. P. Mattson, and F. M. LaFerla, 2000. Capacitative Calcium Entry Deficits and Elevated Luminal Calcium Content in Mutant Presenilin-1 Knockin Mice. The Journal of Cell Biology 149:793–798

356. Yoo, A. S., I. Cheng, S. Chung, T. Z. Grenfell, H. Lee, E. Pack-Chung, M. Handler, J. Shen, W. Xia, G. Tesco, A. J. Saunders, K. Ding, M. P. Frosch, R. E. Tanzi, and T. W. Kim, 2000. Presenilin-Mediated Modulation of Capacitative Calcium Entry. Neuron 27:561–572.

357. Sun, S., H. Zhang, J. Liu, E. Popugaeva, N.-J. Xu, S. Feske, C. L. White, and I. Bezprozvanny, 2014. Reduced Synaptic STIM2 Expression and Impaired Store-Operated Calcium Entry Cause Destabilization of Mature Spines in Mutant Presenilin Mice. Neuron 82:79–93.

358. Zhang, H., L. Wu, E. Pchitskaya, O. Zakharova, T. Saito, T. Saido, and I. Bezprozvanny, 2015. Neuronal Store-Operated Calcium Entry and Mushroom Spine Loss in Amyloid Precursor Protein Knock-In Mouse Model of Alzheimer’s Disease. The Journal of Neuroscience: The Official Journal of the Society for Neuroscience 35:13275–13286

359. Lessard, C. B., M. P. Lussier, S. Cayouette, G. Bourque, and G. Boulay, 2005. The Overexpression of Presenilin2 and Alzheimer’s-Disease-Linked Presenilin2 Variants Influences TRPC6-Enhanced Ca2+ Entry into HEK293 Cells. Cellular Signalling 17:437–445.

360. Bae, C. S., and J. Song, 2017. The Role of Glucagon-Like Peptide 1 (GLP1) in Type 3 Diabetes: GLP-1 Controls Insulin Resistance, Neuroinflammation and Neurogenesis in the Brain. International Journal of Molecular Sciences 18

361. Holst, J. J., 2007. The Physiology of Glucagon-like Peptide 1. Physiological Reviews 87:1409–1439.

362. Holst, J. J., R. Burcelin, and E. Nathanson, 2011. Neuroprotective Properties of GLP-1: Theoretical and Practical Applications. Current Medical Research and Opinion 27:547–558.

363. Duarte, A. I., E. Candeias, S. C. Correia, R. X. Santos, C. Carvalho, S. Cardoso, A. Plácido, M. S. Santos, C. R. Oliveira, and P. I. Moreira, 2013. Crosstalk between Diabetes and Brain: Glucagon-like Peptide-1 Mimetics as a Promising Therapy against Neurodegeneration. Biochimica Et Biophysica Acta 1832:527–541.

364. McClean, P. L., and C. Holscher, 2014. Liraglutide Can Reverse Memory Impairment, Synaptic Loss and Reduce Plaque Load in Aged APP/PS1 Mice, a Model of Alzheimer’s Disease. Neuropharmacology 76 Pt A:57–67.

365. Hölscher, C., 2014. Central Effects of GLP-1: New Opportunities for Treatments of Neurodegenerative Diseases. The Journal of Endocrinology 221:T31–41

366. Akter, K., E. A. Lanza, S. A. Martin, N. Myronyuk, M. Rua, and R. B. Raffa, 2011. Diabetes Mellitus and Alzheimer’s Disease: Shared Pathology and Treatment? British Journal of Clinical Pharmacology 71:365–376.

367. Takeda, S., N. Sato, H. Rakugi, and R. Morishita, 2011. Molecular Mechanisms Linking Diabetes Mellitus and Alzheimer Disease: Beta-Amyloid Peptide, Insulin Signaling, and Neuronal Function. Molecular bioSystems 7:1822–1827.

368. Baglietto-Vargas, D., J. Shi, D. M. Yaeger, R. Ager, and F. M. LaFerla, 2016. Diabetes and Alzheimer’s Disease Crosstalk. Neuroscience and Biobehavioral Reviews 64:272–287.

369. Bailey, A., A. Le Couteur, I. Gottesman, P. Bolton, E. Simonoff, E. Yuzda, and M. Rutter, 1995. Autism as a Strongly Genetic Disorder: Evidence from a British Twin Study. Psychological Medicine 25:63–77.

370. Klei, L., S. J. Sanders, M. T. Murtha, V. Hus, J. K. Lowe, A. J. Willsey, D. Moreno-De-Luca, T. W. Yu, E. Fombonne, D. Geschwind, D. E. Grice, D. H. Ledbetter, C. Lord, S. M. Mane, C. L. Martin, D. M. Martin, E. M. Morrow, C. A. Walsh, N. M. Melhem, P. Chaste, J. S. Sutcliffe, M. W. State, E. H. Cook, K. Roeder, and B. Devlin, 2012. Common Genetic Variants, Acting Additively, Are a Major Source of Risk for Autism. Molecular Autism 3:9.

371. De Rubeis, S., and J. D. Buxbaum, 2015. Recent Advances in the Genetics of Autism Spectrum Disorder. Current Neurology and Neuroscience Reports 15:36.

372. Gokoolparsadh, A., G. J. Sutton, A. Charamko, N. F. O. Green, C. J. Pardy, and I. Voineagu, 2016. Searching for Convergent Pathways in Autism Spectrum Disorders: Insights from Human Brain Transcriptome Studies. Cellular and molecular life sciences: CMLS 73:4517–4530

373. Wen, Y., M. J. Alshikho, and M. R. Herbert, 2016. Pathway Network Analyses for Autism Reveal Multisystem Involvement, Major Overlaps with Other Diseases and Convergence upon MAPK and Calcium Signaling. PloS One 11:e0153329.

374. Phillips, M., and L. Pozzo-Miller, 2015. Dendritic Spine Dysgenesis in Autism Related Disorders. Neuroscience Letters 601:30–40.

375. Hoxha, E., P. Lippiello, B. Scelfo, F. Tempia, M. Ghirardi, and M. C. Miniaci, 2017. Maturation, Refinement, and Serotonergic Modulation of Cerebellar Cortical Circuits in Normal Development and in Murine Models of Autism. Neural Plasticity 2017:6595740.

376. Krey, J. F., and R. E. Dolmetsch, 2007. Molecular Mechanisms of Autism: A Possible Role for Ca2+ Signaling. Current Opinion in Neurobiology 17:112–119.

377. Terasaki, M., T. Shemesh, N. Kasthuri, R. W. Klemm, R. Schalek, K. J. Hayworth, A. R. Hand, M. Yankova, G. Huber, J. W. Lichtman, T. A. Rapoport, and M. M. Kozlov, 2013. Stacked Endoplasmic Reticulum Sheets Are Connected by Helicoidal Membrane Motifs. Cell 154:285–296.

378. Carrasco, S., and T. Meyer, 2011. STIM Proteins and the Endoplasmic Reticulum-Plasma Membrane Junctions. Annual Review of Biochemistry 80:973–1000.

379. Stefan, C. J., A. G. Manford, and S. D. Emr, 2013. ER-PM Connections: Sites of Information Transfer and Inter-Organelle Communication. Current Opinion in Cell Biology 25:434–442.

380. Rowland, A. A., and G. K. Voeltz, 2012. Endoplasmic Reticulum-Mitochondria Contacts: Function of the Junction. Nature Reviews. Molecular Cell Biology 13:607–625.

381. Vance, J. E., 2014. MAM (Mitochondria-Associated Membranes) in Mammalian Cells: Lipids and Beyond. Biochimica Et Biophysica Acta 1841:595–609.

382. Manor, U., S. Bartholomew, G. Golani, E. Christenson, M. Kozlov, H. Higgs, J. Spudich, and J. Lippincott-Schwartz, 2015. A Mitochondria-Anchored Isoform of the Actin-Nucleating Spire Protein Regulates Mitochondrial Division. eLife 4

383. Mertins, B., G. Psakis, and L.-O. Essen, 2014. Voltage-Dependent Anion Channels: The Wizard of the Mitochondrial Outer Membrane. Biological Chemistry 395:1435–1442.

384. Camara, A. K. S., Y. Zhou, P.-C. Wen, E. Tajkhorshid, and W.-M. Kwok, 2017. Mitochondrial VDAC1: A Key Gatekeeper as Potential Therapeutic Target. Frontiers in Physiology 8:460.

385. Noskov, S. Y., T. K. Rostovtseva, A. C. Chamberlin, O. Teijido, W. Jiang, and S. M. Bezrukov, 2016. Current State of Theoretical and Experimental Studies of the Voltage-Dependent Anion Channel (VDAC). Biochimica Et Biophysica Acta 1858:1778–1790.

386. Paillusson, S., R. Stoica, P. Gomez-Suaga, D. H. W. Lau, S. Mueller, T. Miller, and C. C. J. Miller, 2016. There’s Something Wrong with My MAM; the ER-Mitochondria Axis and Neurodegenerative Diseases. Trends in Neurosciences 39:146–157.

387. Area-Gomez, E., and E. A. Schon, 2017. On the Pathogenesis of Alzheimer’s Disease: The MAM Hypothesis. FASEB journal: official publication of the Federation of American Societies for Experimental Biology 31:864–867.

388. La Rovere, R. M. L., G. Roest, G. Bultynck, and J. B. Parys, 2016. Intracellular Ca(2+) Signaling and Ca(2+) Microdomains in the Control of Cell Survival, Apoptosis and Autophagy. Cell Calcium 60:74–87.

389. Phillips, M. J., and G. K. Voeltz, 2016. Structure and Function of ER Membrane Contact Sites with Other Organelles. Nature Reviews. Molecular Cell Biology 17:69–82.

390. Hetzer, M. W., 2010. The Nuclear Envelope. Cold Spring Harbor Perspectives in Biology 2:a000539.

391. Mauger, J.-P., 2012. Role of the Nuclear Envelope in Calcium Signalling. Biology of the Cell 104:70–83.

392. Collas, P., E. G. Lund, and A. R. Oldenburg, 2014. Closing the (Nuclear) Envelope on the Genome: How Nuclear Lamins Interact with Promoters and Modulate Gene Expression. BioEssays: News and Reviews in Molecular, Cellular and Developmental Biology 36:75–83.

393. Vagnarelli, P., 2014. Repo-Man at the Intersection of Chromatin Remodelling, DNA Repair, Nuclear Envelope Organization, and Cancer Progression. Advances in Experimental Medicine and Biology 773:401–414.

